# Coexistence of Photosynthetic Marine Microorganisms, Viruses, and Grazers: Toward Integration in Ocean Ecosystem Models

**DOI:** 10.1101/2025.07.11.664470

**Authors:** Paul Frémont, Stephen J. Beckett, David Demory, Eric Carr, Christopher L. Follett, Debbie Lindell, David Talmy, Stephanie Dutkiewicz, Joshua S. Weitz

## Abstract

Photosynthetic microorganisms are responsible for primary production at the base of the marine food web that impacts ocean biogeochemistry and ecological interactions. The growth of these microorganisms is balanced by mortality processes, including top-down losses from zooplankton grazing and infection and lysis by viruses. Multiple types of grazers and viruses often coexist despite apparent competition for the same (or similar) microorganisms. Here, we develop a community model of photosynthetic microorganisms, grazers, and viruses that accounts for molar cell and virion elemental quotas suitable for incorporation into ocean ecosystem models. Our aim is to investigate mechanisms that enable coexistence of a virus and a grazer with a single phytoplankton type. To do so, we evaluate the extent to which coexistence is facilitated by: (i) infected phytoplankton, potentially subject to non-preferential intraguild predation, where grazers feed on virally infected microorganisms; (ii) heterogeneity in susceptibility to infection, where microorganisms vary in their resistance to the virus either intracellularly or extracellularly; and (iii) the inclusion of higher-order mortality terms for the predators. We find evidence for a trade-off between the virus latent period and virulence in facilitating a coexistence regime. The inclusion of an explicit latent period can generate oscillations of all populations that facilitate coexistence by reducing the growth rate of the free virus, or lead to system collapse when oscillations become too large. Heterogeneity in phage susceptibility promotes coexistence through resource partitioning between the predators, while higher-order mortality terms widen the coexistence regime by stabilizing the system. We observe strong sensitivity of model outcomes to viral life history traits, including shifts in infected cell percentages and the balance between virally and zooplankton-induced mortality. Finally, taking advantage of algebraic calculation of model equilibria, we identify life history trait parameter combinations that yield realistic ecological properties in simplified oligotrophic and mesotrophic epipelagic environments. Importantly, our ecological models suggest that ongoing efforts to embed virus dynamics in large-scale ocean ecosystem models should likely include phytoplankton types that are moderately to strongly resistant to viral infection and lysis.

## Introduction

Marine phytoplankton are a diverse group of photosynthetic microorganisms at the base of the ocean food web (Field et al. 1998) that play a crucial role in global biogeochemical cycles (Eppley and Peterson 1979) and are key contributors to the biological carbon pump (Henson et al. 2012; Guidi et al. 2015; Frémont et al. 2022). Phytoplankton are subject to multiple mortality. One such mortality mode is predation/consumption by zooplankton such as heterotrophic nanoflagelattes (Zwirglmaier et al. 2009), dinoflagellates (Sherr and Sherr 1994), and copepods (Dussart 1965). Marine phytoplankton can also be infected by viruses (Suttle 2007), leading to the release of their cellular contents into the environment through cell lysis (Fuhrman and Noble 1995; Wilhelm and Suttle 1999; Weitz 2015). Viruses are present at high densities, typically exceeding 10^10^*/L* in marine surface waters (Wigington et al. 2016) and can be major drivers of phytoplankton mortality (Bergh et al. 1989; Proctor and Fuhrman 1990; Fuhrman and Noble 1995; Suttle 2007; Mojica et al. 2016; Talmy et al. 2019b; Biggs et al. 2021; Carlson et al. 2022; Beckett et al. 2024) which, in turn, can lead to significant impacts on the flow of matter and nutrients (Wilhelm and Suttle 1999; Jover et al. 2014; Weitz et al. 2015; Guidi et al. 2015; Talmy et al. 2019a).

Current large-scale ocean biogeochemistry models include various forms of biological complexity in their representation of biological and inorganic components (Ilyina et al. 2013; Aumont et al. 2015; Séférian et al. 2020; Stock et al. 2020; Negrete-García et al. 2022). Some models incorporate comprehensive size-structured food webs, featuring numerous classes of phytoplankton, diazotrophs, mixotrophs, and several types of grazers (Dutkiewicz et al. 2020; Stock et al. 2020). The increasing complexity of these models has allowed the theoretical exploration of numerous processes on a large scale, such as the emergence of global patterns of phytoplankton diversity (Dutkiewicz et al. 2020), the impact of ocean acidification (Dutkiewicz et al. 2015a), or the importance of iron in the limitation of phytoplankton growth (Tagliabue et al. 2016). However, these large-scale models generally lack an explicit representation of the processes linked to phytoplankton infection by viruses, which has been recognized as an important future challenge (Mateus 2017). The parameterization and implementation of such models remain challenging due to the complexity of virus-host and virus-environment interactions. Critically, this challenge requires both experiments characterizing viral life history traits as well as *in situ* measurements of viral and phytoplankton concentrations to inform the models.

Datasets of experimental measurements of life history traits of phytoplankton viruses are increasingly available (Edwards and Steward 2018; Edwards et al. 2021; Maidanik et al. 2022). Measured traits include burst size (the number of virions produced in a single viral infection), latent period (the time between infection and viral lysis), and adsorption rate (the rate of encounter at which an infection is initiated). In particular, the adsorption rate of viruses can be highly variable and has been identified as crucial in determining viral population dynamics (Talmy et al. 2019b). This parameter depends both on the size of the host and the virus in relation to the size dependence of Brownian motion (Berg and Purcell 1977) and the swimming speed of host microbes (Murray and Jackson 1992; Talmy et al. 2019b; Edwards et al. 2021). For viruses, the adsorption rate can vary by several orders of magnitude and is typically less (sometimes significantly less) than biophysical limits (Talmy et al. 2019b). Efforts are also underway to harmonize the inference of viral life history traits by combining *in vitro* experiments with dynamical models of viral infection (Hinson et al. 2023; Dominguez-Mirazo et al. 2024).

Likewise, *in situ* measurements of virus concentrations and infected cells (*e*.*g*., via the Polony (Baran et al. 2018; Goldin et al. 2020) and iPolony methods (Mruwat et al. 2021; Carlson et al. 2022)) are increasingly common and can be used to compare viral-induced lysis to zooplankton grazing (Baudoux et al. 2007; Mojica et al. 2016; Carlson et al. 2022; Biggs et al. 2021; Beckett et al. 2024). For example, the observed infection percentage of *Prochlorococcus* cells is low in the surface waters of the North Pacific Subtropical gyre (NPSG) (Mruwat et al. 2021; Carlson et al. 2022), with cyanobacteria mortality dominated by zooplankton grazing (Beckett et al. 2024; Connell et al. 2020). However, other *in situ* studies have shown significant latitudinal variation in the relative contribution to mortality by viruses and grazers (Mojica et al. 2016; Carlson et al. 2022). In the North Pacific, a hot spot of infection has been consistently reported over multiple years in the transition zone between subtropical and subpolar gyres (Carlson et al. 2022). In the southern ocean, high levels of cyanobacteria infection have also been reported in surface waters (Gochev et al. 2025). In the North Atlantic, virus-induced mortality of cyanobacteria is an important factor at low and mid latitudes, while mortality associated with zooplankton dominated at higher latitudes (*>* 56^°^N) (Mojica et al. 2016). In eukaryotic phytoplankton, the grazing rates of *Phaeocystis* and picoeukaryotes have been shown to be dominated by viruses or zooplankton, but both rates were rarely found to be elevated simultaneously in the Southern Ocean (Biggs et al. 2021). These data suggest that there are cases where either viruses or zooplankton appear to be the dominant mortality agent, and their relative impact can vary depending on the eukaryotic phytoplankton taxon. For example, virally-induced losses have been found to be more important in cryptophytes, while zooplankton grazing was found to be more important in diatoms (Biggs et al. 2021). Finally, in the northeastern oligotrophic Atlantic, viral-induced mortality was the dominant process for picoeukaryote phytoplankton while zooplankton mortality was the dominant process for cyanobacteria (Baudoux et al. 2007).

Multiple studies have begun to integrate viral lysis into biogeochemical models, exploring their behavior, and investigating various questions related to the ecology and biogeochemical roles of viral infections in the oceans (Fuhrman 1999; Weitz et al. 2015; Talmy et al. 2019a; Talmy et al. 2019b; Biswas et al. 2020; Demory et al. 2020; Demory et al. 2021; Flynn et al. 2022; Beckett et al. 2024). From a biogeochemical perspective, viral lysis is hypothesized to increase organic matter recycling, increase net primary production, and reduce energy transfer to higher trophic levels, through reduced grazing by zooplankton, a set of processes known as the viral “shunt” (Fuhrman 1999; Wilhelm and Suttle 1999; Weitz et al. 2015). In a modeling study of the California Current Ecosystem (Talmy et al. 2019a), it was estimated that a significant portion of the phytoplankton carbon loss from viral lysis, is redirected into nutrient recycling through viral lysates. However, this estimate comes with considerable uncertainty (29 ± 20%) (Talmy et al. 2019a). It also remains unclear to what extent viral lysis could enhance carbon export through the viral “shuttle”, the process by which viral lysis products stimulate aggregation and sinking of particulate organic matter, thereby enhancing carbon export to the deep ocean (Sullivan et al. 2017; Guidi et al. 2015; Zimmerman et al. 2020). *In situ* observations suggest that this is the case through aggregate formation resulting from cell lysis and faster sinking than individual cells (Yamada et al. 2018; Laber et al. 2018), and potentially through increased grazing on viral particles and infected cells (Zimmerman et al. 2020). Including viral lysis in large-scale biogeochemical models would enable exploration of the relative balance of virus versus grazer impacts and “shunt” and “shuttle” mechanisms at global scales. Prior to incorporating viral lysis into large-scale biogeochemical models, it is crucial to first develop and understand these ecological models in simpler settings. Doing so allows for a clearer understanding of virus, grazer, and microbial dynamics.

A prerequisite to integration in large-scale models, viral lysis models have to enable coexistence between the virus, the host and its zooplankton grazers. While a substantial body of literature explores the question of coexistence in ecological systems (Gause 1934; Hardin 1960; Armstrong and McGehee 1980; Chesson 1982; Polis and Holt 1992; Holt and Polis 1997; Huisman and Weissing 1999; Chesson 2000; Hin et al. 2011; Ellner et al. 2019; Yamamichi et al. 2022; Sieben et al. 2022; Orr et al. 2025), relatively few studies have examined how coexistence regimes emerge between viruses and zooplankton that both exploit phytoplankton as a resource (Levin et al. 1977; Weitz et al. 2015; Thingstad and Våge 2019; Biswas et al. 2020; Flynn et al. 2022). In the case of the Lotka-Volterra model (Lotka 1920), the competitive exclusion principle applies to a system with two predators and one prey: one predator excludes the other (Armstrong and McGehee 1980). In resource competition models, increasing the complexity of the model by adding new resources can lead to chaotic behavior, where coexistence can emerge from such dynamics, as demonstrated in the case of the paradox of plankton (Huisman and Weissing 1999). Coexistence equilibria have also been investigated in host-virus systems with the inclusion of an infected class and explicit nutrients (Levin et al. 1977), as well as in multitrophic marine food webs, including zooplankton grazing (Weitz et al. 2015). These studies suggest that fluctuating resource environments, endogenous oscillations, and stabilizing community effects can enable coexistence. Another theoretical study explored the role of selective grazing, where zooplankton avoid infected cells, in shaping the coexistence and dynamic behavior (chaotic or stable) of a model that includes a grazer, a free virus, and an infected class of phytoplankton (Biswas et al. 2020). An adaptive host fitness model that responds to high viral abundance by increasing viral resistance has also been proposed to facilitate the coexistence between viruses, grazers, and phytoplankton (Thingstad and Våge 2019). Finally, the inclusion of parasites, such as viruses, is expected to increase the coexistence within plankton communities and, consequently, the diversity of marine food webs (Thingstad 2000; Dunne et al. 2013; Weitz et al. 2015; Flynn et al. 2022).

In this study, we first develop an elemental quota version of the **S**usceptible cell -**I**nfected cell – **V**irus (*SIV* ) model (Levin et al. 1977; Weitz 2015) which is suitable for large-scale biogeochemical models accounting for cell and virion quotas. Including quotas provides a rationale for conversion between count and elemental concentration frameworks of viral ecology (Figure 1a). We develop extensions to this baseline model, including the addition of a grazing class, in order to explore how increasing ecological complexity (to include biologically relevant mechanisms) affects coexistence regimes for susceptible cells and their grazing and viral predators. We first explore the relevance of including an infected class (*I*) for the coexistence of the predators by comparing simplified models (*SV Z* and *SIV Z*, Figure 1a, b) across the virulence spectrum of the virus. Next, we incorporate mutations from the susceptible type (*S*) to an extracellular or intracellular resistant type in the *SV RZ* and *SIV RZ* models (Figure 1a, b), with varying resistance strengths, and examine the resulting coexistence regimes. Following this, we investigate the impact of including quadratic mortality terms, commonly used in biogeochemical models, for both viral and grazing predators. Finally, we define ecologically relevant concentrations and infection percentages, along with virus-induced mortality rates, that the system is expected to reach for simplified oligotrophic and mesotrophic epipelagic environments and for four types of phytoplankton (Figure 1b). We then compare these values to model output and life history trait models, and discuss the empirical and theoretical relevance of our results to improving understanding of the joint role of viral and grazing predation to ocean ecosystems.

**Figure 1.**
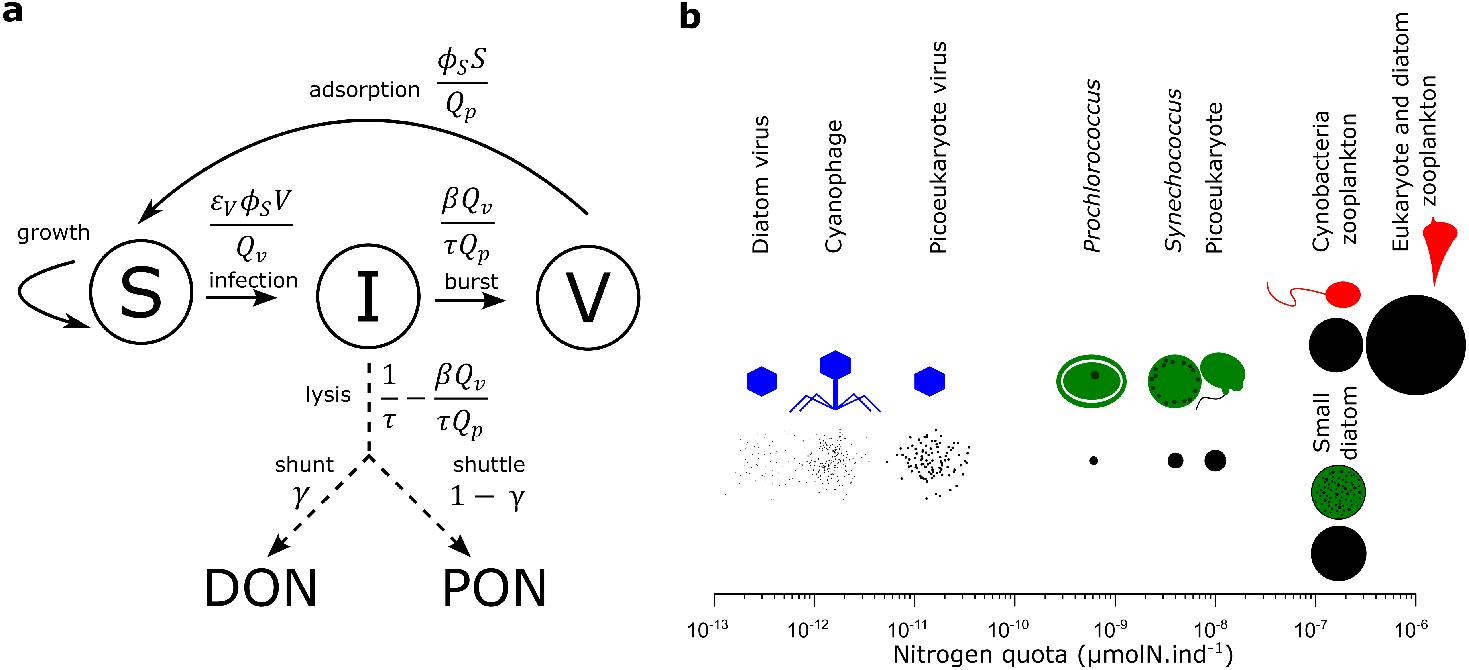
From the count *SIV* model to the elemental concentration *SIV* model. (**a**) Model sketch of the *SIV* elemental concentration model including a shunt and a shuttle term represented respectively as a flux (dashed arrows, not explicitly simulated) to the dissolved organic nitrogen (DON) and the particulate organic nitrogen pool (PON). *DON* and *PON* are not represented as explicit compartments here as we did not model their full dynamics. A fraction *γ* (respectively 1 − *γ*) of the lysate is “shunted” (“shuttled”) to *DON* (to *PON* ). The rates of matter fluxes (in *d*^−1^) are presented for each arrow linked to the inclusion of the viral lysis component (the viral linear decay term is not represented here). (**b**) Nitrogen molar quotas of the different phytoplankton, their respective virus and grazer used in this study. Under the representation of each organism, the radius of each black circle is proportional to the cube root of the nitrogen molar quota, reflecting the scaling of nitrogen with cell volume. Each virus is shown 100 times for better visibility. The cyanobacteria morphology are inpired by *Prochlorococcus* and *Synechococcus*. The picoeukaryote morphology is inspired from *Micromonas pusilla* and the small diatom from *Minidiscus comicus*. Zooplankton are represented by a heterotrophic nanoflagellate for cyanobacteria grazing and by a heterotrophic dinoflagellate for eukaryotic phytoplankton grazing.

## Methods

### From the *SIV* count model to the *SIV* elemental concentration model

We first convert the susceptible-infected-virus (*SIV* ) count model (Levin et al. 1977; Weitz 2015), in units of individuals per volume (individual phytoplankton cell or virion), to an elemental concentration model, typically the currency of biogeochemical models (see Supplementary material for details of the calculation). Here, we arbitrarily chose units of nitrogen molar concentrations. In doing so, we write the viral “shunt” as a flux of the resources from inside the phytoplankton cell to the dissolved organic nitrogen (DON) pool and the viral “shuttle” as a flux to the particulate organic nitrogen (PON) pool but do not include the full microbial loop dynamics. The resulting SIV model for the virus-phytoplankton dynamics (and lysate fluxes to dissolved and particulate organic matter pools) can be written as follows:

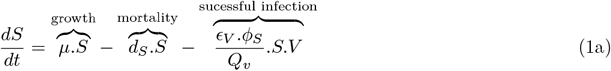

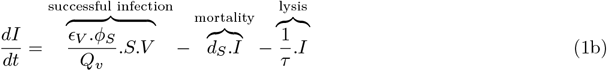

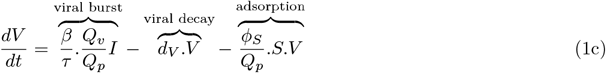

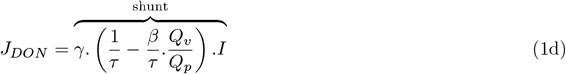

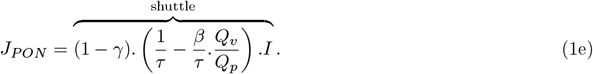

In the above, *µ* is the division rate of susceptible cells (*S*), *d*_*S*_ is the cell mortality rate (natural cell death), *ϕ*_*S*_ is the adsorption rate of viruses to susceptible cells, *ϵ*_*V*_ represents the inverse of intracellular resistance to the virus that is the fraction of adsorption events that lead to a successful infection - and *d*_*V*_ represents the decay rate of infectious virions. Infections have an average duration of *τ*, denoting the latent period, after which a burst size, *β*, of new virions are released into the environment. *Q*_*p*_ and *Q*_*v*_ are the respective nitrogen quota of the phytoplankton and the virus. The fluxes to the dissolved and particulate pools, *J*_*DON*_ and *J*_*PON*_ respectively, are included here for completeness. The microbial loop is not modeled explicitly but could be embedded as part of global ecosystem models (see Discussion for further details). Assuming a fixed stoichiometry, equation 1 also applies to other elements (*e*.*g*., C or P) by substituting the nitrogen quota with the corresponding carbon or phosphorus quota, reflecting the different stoichiometry of viruses (Jover et al. 2014) and phytoplankton (Redfield 1934). We summarize the default set of parameters associated with all models in Table S1-3.

### Elemental concentration models’ equations: *SV Z, SIV Z, SV RZ, SIV RZ*

In this manuscript we explore four elemental concentration models of different complexity based on the above framework: the *SV Z, SIV Z, SV RZ* and *SIV RZ* models. These extensions also include a zooplankton, *Z*, class; and the *SV RZ* and *SIV RZ* models include a resistant, *R*, subpopulation of cells with (partial or full) resistance to infection. The equations of the models are the following (for each model, we emphasize the added terms):

- *SV Z*

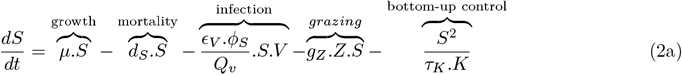

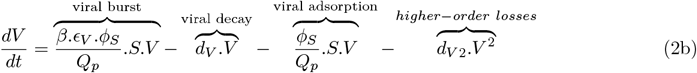

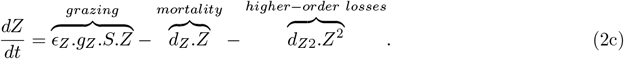
- *SIV Z*

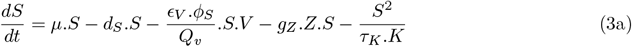

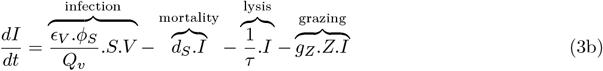

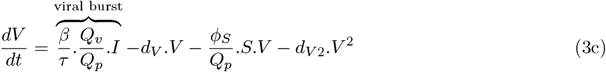

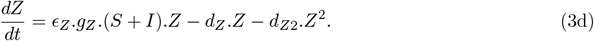
- *SV RZ*

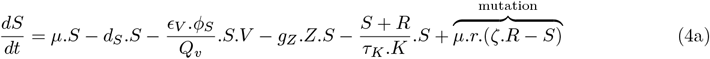

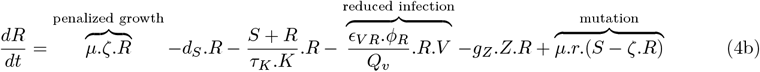

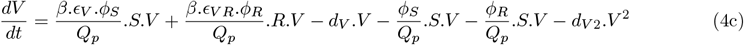

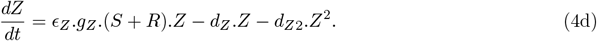
- *SIV RZ dt*

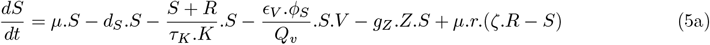

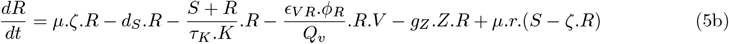

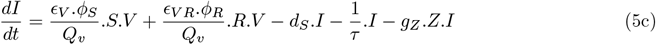

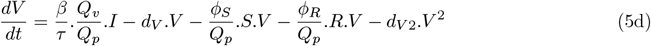

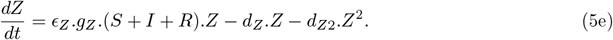

The *SV Z* model, used as a baseline model, corresponds to the simplest predator-prey model, which represents viral infection and lysis as a simple predator (Lotka 1920). In all models, we include the following:

- A carrying capacity term, *K* for the phytoplankton, to approximate bottom-up control by nutrients which has relaxation time *τ*_*K*_ (see later section for an approximation with the resource-consumer model).
- Zooplankton grazing with a clearance rate *g*_*Z*_, gross growth efficiency *ϵ*_*Z*_, and a linear per capita mortality rate of *d*_*Z*_.
- The potential for inclusion of quadratic mortality terms for the virus (*d*_*V* 2_) and the zooplankton (*d*_*Z*2_) (see later sections for rationale).

Each model either includes or does not include the *I* class (*SIV Z* and *SIV RZ* model) and/or a resistant type to the virus (*R* class, *SV RZ* and *SIV RZ* models). Models that include the *I* class introduce an additional parameter *τ*, the latent period of the virus, and a grazing term of the infected class by the zooplankton. The latter term also introduces intraguild predation of the virus (Holt and Polis 1997). Models that include the *R* class introduce a mutation term, *r*, between the *S* and *R* class, extracellular resistance to the virus (through *ϕ*_*R*_) and intracellular resistance to the virus for the *R* type (through *ϵ*_*VR*_). Extracellular resistance is modeled as a reduction in the adsorption rate while intracellular resistance is modeled as a decrease in the probability of an infection that results in the production of virus progeny. Note that to take into account the elemental quota of the zooplankton the grazing terms of the zooplankton (which we call *Z*_*Sgrazing*_ and *Z*_*Igrazing*_ respectively for the *S* and *I* class) could be written as follows:

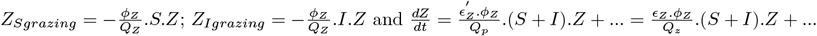

Which gives: 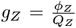 and 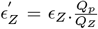. Where *ϕ*_*Z*_ is the zooplankton-phytoplankton encounter rate, *Q*_*z*_ is the zooplankton elemental quota and 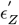 is the grazing efficiency per prey cells consumed.

We summarize the default set of parameters associated with all models in Table S1-3.

### Theoretical equilibria, stability and alternate stable states

For the *SV Z* model and the *SIV Z* model, five possible equilibria are possible:

- *S*^∗^, *I*^∗^, *V* ^∗^, *Z*^∗^
- *S*^∗^, 0, 0, *Z*^∗^
- *S*^∗^, *I*^∗^, *V* ^∗^, 0
- *S*^∗^, 0, 0, 0
- 0, 0, 0, 0 .

For the *SV RZ* and *SIV RZ* model, eleven possible equilibria are possible:

- *S*^∗^, *I*^∗^, *V* ^∗^, *R*^∗^, *Z*^∗^
- 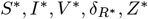
- 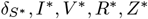
- 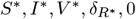
- 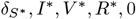
- 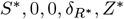
- 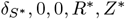
- 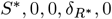
- 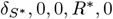
- 0, 0, 0, 0

Where 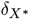 is a state in which the variable *X* is in very low abundance: either *S* or *R* can be in such a state due to the mutation from one to another, while the other is abundant. Note that the five states where *R* is either 0 or 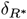 can be mapped to the states of the *SIV Z* and *SV Z* models.

For the purpose of finding ecologically relevant equilibria, we derive the five possible equilibria of the *SIV Z* model (equation 3) and eleven of the *SIV RZ* model (equation 5) in two cases (Supplementary Material):

- *d*_*V* 2_ = 0 and *d*_*Z*2_ *>* 0
- *d*_*V* 2_ *>* 0 and *d*_*Z*2_ *>* 0

We present the equilibria of the models including the *I* class – the equilibria of the other models can be calculated analogously and are included in the full model simulations.

### Size and elemental quotas

In this study, we focus on four different types of phytoplankton: *Prochlorococcus, Synechococcus*, a picoeukaryote (non-diatom) and a small diatom. Each phytoplankton type is considered separately with its own virus and grazer. The elemental concentration model we develop necessitates to fix their sizes to define their elemental quotas, *i*.*e*. the amount of nitrogen in one phytoplankton cell, virion or grazer.

#### Phytoplankton

For eukaryotes, we obtain nitrogen cell quotas following Menden-Deuer and Lessard (2000) and the Redfield ratio (Redfield 1934):

- Diatoms:

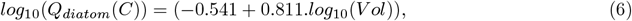
- Eukaryotic protists excluding diatoms:

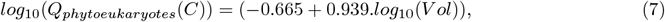

where *V ol* is the cell volume. We then convert to nitrogen units (*µmolN*.*cell*^−1^):

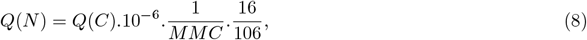

where 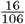 is the Redfield ratio of nitrogen to carbon and *MMC* is the molar mass of carbon.

For cyanobacteria, *Synechococcus* and *Prochlorococcus*, we use the carbon density of *Synechococcus* from Verity et al. (1992):

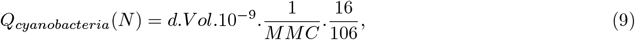

where *d* is the mass density of *Synechococcus*: d=470 fgC.*µ*m^−3^ (Verity et al. 1992) and consistent with other measurements for *Prochlorococcus* (Roth-Rosenberg et al. 2021). For each phytoplankton type, we consider the smallest of each group as modeled in Dutkiewicz et al. (2020) (Table S1).

#### Virus

For each class of phytoplankton, the size of the virus is set as the average size of viruses from the given group from the dataset of Edwards and Steward (2018) (Table S1). Empirical elemental quota relationships for viral capsids are set from Jover et al. (2014):

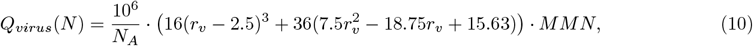

where *r*_*v*_ is the radius of the virion, *N*_*A*_ is the Avogadro number and *MMN* is the molar mass of nitrogen. *Q*_*virus*_(*N* ) is in *µmolN*.*virion*^−1^.

#### Zooplankton

We consider a grazer with a 2.5 *µm* radius for cyanobacteria and 5 *µm* for a picoeukaryote and a diatom. We follow Menden-Deuer and Lessard (2000) for the size-structured elemental concentrations:

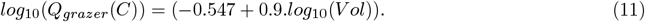

### Life history traits models and parameterization

#### Viral burst size and latent period

We used data from Edwards and Steward (2018) to calibrate models of burst size as a function of different host and virus traits. Burst size (log_10_) was found to be the most highly correlated with host volume (log_10_) and secondarily with the radius of the virus. In addition to these predictors, we add the virus type (ssDNA, dsDNA, ssRNA) and the host type (prokaryote, diatom and eukaryotes) as predictors. Pairs of host type-virus type represented less than 5 times in the dataset are removed from the training set. For eukaryotes, we exclude classes of eukaryotes represented less than four times in the dataset (Pelagophyte, Chlorophyte, Raphidophyte and Cryptophyte), leaving three classes of non diatom eukaryotes: Prasinopyte, Haptophyte and Dinoflagellates. We used leave-one-out cross validation (typically used for small datasets) (Hastie et al. 2009) to assess the performance of 4 types of models: linear models, generalized additive models (GAM) (Wood 2004), random forest (RF) (Breiman and Cutler 2012) and single-layer neural networks (NN) (Venables and Ripley 2002). The cross validation is also used to optimize the hyperparameters of the three latter models: the number of splines for GAM; the number of neurons, learning rate and seed for NN; the number of trees and the number of variables to randomly sample as candidates at each split for RF. The NN model performed the best on average to minimize the root mean square error (RMSE) in leave-one-out cross validation (Figure S1-4). The seed of the NN model for the latent period is not optimized as we empirically find that it overfit the dataset. The optimized NN models are used to set burst sizes and latent period (except for the burst size of diatom viruses where we instead used the GAM model due to apparent overfitting of the NN model (Figure S1-2)). Performances and fitting of the models are summarized for burst size in Figure S1-2 and latent period in Figure S3-4.

#### Viral quadratic mortality

For the higher-order quadratic mortality term of the virus, we use the value of Beckett et al. (2024), that fitted a quadratic mortality for *Prochlorococcus* viruses at station ALOHA (Table S2). The quadratic mortality term, *d*_*V* 2_.*V* ^2^, for the virus represents a generalized nonspecific loss term due, in part, to unspecific binding to other particles present in the ocean, including self collisions and adsorption to particulate organic matter (*POM* ).

#### Viral encounter rate

We use an encounter rate kernel that assumes that the swimming speed of the phytoplankton is considered to have a multiplicative effect on the diffusion kernel (Murray and Jackson 1992; Edwards et al. 2021):

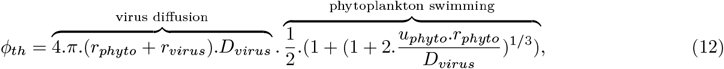

where *r*_*phyto*_ is the radius of the phytoplankton. *u*_*phyto*_ is the swimming speed of the phytoplankton (defined below). *D*_*virus*_ is the diffusivity constant of the virus (defined below), the one of the phytoplankton being negligible in comparison. The diffusivity kernel of the virus is defined as follows (Einstein 1905):

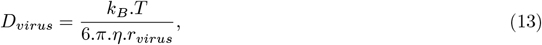

where *k*_*B*_ is the Boltzman constant, T is the temperature in Kelvin, *η* is the dynamic viscosity of water. For the phytoplankton swimming speed we use the following empirical model (Kiørboe 2011; Talmy et al. 2019b) for eukaryotes:

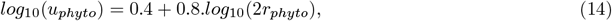

where *r*_*phyto*_ is in *cm* and *u*_*phyto*_ in *cm*.*s*^−1^ (then converted in *m*.*s*^−1^). For diatoms, we considered a constant swimming speed of 10^−6^ *m*.*s*^−1^ (Murase et al. 2011). For cyanobacteria, we consider only diffusion (no swimming speed).

#### Phytoplankton: maximum growth rate and half saturation constant

For the maximum growth rates (in *d*^−1^), we follow the empirical allometric relationship with cell volume from Dutkiewicz et al. (2020):

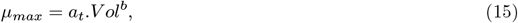

in which *a*_*t*_ depends on the phytoplankton type (diatoms: 3.9, other eukaryotes: 1.4, cyanobacteria (*Prochlorococcus* and *Synechococcus*): 0.8). *b* is equal to -0.08 for diatoms and picoeukaryotes and 0.08 for cyanobacteria. For each of the four types (diatom, picoeukaryote, *Prochlorococcus* and *Synechococcus*), we consider a small cell of each type (Table S1). Each of them is infected by a virus of the average size of viruses from Edwards and Steward (2018) from the respective categories (Table S1). The volume of the phytoplankton and the radius of the virus also allows to set the burst size and latent period of each virus. For the nutrient dependency, *N*_*c*_ the half saturation constant for growth is set as followed:

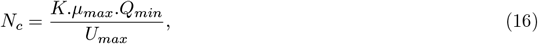

where *K* is the cell nutrient uptake half saturation constant, *Q*_*min*_ is the cell minimum stoichiometric quota, *U*_*max*_ is the maximal cell nutrient uptake rate. For all three parameters, an empirical allometric relationship with the cell volume *V ol* is used (see details in Dutkiewicz et al. 2020; Zakem et al. 2018). Finally, we assume non-preferential grazing by zooplankton with respect to host infection or resistance status.

#### Zooplankton predation

We parameterize zooplankton predation following Dutkiewicz et al. (2020). We consider a zooplankton grazer with a grazing rate of 9.8 (*µmolN*.*L*^−1^)^−1^.*d*^−1^, quadratic mortality of 1.4 (*µmolN*.*L*^−1^)^−1^.*d*^−1^ and linear mortality of 0.067 *d*^−1^ (Dutkiewicz et al. 2020; Dutkiewicz et al. 2015b). Gross growth efficiency is set to 0.3 (Straile 1997). The quadratic mortality term, *d*_*Z*2_.*Z*^2^, represents higher order predation of the zooplankton which is not explicitly represented. The grazer of the cyanobacteria and eukaryotes (picoeukaryote (non-diatom) and small diatom) have the same parameters except their size.

### Simplified epipelagic environments

With the aim to better parameterize viral lysis models for large-scale biogeochemical models, we construct idealized epipelagic environments that account for their carrying capacity, temperature and nutrient limitation of growth and mortality terms and biotic limitation of grazing.

#### Approximation of the resource-consumer model by the carrying capacity model

In biogeochemical models, limiting resources for phytoplankton growth, *e*.*g*. nitrate, are explicitly represented. To simplify our model, we use a carrying capacity term instead. To make an equivalence between the two types of model we set the carrying capacity as the equilibrium concentration of the consumer in the resource-consumer model with a constant nutrient concentration. The resource-consumer model can be written as follow:

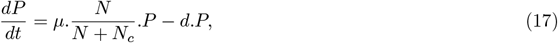

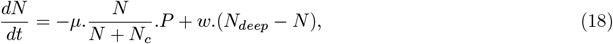

where *P* is the phytoplankton concentration, *N* the nutrient concentration, *N*_*c*_ is the half saturation constant, *w* is the surface-deep mixing rate and *N*_*deep*_ is the deep nutrient concentration, following Weitz et al. (2015). Assuming a constant nutrient concentration, we have:

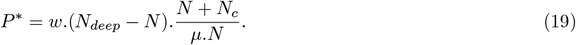

*P* ^∗^ is used as the carrying capacity *K* in the carrying capacity model (Table S3 for parameter values).

#### Growth limitations

We define two typical model epipelagic open ocean environments: oligotrophic and mesotrophic. To represent them in our models, in addition to modifying the carrying capacity, we add constant growth limitations:

- A constant penalty on the growth rate of the phytoplankton to represent changes in nutrient affinity considering a constant nutrient concentration. It is represented with a Michaelis–Menten term:

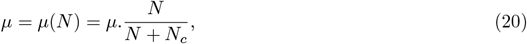

where *N*_*c*_ is the half saturation constant.
- A constant penalty/advantage on the growth rates and all mortality parameters through modulation by temperature following the Eppley curve, parameterized with values from Dutkiewicz et al. (2020):

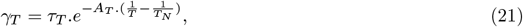

where *T* is the temperature in Kelvin, *τ*_*T*_, *A*_*T*_, *T*_*N*_ are parameters regulating the maximum value and the shape of the limitation curve.

We summarize two sets of values chosen to be representative of the oligotrophic and mesopelagic environments respectively, as well as the effects of these terms in Table S3.

### Coexistence analysis

#### Predator growth rates

To analyze the mechanisms facilitating coexistence in the *SIV Z* model, it is necessary to define the growth rates of the predators (Chesson 1982; Chesson and Ellner 1989; Ellner et al. 2019). The growth rate of the virus is set as the largest real part of the eigenvalues of the Jacobian *J* of the infected cell and virus, *I, V* subsystem, with:

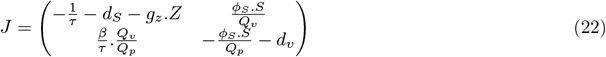

We can calculate the eigenvalues:

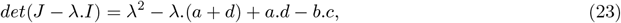

with: 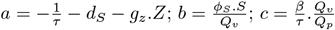 and 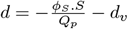.

The eigenvalues of *J* are a solution to the former quadratic equation. The largest eigenvalue is positive if *a*.*d* − *b*.*c <* 0 which is the condition for the virus growth rate to be positive.

The per capita growth rate of the free virus is:

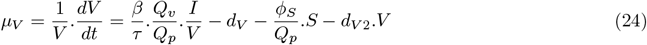

The per capita growth rate of the zooplankton is:

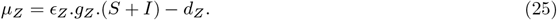

#### Modern coexistence theory

To assess the mechanisms of coexistence of the *SIV Z* model (without quadratic mortality terms of the predators), we perform an invasion analysis following Ellner et al. (2019). Briefly, Modern Coexistence Theory (Ellner et al. 2019) shows that nonlinear responses to fluctuations in resources -either exogenous or endogenous- can facilitate coexistence when species differ in how they respond to those fluctuations. We calculate the differences in growth rates between the invader species and the resident species in the case of both invasion (virus invading and zooplankton invading), and for the different cases of constant, varying, and covarying resources (*I* class and *S* class) (equation 12 from Ellner et al. 2019):

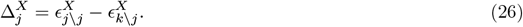

where 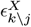 denotes a term (see Ellner et al. 2019 for the complete definition based on the predators’ growth rates) computed for species *k* when species *j* is the invader. *X* corresponds to the different cases of constant, varying, and covarying resources:

- *X* = 0: both resources are constant (average over a periodic cycle from the simulation with species *j* as the invader);
- *X* = *S*: constant S;
- *X* = *I*: constant I;
- *X* = (*S*#*I*): independent variation of I and S;
- *X* = (*SI*): covariance component.

For instance, the term 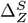 is the *relative nonlinearity* in *S* for the zooplankton, *relative* in the sense that it reflects the difference in how the zooplankton (*Z*) and the virus (*V* ) respond to fluctuations in the prey *S*. The average invasion growth rate of species *j* is the sum of all components:

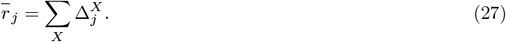

Coexistence is possible in the case of mutual invasibility *i*.*e*. if the invasion growth rate of each species invading the other is positive following Chesson’s criterion (Chesson 1982; Chesson and Ellner 1989). For this analysis we consider the virus as the *I, V* subsystem (equation 22). When fluctuation exists in the resident regime, we perform the analysis over one periodic cycle of the Susceptible type detected using the *find peaks* function from the *SciPy* Python library (Virtanen et al. 2020). In the case of no fluctuation (*e*.*g*. resident Zooplankton), we perform the analysis over the last year of simulation (20 years).

### Growth rate analysis in the coexistence regime

To assess the effect of adding the *I* class on the growth rate of the virus and the zooplankton, we decompose the growth rate of the zooplankton and, in this case, of the free virus (equation 24) in the coexistence regime, with or without fluctuations of their resources. The fluctuation free growth rate of predator *P* (either *V* or *Z*), *i*.*e*. when *I* and *S* concentrations are considered constant, is defined as follows:

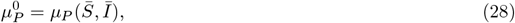

where 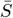 and *Ī* are respectively the average concentration of *S* and *I* (over a periodic cycle in our case). The nonlinearity in *S* (respectively, in *I*) is quantified as the difference between the average growth rate under fluctuating *S* (or *I*) and the growth rate at the average (*i*.*e*. fluctuation-free) values of *S* and *I*:

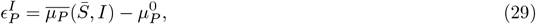

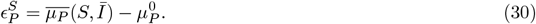

where 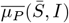 and 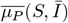 are the average growth rate of *P* when *S* (respectively *I*) is constant and *I* (respectively *S*) is fluctuating over one cycle.

### Fourier analysis

To assess whether the system is stable or oscillatory outside of transient dynamics, we performed a Fourier analysis of the virus raw time series in the last year of simulation (of 20 years). The fft function from the *NumPy* Python library (Harris et al. 2020) is used and the frequency with the largest modulus is extracted. We consider the system to be oscillatory when the largest modulus is greater than 1.

### Ecological metrics

We define the percentage of infected cells as follows:

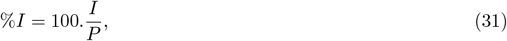

where *P* = *S* + *I* + *R* is the total phytoplankton concentration. The percentage of resistant cells (without considering infected cells) is:

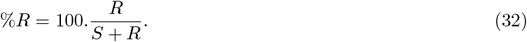

We define the per-capita virus-induced mortality, *m*_*V*_, and zooplankton-induced mortality, *m*_*Z*_, as follows:

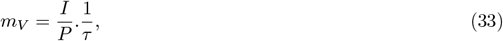

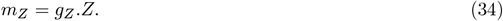

From these metrics, we define the percentage of virus-induced mortality:

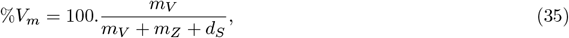

where *d*_*S*_ is the cell mortality rate (natural cell death).

The net primary productivity at equilibrium is defined as the difference between photosynthesis and mortality of the phytoplankton (with contributions from susceptible and resistant cell classes):

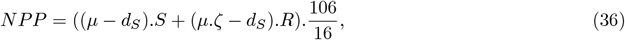

where the C:N Redfield ratio 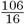 is used to convert into units of *µmolC*.*d*^−1^.*L*^−1^.

### Model testing

#### Target concentrations

We define target concentrations based on the values of virus and phytoplankton concentration measured in the field, generally measured in *individual*.*L*^−1^ (noted *ind*.*L*^−1^ for any tracer throughout the text) (Mojica et al. 2016; Leblanc et al. 2018; Carlson et al. 2022; Beckett et al. 2024) (including these targets in Table S4). For *Prochlorococcus*, the target concentration is defined as 1.5.10^8^ *ind*.*L*^−1^ in an oligotrophic environment and 2.10^8^ *ind*.*L*^−1^ in a model mesotrophic environment, while the target concentration of phages are, respectively, 5.10^8^ and 10^9^ *ind*.*L*^−1^ (Mojica et al. 2016; Carlson et al. 2022). For picoeukaryotes (based on picoeukaryote from Mojica et al. 2016) and a small diatom (based on concentrations of small diatoms from Leblanc et al. 2018 and similar to concentration of nanoeukaryotes in Mojica et al. 2016), in a model mesotrophic environment, target concentrations are respectively 2.10^7^ and 2.10^6^ *ind*.*L*^−1^ (Mojica et al. 2016; Leblanc et al. 2018) and 2.10^7^ and 10^9^ *ind*.*L*^−1^ for their respective viruses. We set the same target concentration for the diatom virus as the concentration for the *Prochlorococcus* phage as the diatom virus is of a similar size and diatom viruses can reach very high concentrations in culture experiments (Tomaru et al. 2021; Arsenieff et al. 2019). Finally, measured values of 1 to 2.10^5^ *ind*.*L*^−1^ for zooplankton are used (Schartau et al. 2010). We consider heterotrophic flagellates with a radius of 2.5 *µm* for cyanobacteria and a radius of 5 *µm* for the picoeukaryote and the small diatom (Schartau et al. 2010; Sherr and Sherr 1994). Note that our model does not include competition which occurs in the ocean. Therefore our target concentration reflect approximate expected values without competition. To further constrain the model outputs, we impose biologically realistic ranges for the percentage of infected cells (0.5% to 10% in the model mesotrophic environment and 0 to 5% in the model oligotrophic environment; Carlson et al. (2022)) and the proportion of virus-induced mortality (0% to 50%). For diatoms, the upper bound on the percentage of infected cells is relaxed due to limited empirical constraints. These bounds are not included in the target concentration error metric (see below), but they must be satisfied during the optimization procedure (see below).

#### Model error

To assess the error between target concentrations and modeled concentrations for an environment *e*, we define the total absolute error, in %, as the sum of absolute relative errors across tracers:

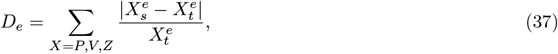

where *X*_*s*_ and *X*_*t*_ are respectively the theoretical and targeted concentrations of a tracer (*P, V* or *Z*). We define the average absolute error across environments (*O* oligotrophic, *M* mesotrophic) as a weighted mean across environments:

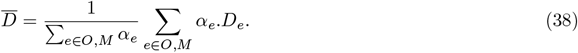

For the picoeukaryote and the small diatom, a weight (parameter *α*_*e*_) of 2 (*i*.*e*. it is counted twice) is given to the error in the model mesotrophic environment as there exist no precise measurements of these organisms in oligotrophic environments (dominated by cyanobacteria) and target concentrations are more arbitrary.

#### Parameter optimization

To minimize the error between theoretical model steady states and target concentrations we perform a grid search of the parameter space using theoretical equilibria of the *SIV Z* model (Supplementary Information). The following parameters are searched: intracellular and extracellular resistance, the resistance cost, the linear and quadratic mortality of the virus (that can be null), and the quadratic mortality of the zooplankton. Explored ranges are summarized in Table 1. We define the extracellular resistance as the ratio between the theoretical encounter rate and the fitted adsorption rate 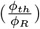 while the intracellular resistance is defined as the inverse of the probability of infection 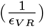 .In our approach, the constraints on the percentage of infected cells and the percentage of virally-induced mortality must be satisfied for an equilibrium to be valid.

**Table 1.** Parameter optimization to target concentrations for an oligotrophic and a mesotrophic simplified epipelagic environment using the approximate theoretical equilibria for the four phytoplankton types. The best parameter value is defined as the one minimizing the average absolute total error across the two environments for a given phytoplankton. The parameter range reflects the minimum and maximum value of a parameter within the first 200 best-fitting models. Parameter ranges tested are indicated below each parameter. Adsorption rate is in *L*.*d*^−1^, extracellular, intracellular resistance and resistance cost are unitless, linear mortality is in *d*^−1^ and quadratic mortality terms are in (*µmolN*.*L*^−1^)^−1^.*d*^−1^. Parameters that are kept fixed are described in Table S1-3.

| Variable |  | *Prochlorococcus* | *Synechococcus* | Picoeukaryote | Diatom |
| --- | --- | --- | --- | --- | --- |
| Adsorption rate<br>$\phi_R$<br>$10^{-12} - 10^{-8}$ | Best | $1.7..10^{-10}$ | $2.15.10^{-10}$ | $1.7.10^{-9}$ | $1.3.10^{-10}$ |
| | Range | $1.6.10^{-11} - 7.7.10^{-10}$ | $7.7.10^{-11} - 1.7.10^{-9}$ | $7.7.10^{-10} - 1.7.10^{-9}$ | $7.7.10^{-11} - 10^{-8}$ |
| Extracellular<br>resistance $\frac{\phi_{th}}{\phi_R}$ | Best | 12.9 | 17.6 | 3.3 | 291.9 |
|  | Range | 3.6 – 129 | 2.3 – 48.8 | 3.3 – 7 | 3.8 – 487 |
| Intracellular<br>resistance $\frac{1}{\epsilon_{VR}}$<br>$1 - 10^4$ | Best | 3.6 | 1.7 | 1.7 | 1 |
|  | Range | 1 – 7.7 | 1 – 12.9 | 1 – 21.5 | 1 – 60 |
| Total<br>resistance<br>$\frac{\phi_{th}}{\phi_R} \cdot \frac{1}{\epsilon_{VR}}$ | Best | 46 | 29 | 5 | 292 |
|  | Range | 10 – 129 | 13 – 63 | 3 – 70 | 135 – 487 |
| Resistance<br>cost $1 - \zeta$<br>$0 - 0.5$ | Best | 0.05 | 0 | 0 | 0.4 |
|  | Range | 0 – 0.5 | 0 – 0.3 | 0 – 0.2 | 0.2 – 0.5 |
| Linear $V$<br>mortality $d_V$<br>$0.01 - 1$ | Best | 0.01 | 0.01 | 0.01 | 0.01 |
|  | Range | 0.01 – 0.1 | 0.01 – 0.05 | 0.01 – 0.5 | 0.01 |
| Quadratic $V$<br>mortality $d_{V2}$<br>$0 - 2000$ | Best | 70 | 90 | 1200 | 100 |
|  | Range | 20 – 500 | 30 – 200 | 80 – 2000 | 70 – 400 |
| Quadratic $Z$<br>mortality $d_{Z2}$<br>$2 - 30$ | Best | 12 | 12 | 22 | 14 |
|  | Range | 10 – 14 | 10 – 18 | 12 – 30 | 10 – 30 |

### Simulations and criterion of existence

Each model is simulated for 20 years using an integration time step of 30 minutes following the Runge-Kutta 4 integration scheme (Lotkin 1951) in a spatially implicit setup. Simulations are run for a range of parameters values for each model, where we choose to provide particular focus on variations in the adsorption rate. For the *SIV Z* and *SIV RZ* models, which include an infected cell class, we also include assessment with respect to varying the latent period. The burst size is fixed by the life history trait model (see Life history traits and parameterization section). For the *SIV RZ* model we also explore the adsorption rate and the resistance strength space while the latent period and burst size are fixed by the life history traits models. To operationally assess the final state of the model, we check whether the concentration of each tracer, *i*.*e* each model compartment, was found to be greater than 1 *ind*.*L*^−1^ in the last year of simulation.

## Results

### Mechanisms of coexistence between the virus and the zooplankton

#### The inclusion of the infected class or the resistant type facilitates regimes of coexistence

We begin by assessing how choice of model structural assumptions promotes coexistence between a virus and a zooplankton population both exploiting the same *Prochlorococcus* population (used as a model organism). We compare simulations of the baseline competition model (*SV Z*, Figure 2a) with those of the more complex *SIV Z* and *SV RZ* models (Figure 2b-c), that incorporate, respectively, an infected host class *I* and a resistant host type *R*. To evaluate long-term population dynamics, beyond initial transient dynamics dependent on initial conditions, we run simulations of each model for 20 years for different viral adsorption rates (*ϕ*_*S*_ = 10^−11^, 10^−10^, 10^−9^, 10^−8^ *L*.*d*^−1^, Figure 3 for 5 first years time series). For the *SV Z* model of *Prochlorococcus* (Figure 2a; equation 2), which resembles a Lotka-Volterra system with one prey and two predators (Lotka 1920), the virus is excluded for *ϕ*_*S*_ = 10^−11^, 10^−10^ *L*.*d*^−1^ (Figure 3a,b) while the zooplankton is excluded for *ϕ*_*S*_ = 10^−9^, 10^−8^ *L*.*d*^−1^ (Figure 3c,d). This aligns with the competitive exclusion principle (Armstrong and McGehee 1980), which states that two species competing for the same resource with linear growth rates cannot coexist indefinitely.

**Figure 2.**
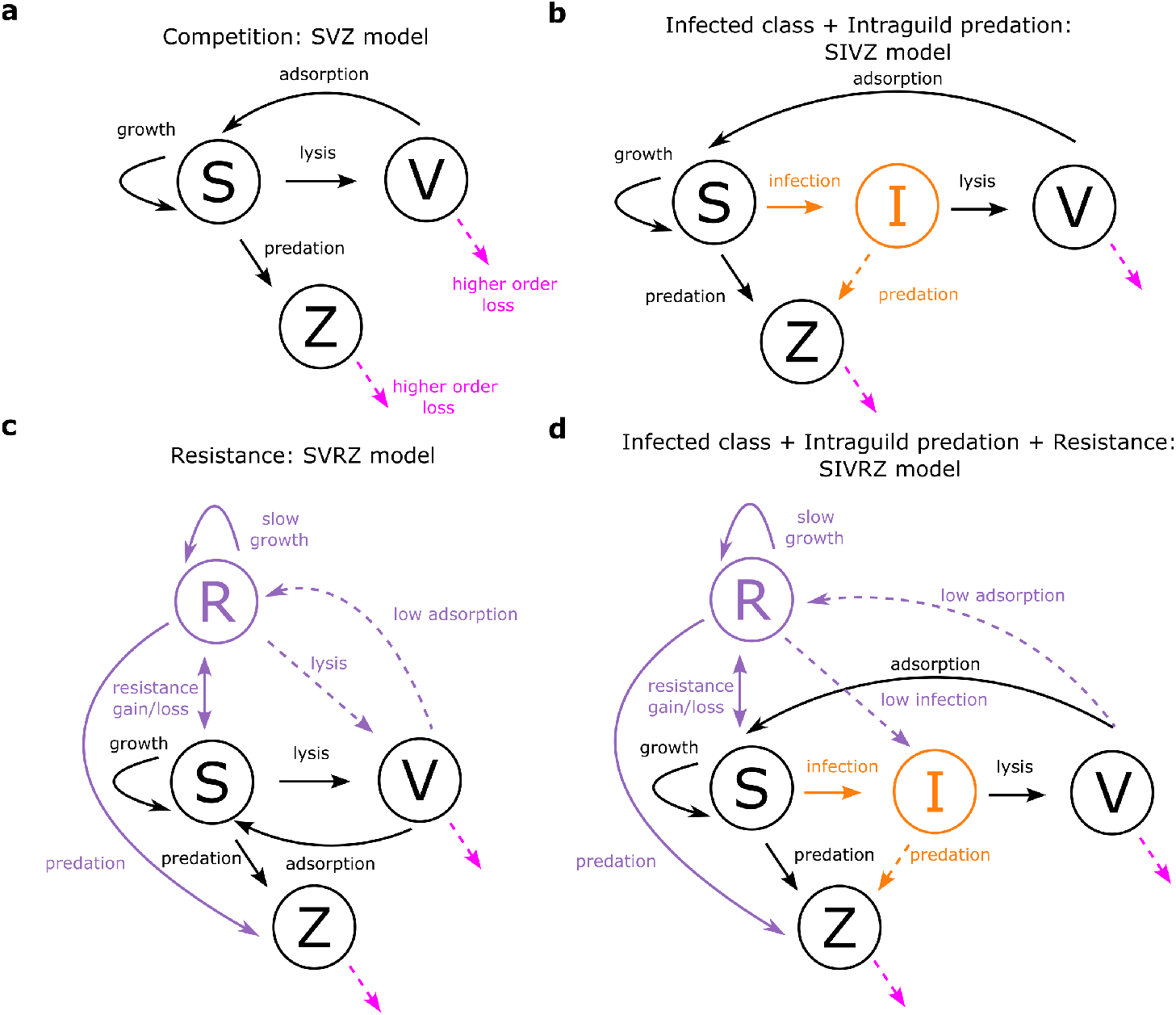
Model sketches illustrating model complexification to facilitate coexistence regimes between the virus and the zooplankton. Sketches of the different viral infection models used in the study: (**a**) *SV Z* model. (**b**) *SV RZ* model. (**c**) *SIV Z* model. (**d**) *SIV RZ* model . S: Susceptible, I: Infected, V: Virus, Z: Zooplankton, R: Resistant type. Arrows represent fluxes of matter. Dashed arrows represent optional terms including higher order losses (quadratic mortality). Orange and purple color respectively indicate fluxes linked to the addition of the *I* and *R* type.

**Figure 3.**
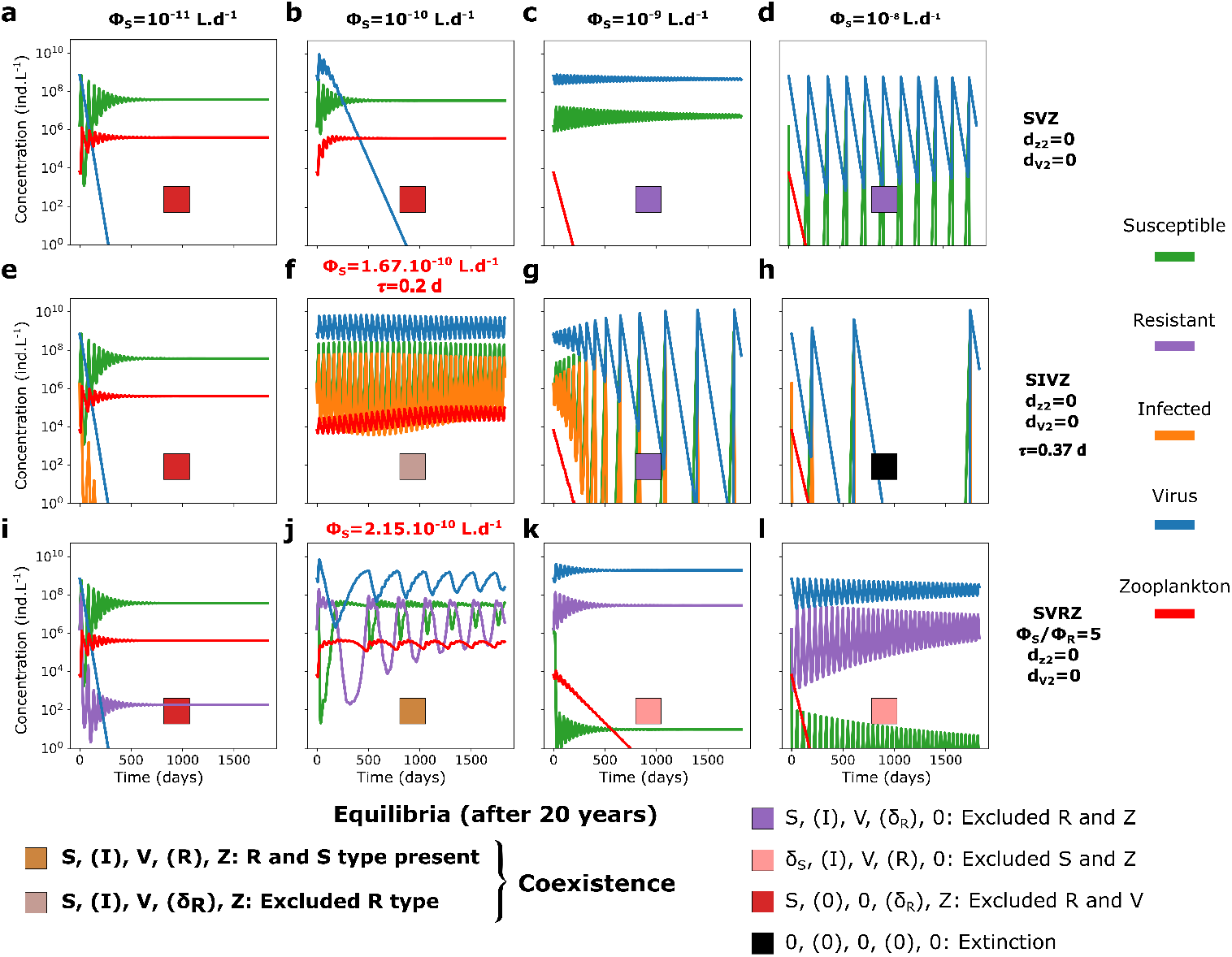
Five year time series of the *SV Z, SIV Z* and the *SV RZ* models for *Prochlorococcus* for four different adsorption rates of the virus without quadratic mortality terms. Adsorption rate from *ϕ*_*S*_ = 10^−11^ *L*.*d*^−1^ to *ϕ*_*S*_ = 10^−8^ *L*.*d*^−1^ (*ϵ*_*V*_ = 1) for (**a**-**d**) the *SV Z* model, (**e**-**h**) the *SIV Z* model and (**i**-**l**) the *SV RZ* model. The tracers’ molar concentrations were converted to count concentrations using the cellular and virion quotas (see Figure S5 for time series in molar concentrations). A latent period of 0.37 *days* and a burst size of 15 were used as parameterized by the life history trait model (see Methods). To showscase the coexistence regime facilitated by the *I* class (Infected) and *R* (Resistant), we slightly changed the parameters (**g**) *τ* = 2.15 *d* and (**j**) *ϕ*_*S*_ = 1.67.10^−10^ *L*.*d*^−1^ (highlighted in red).

To investigate model complexification that may facilitate coexistence between the two predators, we first introduce an infected class of phytoplankton (*I*): the *SIV Z* model (Figure 1b; equation 3). Introducing the infected class to the model adds a new parameter: the latent period of infection that we fix to 0.37 *d* using the life history trait model (Methods, Figure S3 and Table S1). Note, this is representative of measured *Prochlorococcus* cyanophage infections from Mruwat et al. (2021). For *ϕ*_*S*_ = 10^−11^ *L*.*d*^−1^, the virus is still excluded (Figure 3e). However, we find that by shortening the latent period (to 0.2 *d*) and decreasing the adsorption rate (to 1.167 × 10^−10^ *L*.*d*^−1^), an oscillatory coexistence regime results (Figure 3f). When further increasing the adsorption rate of the virus, the zooplankton is either excluded (Figure 3g) or the system collapses, *i*.*e*. all tracers are excluded (Figure 3h). Notably, the latent period generates larger oscillations in the system, and this can lead to system collapse, which was not the case for the *SV Z* model.

An additional model complexification that may facilitate the coexistence of the virus and the zooplankton is the introduction of a resistant type (*R*) with an associated cost to its growth rate. This is represented by the *SV RZ* model (Figure 2c, equation 4). We specifically consider extracellular resistance to viral infection that prevent adsorption, thus the resistance is modeled by a reduced viral adsorption rate, denoted by the parameter *ϕ*_*R*_ (equations 4b and 4c). We arbitrarily set a 5-fold ratio between the adsorption rate of the susceptible cell population and the one of the population of resistant cells 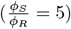. A slight increase in the adsorption rate of the virus from 10^−10^ to 2.15.10^−10^ *L*.*d*^−1^, results in a coexistence regime between the virus and the zooplankton with oscillatory behavior of all classes (Figure 3j). For *ϕ*_*S*_ = 10^−9^, 10^−8^ *L*.*d*^−1^, the zooplankton is excluded. In this regime, the resistant phytoplankton type largely dominates in abundance relative to the susceptible. The latter is only sustained in very low abundance via mutation (Figure 3k,l).

#### A trade-off between latent period and virulence facilitates coexistence

To analyze coexistence regimes beyond single parameter dynamics, we compute coexistence regimes across the virulence spectrum of the virus. For simplicity, we refer to the viral adsorption rate (*ϕ*_*S*_) multiplied by the probability of infection resulting in the production of new virions (*ϵ*_*V*_ ) as its virulence throughout the remainder of the text. Operationally, we use the terms very, moderate, or mildly virulent to denote effective adsorption rates of *ϵ*_*V*_ .*ϕ*_*S*_ *>* 10^−9^, 10^−10^ *< ϵ*_*V*_ *ϕ*_*S*_ *<* 10^−9^, and *ϵ*_*V*_ .*ϕ*_*S*_ *<* 10^−10^ *L*.*d*^−1^ respectively (with *ϵ*_*V*_ = 1, the probability of successful infection). For the *SV Z* model of *Prochlorococcus*, as expected from previously described time series, coexistence occurs between the phytoplankton and either the virus or the zooplankton for all adsorption rates tested, but not both simultaneously. The outcome depends on the viral adsorption rate, with one predator excluding the other (Figure 4a).

**Figure 4.**
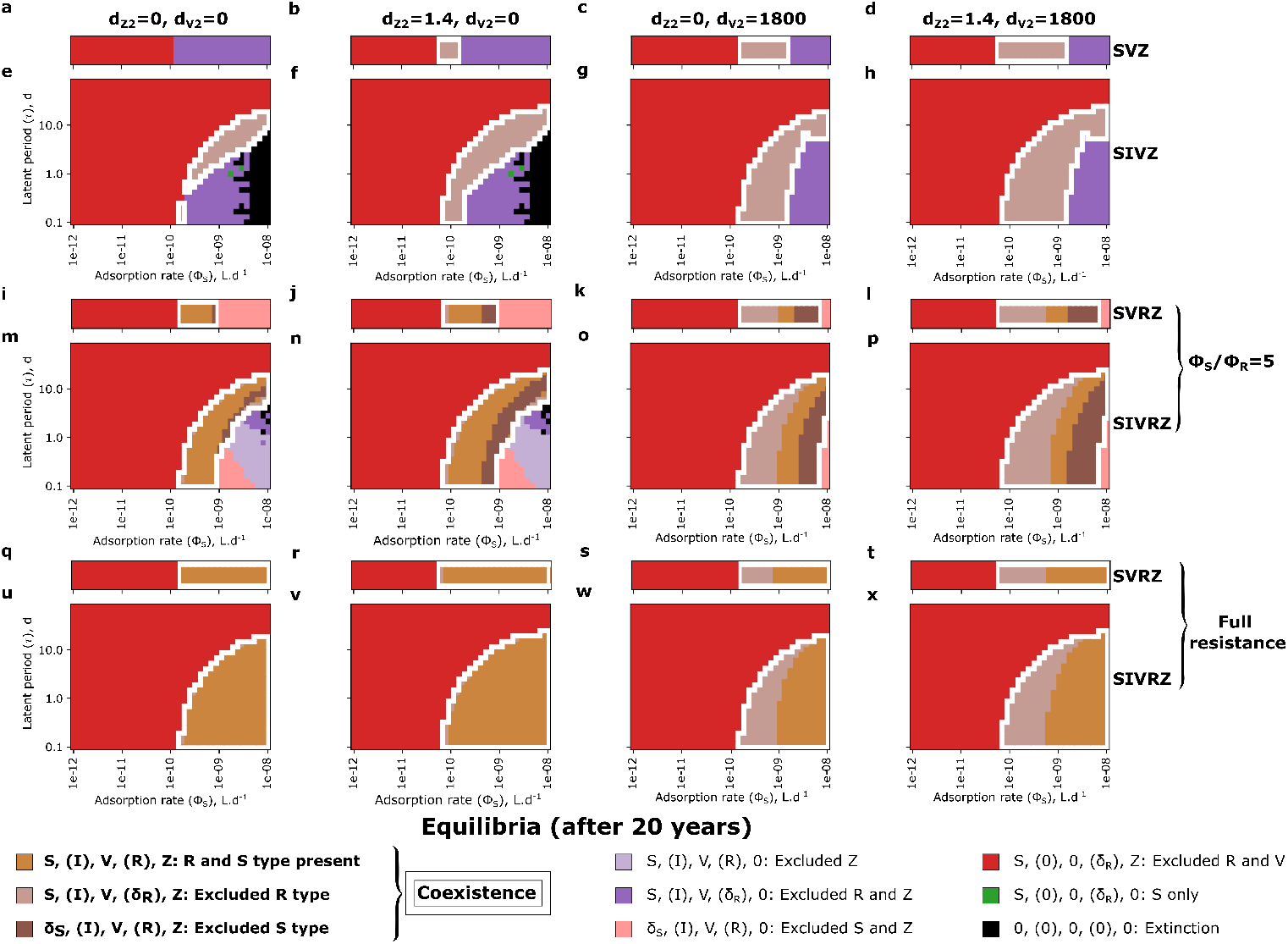
Approximate simulated equilibrium regimes of a virus and a zooplankton modeled as predators of a *Prochlorococcus* for different types of viral lysis models across the adsorption rate and the latent period parameter space. (**a**-**d**) *SV Z* model. (**e**-**h**) *SIV Z* model. (**i**-**l**) Extracellular *SV RZ* model with a partially resistant type: 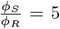. (**m**-**p**) Extracellular *SIV RZ* model with a partially resistant type: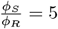. (**q**-**t**) Extracellular *SV RZ* model with a fully resistant type. (**u**-**x**) Extracellular *SIV RZ* model with afully resistant type. From left to right panels: quadratic mortality terms respectively absent, present for the zooplankton only, present for the virus only, and present for both the zooplankton and the virus. The quadratic mortality terms are in (*µmolN*.*L*^−1^).*d*^−1^. The *SV Z* model was run for 25 years to allow full exclusion while the other models were run for 20 years. In the latter case, the equilibrium might not be reached yet. In addition some numerical instabilities might explain the fuzzy transitions between regions with different regimes. To define the equilibrium reached in each simulation we test whether each tracer had a concentration superior to 1 *ind*.*L*^−1^ in the last year of simulation. For each panel, the white contour denotes the coexistence regime.

For the *SIV Z* model, we explore a parameter range for the latent period beyond measured maximum latent periods (maximum of 1.75 days in Edwards and Steward 2018) and most likely beyond what is expected in the ocean, to assess the effect of this parameter across a large range and visualize transitions in model outputs. We find that the inclusion of the *I* class alone, regardless of whether it is grazed or not can facilitate coexistence between the virus and zooplankton. Virus-grazer coexistence is observed only when the carrying capacity term for the phytoplankton is retained and occurs within a narrow range of the viral latent period and adsorption rate (Figure 4e). Only moderate to very virulent viruses are able to coexist with the zooplankton in the region of the virulence spectrum where the virus excludes the grazer in the *SV Z* model (Figure 4a versus Figure 4e). As the virulence increases, the latent period required for the virus to persist also increases, reaching up to approximately 10 days for the most virulent viruses (much longer than the doubling time of the host, Figure 4e). These coexistence regimes are found to be unstable, generating limit cycles that are away from the equilibrium points of the system (Figure S6). This suggests a trade-off between the viral adsorption rate and latent period to facilitate coexistence, where sufficiently small oscillations are necessary to prevent the system from collapsing. The inclusion of the *I* class appears to have two opposing effects. First, in the case of coexistence, the time delay between the viral adsorption to the host and the burst of new viruses generates a limit cycle, with a short time shift between the peaks (highs and lows) of the virus and zooplankton population dynamics (Figure S6). These peaks may either follow one another in the same order (Figure S6a) or not (Figure S6b). Second, for very virulent viruses with a relatively short latent period, the time delay causes strong oscillations in the *SIV* dynamics (within the *SIV Z* model), which can drive the system to extinction (Figure 4e), a behavior not seen in the *SV Z* model (Figure 4a). Interestingly, the inclusion of the *I* class can either support coexistence or lead to extinction, depending on the underlying ecological life-history traits.

#### Modern Coexistence Theory of the *SIV Z* model: non linear oscillations of the susceptible type facilitates coexistence

To further investigate how the inclusion of the infected class (*I*) facilitates the coexistence regime, we performed an invasibility analysis following Chesson’s mutual invasibility criterion of coexistence (Chesson 1982; Chesson and Ellner 1989). When the resident state exhibits fluctuations, invasibility analysis is performed over a single periodic cycle of *S* (Methods). Across 10 replicates of the invasibility analysis, we find good agreement between mutual invasibility and the coexistence regime (Figure 5a, b). However, the region of coexistence extends slightly beyond the parameter range where Chesson’s criterion holds. For highly virulent viruses with long latent periods, the *SIV* model collapses, while the *SIV Z* model still supports coexistence, suggesting a stabilizing effect of the zooplankton in this case. The sensitivity of Chesson’s criterion was 0.76 when requiring all invasibility analysis replicates to be valid and 0.91 when requiring at least one replicate to be valid; these values decreased to 0.71 and 0.85, respectively, when including the *SIV* collapse region as false negatives. It is to be noted that some dynamics within the coexistence regime are still on a slow trajectory toward exclusion, as simulations over 20 years might not fully capture long-term transients. Visualizing the invasion growth rate of the virus (Figure 5b) and of the zooplankton (Figure 5c) shows that the coexistence regime exactly matches the upper border of positive virus invasion growth rate while it approximately corresponds to the lower border of positive invasion growth rate of the zooplankton (likely due to numerical instabilities in the invasion analysis). Following Modern Coexistence Theory (MCT, Ellner et al. 2019), we further decompose these invasion growth rates into their relative components (equation 26). First, when the virus invades the *SZ* system, the fluctuation-free and total invasion growth rates coincide (Figure 5b, d), as *SZ* dynamics always converge to a stable node equilibrium (no fluctuations). In contrast, when the zooplankton invades the resident *SIV* system, the fluctuation free relative invasion growth rate of the zooplankton is negative in the coexistence regime (Figure 5e). Nevertheless, this is compensated by a positive relative nonlinearity in *S* (Figure 5f): the zooplankton takes better advantages of the fluctuations of *S* than does the virus. This suggests that this nonlinearity acts as the main mechanism of coexistence between the virus and the zooplankton. Importantly, this nonlinearity alone is not sufficient for coexistence, as it is positive over a broad region of the parameter space. For coexistence to be enabled, in addition to a positive invasion growth rate of the virus, the positive relative nonlinearity in *S* must be strong enough to offset the negative fluctuation-free invasion growth rate when the zooplankton invades.

**Figure 5.**
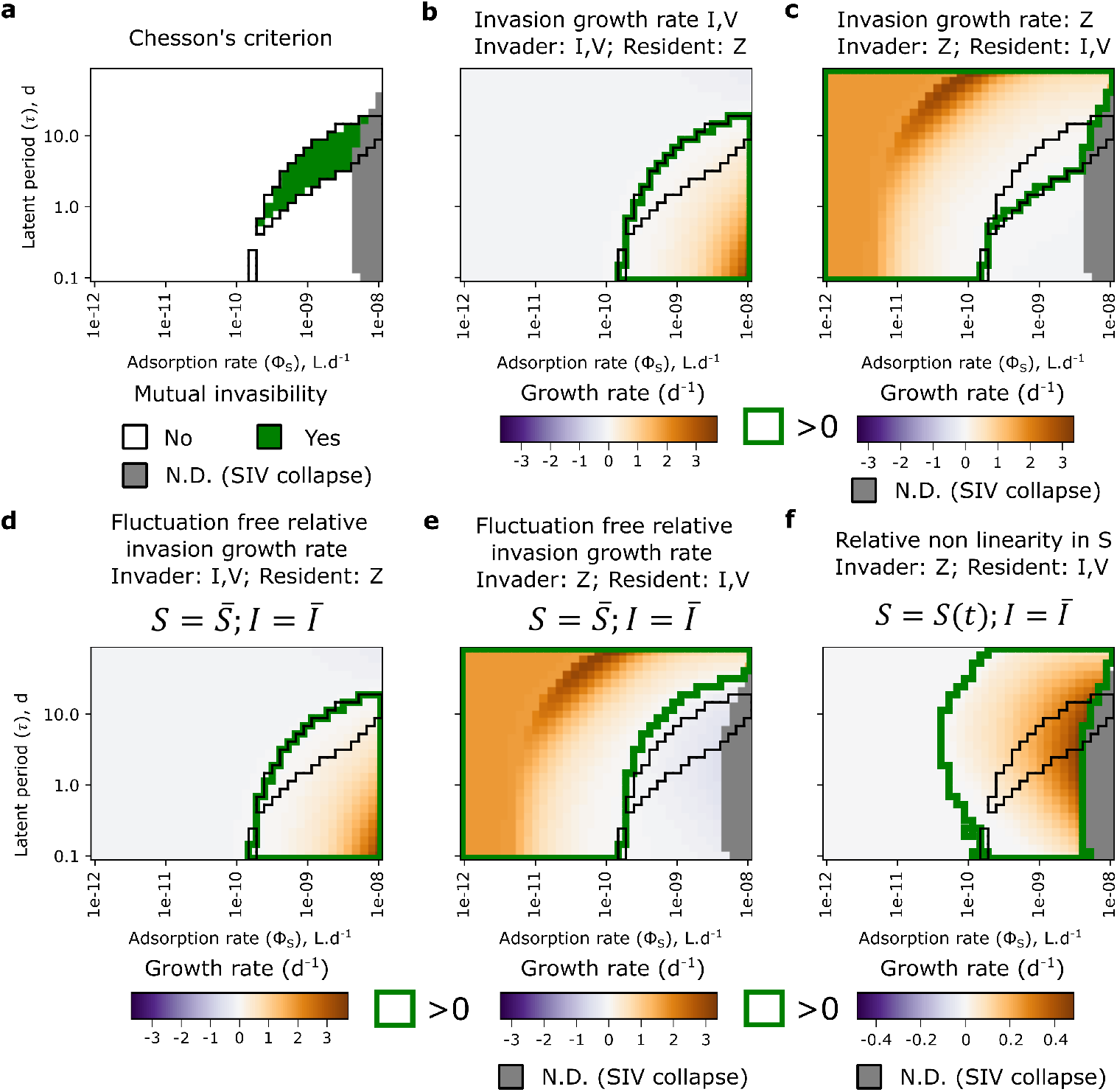
Modern coexistence theory (MCT) analysis of the mechanisms facilitating coexistence of the zooplankton and the virus in the *SIV Z* model without quadratic mortality of the predators. (**a**) Chesson’s criterion of mutual invasibility across 10 replicates of the Modern Coexistence theory analysis: in the green area, mutual invasion (*i*.*e*.) both the invasion growth rate of the virus and the zooplankton are positive for all replicates of the MCT analysis. In the grey area, the analysis is not possible as the *SIV* model collapses in this region. (**b**) Invasion growth rate of the virus (considered as the subsystem IV). (**c**) Invasion growth rate of the zooplankton. (**d**) Fluctuation free relative growth rate of the virus when the virus is invading. (**e**) Fluctuation free relative growth rate of the zooplankton when the zooplankton is invading. (**f** ) Relative nonlinearity in *S* when the zooplankton is invading. For each panel, the coexistence regime reached by the *SIV Z* model after 20 years of simulations, without quadratic mortality terms for the predators, is represented as the black contour. N.D.: Not defined.

To further characterize the underlying mechanism underpinning the relative nonlinearity in *S*, we decompose the average growth rates of the free virus and the zooplankton in the coexistence regime (*SIV Z* model) with or without fluctuations of the *I* and *S* classes (Methods, equations 28-30). We find that the fluctuation-free growth rate of the free virus, *i*.*e*., considering the concentration of the *I* and *S* class to be constant (set as their mean over a period cycle of *S*), varies from near zero to extremely high values (*>* 10^3^ *d*^−1^, Figure S7a). As expected from the formula of the free virus growth rate (equation 25), this result comes from a negative linear component in *I* for the growth rate of the free virus (Figure S7b). The inclusion of the infected class thus has a detrimental effect on the growth rate of the free virus, which in turn facilitates the coexistence. The lower clearance rate (*g*_*Z*_) of the zooplankton compared to the infection rate of the virus 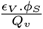, which is the viral clearance rate in the *SV Z* model) is effectively compensated by a reduced growth rate of the free virus population due to the dynamics of the oscillating infected population. In turn, this reduced growth rate of the free virus allows the zooplankton to take better advantage of the susceptible cells through the relative nonlinearity in *S* described in the MCT analysis. For the zooplankton, as expected from the linear growth rate of the latter, we observe no significant nonlinearities in the coexistence regime.

#### The inclusion of resistance facilitates coexistence through resource partitioning

We assess the effects of inclusion of a resistant phytoplankton type alongside the susceptible phytoplankter, the *SV RZ* model. We examine two scenarios: one where the resistant type exhibits partial resistance with a ratio of 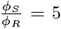, and another where the resistance is complete, meaning *ϕ*_*R*_ = 0 (fully resistant type). For the partially resistant type, a coexistence regime is observed in the region characterized by viruses of moderate virulence (Figure 4i). In contrast, for the fully resistant type, this region extends into the domain characterized by very virulent viruses (Figure 4q). As anticipated, these coexistence regions are located within the part of the virulence spectrum where the virus excludes the grazer in the *SV Z* model (Figure 4a versus Figure 4i, q). Intuitively, the introduction of a resistant type decreases the viral growth rate, while allowing the zooplankton to graze on the resistant type, which experiences reduced susceptibility to viral effects. To illustrate this, we plot the dynamics of the *SV RZ* model within the coexistence regime (Figure S8). Two distinct dynamic patterns emerge in this regime: either slowly dampened oscillations or sustained periodic oscillations. For a moderately virulent virus, the oscillations persist after 20 years of simulations, with the virus and the resistant type oscillating in opposite phases (Figure S8a, c). In the case of a slightly more virulent virus and a partially resistant type, the oscillations of the resistant type and the zooplankton dampen more quickly, while those of the susceptible type and the virus remain coupled (Figure S8b). Finally, for moderately virulent viruses, in the case of full resistance, we observe a stable equilibrium (Figure S8d). Importantly, adding the resistant type reduces the total virus abundance, and as viral virulence increases, the system transitions from dominance of the susceptible cell type to the resistant cell type. Overall, these results confirm that introducing a resistant type creates resource partitioning between the virus and the zooplankton, with the virus mainly targeting the susceptible type (though depending on the relative abundances of the *S* and *R* type), while the zooplankton consumes both types, primarily feeding on the more abundant one.

To further investigate this model, we set the latent period at 0.37 *days* ( ≈ 9 h, based on the life history trait model) and examine the impact of resistance strength with a constant cost on the growth rate of the *R* type (Figure S9). As the virus virulence increases, the resistance strength required to achieve the coexistence state also increases (Figure S9a). Adding the infected class (*SIV RZ* model, Figure 1f, equation 5) the state where the zooplankton is excluded in the *SV RZ* model now supports coexistence between the *S* and *R* types (Figure S9a versus S9e). Finally, we also test for an intracellular resistance model by modulating the probability of entering the infected state (parameter *ϵ*_*VR*_). In this case, the coexistence state is reached at lower resistance strengths for very virulent viruses (Figure S9a, e versus S9i, m). These findings suggest that extracellular resistance might be more favorable to invade a population of susceptible cells and become dominant.

#### The inclusion of higher-order mortality terms for predators facilitates coexistence and stabilization

In ocean ecosystem models, higher-order mortality terms represented by quadratic mortality terms are often used for grazers as a closure term for the system (*e*.*g*., Dutkiewicz et al. 2020) and have also been used as a loss term for viruses (Talmy et al. 2019a; Beckett et al. 2024). For the zooplankton, the quadratic mortality represents top-down control by higher predators, assumed to scale with zooplankton density. For the virus, the quadratic mortality represents losses due to unspecific binding *e*.*g*. collision with particulate matter. We find that the inclusion of a quadratic mortality term (Steele 1974) for the zooplankton in the *SV Z* model facilitates the coexistence regime in the adsorption rate region where the virus was previously excluded (Figure 4b versus 4a). Similarly, the inclusion of a quadratic mortality term for the virus facilitates coexistence in the region where the zooplankton was previously excluded (Figure 4c versus 4a). The inclusion of both terms results in both regions entering the coexistence regime (Figure 4d). In the *SIV Z, SV RZ* and *SIV RZ* models, the effects are similar (Figure 4 and Figure S9). Notably, for very virulent viruses and a partially resistant type 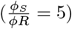, the inclusion of these higher-order loss terms also facilitates a coexistence state (of *V* and *Z*) where the *R* type dominates, excluding the *S* type (dark brown regions, Figure 4j, k, l, n, o, p). Interestingly, the region of *R* type dominance is larger for extracellular resistance than for intracellular resistance. This further suggests that extracellular resistance might be more beneficial assuming the same cost of resistance (Figure S9d, h versus S9l, p). Additionally, in the *S*(*I*)*V RZ* models, the inclusion of the virus quadratic mortality extends the region where the *S* type dominates, excluding the *R* type (light brown region, Figure 4k, l, o, p, s, t, w, x). For the *S*(*I*)*V RZ* model with a partially resistant type, as the virulence of the virus increases, there is a succession of coexistence states: first, the *S* type dominates (light brown region), then the *S* and *R* types coexist (gold brown region), and finally, the *R* type dominates (dark brown region, Figure 4l, p). It should be noted that the cases of *S* and *R* type dominance correspond to the same equilibrium. The only difference is the cost to the growth rate for the latter. These results suggests that higher-order losses favors phenotype selection by favoring regimes of coexistence either dominated by the susceptible or partially resistant type.

It is well known that quadratic mortality terms have a stabilizing effect on community dynamics (Poethke and Kirchberg 1987). To verify this in our system, we derive approximate model equilibria for the *SIV Z* and *SIV RZ* models in two cases: *d*_*Z*2_ *>* 0 with either *d*_*V* 2_ = 0 or *d*_*V* 2_ *>* 0 (see Supplementary Information for the mathematical derivations, equations S14-48, and Figure S10-14). We simulate the dynamics of the models and calculate the approximate theory (Methods) for four types of small phytoplankton: a small *Prochlorococcus*, a small *Synechococcus*, a picoeukaryote (non-diatom), and a small diatom (Table S1). We find a strong agreement between the approximate theory and the simulations (*SIV Z*, Figure S10 and *SIV RZ*, Figure S12), including system stability (shown in Figure S11 and S13), when compared to the Fourier analysis (Figure S14). The quadratic mortality terms strongly stabilize the system, with the viral quadratic mortality term acting to stabilize viral population dynamics for more virulent viruses. The oscillation dynamics is still maintained only for very virulent viruses and long latent periods (Figure S14). As expected from encounter rate theory (Huisman and Weissing 1999; Levin et al. 1977; Talmy et al. 2019b), the coexistence regime shifts toward more virulent viruses for larger organisms (Figure S10 and S12). The approximate theory is less accurate for larger organisms (diatoms), particularly for very virulent viruses in the *SIV Z* model (Figure S10p) and for long latent periods in the *SIV RZ* model, but it still effectively captures the coexistence regime (Figure S12p). Overall, these results underscore the stabilizing influence and broader coexistence regimes that emerge when higher-order loss processes, formulated as quadratic mortality terms, are included in biogeochemical models.

### Toward the integration of viral lysis in ocean ecosystem models

#### The effect of including a virus: critical transitions and impact on the system

A crucial challenge in implementing viral lysis models in ocean ecosystem models is in assessing the viral impact on the ecosystem and identifying ecological parameter ranges that produce realistic concentrations of different biogeochemical tracers. To address this issue, we first analyze the impact of including the virus on system dynamics and equilibria for the set of simulations including quadratic mortality terms (generally used in large scale ecosystem models) (Figure 6 for count concentrations and Figure S15 for nitrogen molar concentrations). We use a fixed latent period for our model *Prochlorococcus*, as determined by the life history trait model (Figure S3-4, Table S1). For all models we consider, mildly virulent viruses are excluded after approximately one year of simulation (Figure 6a, e, i, m). As the virulence of the virus increases, stable coexistence equilibria are first enabled by the addition of zooplankton quadratic mortality (Figure 6b), but these equilibria become unstable for more virulent viruses (Figure 6d, e). High virus count concentrations are reached in this case, generally much higher than the phytoplankton count concentration. Adding a viral quadratic mortality term strongly decreases the equilibrium virus concentration below the phytoplankton concentration (Figure 6f). As virulence increases, the virus concentration increases, leading to a decrease in phytoplankton concentration (Figure 6g). In the cases of very virulent viruses, virus concentration decreases and is accompanied by a sharp decline in phytoplankton concentration (Figure 6h). In the case of a partially resistant type, the same behavior is observed (Figure 6j, k), except for very virulent viruses, where the regime allows a new virus concentration peak where the resistant type dominates but the zooplankton is excluded (Figure 6l). For the fully resistant type (Figure 6m-p) coexistence with the zooplankton is maintained for very virulent viruses (Figure 6p).

**Figure 6.**
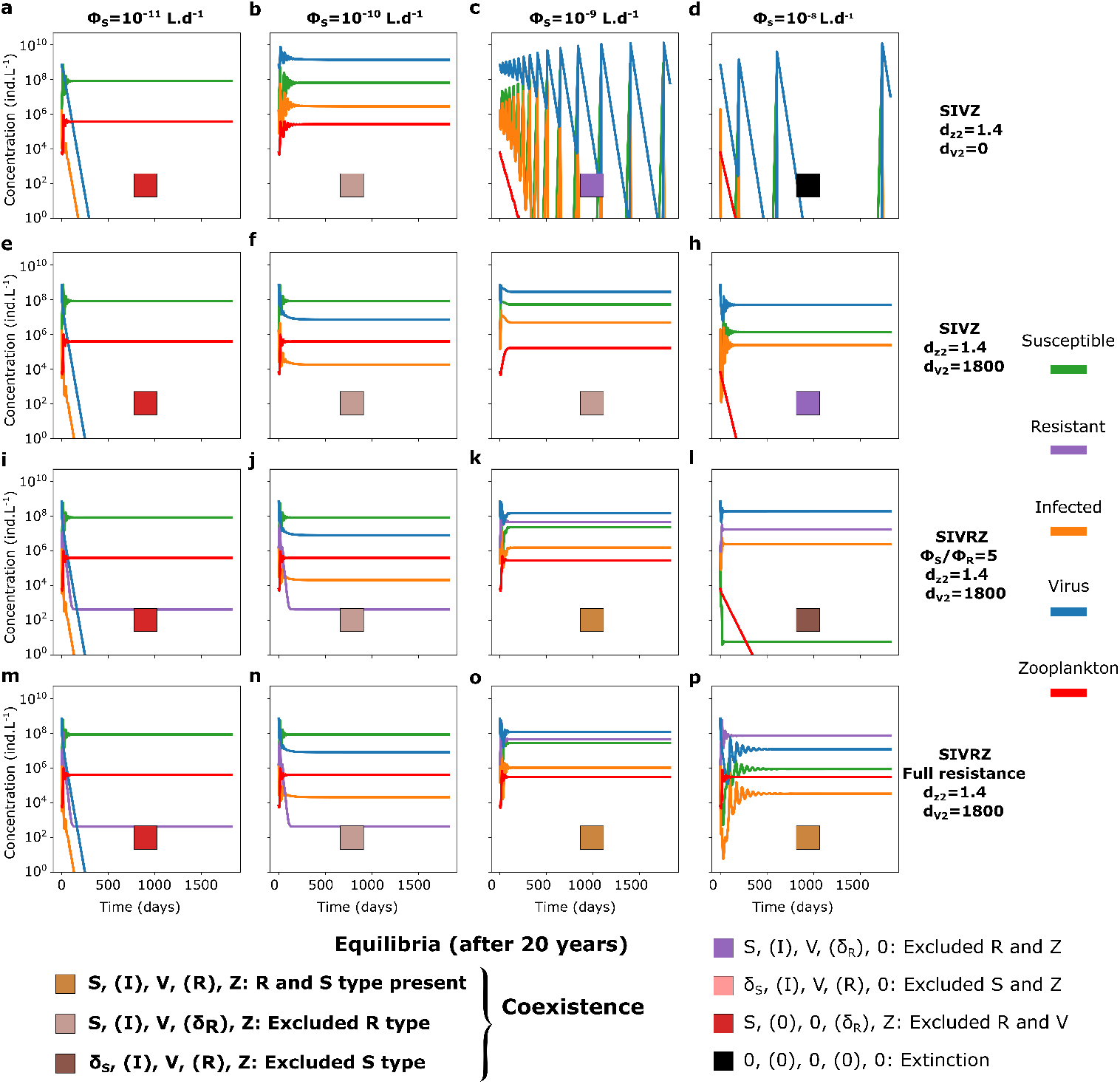
Model output of five year time series of the *SIV Z* and the extracellular *SIV RZ* models for *Prochlorococcus* for four different adsorption rates of the virus and different resistant type strength. Adsorption rate from *ϕ*_*S*_ = 10^−11^ *L*.*d*^−1^ to *ϕ*_*S*_ = 10^−8^ *L*.*d*^−1^ (*ϵ*_*V*_ = 1) for the *SIV Z* model (**a**-**d**) without and (**e**-**h**) with the quadratic mortality term of the virus, for the *SIV RZ* model (**i**-**l**) with a partially resistant type and (**m**-**p**) a fully resistant type, both with the quadratic mortality term of the virus. The tracers’ molar concentrations were converted in count concentrations using the cellular and virion quotas (see Figure S13 for time series in molar concentrations). A latent period of 0.37 *days* and a burst size of 15 were used as parameterized by the life history trait model (see Methods). The quadratic mortality terms are in .(*µmolN*.*L*^−1^)^−1^.*d*^−1^.

To better understand these effects, we visualize the equilibrium concentrations of total phytoplankton, virus, and zooplankton (averaged over the last year in the case of oscillations) across the virulence spectrum and the latent period parameter space for the different models in the set of simulations that include the quadratic mortality terms (Figure 7a-o). The total concentration of phytoplankton is significantly decreased by the presence of the virus only when the virus becomes sufficiently virulent, its encounter rate (infection rate) surpassing that of the zooplankton (Figure 7a). The total phytoplankton concentration decreases from ∼ 10^−1.2^ to ∼ 10^−3.1^ *µmolN*.*L*^−1^ ( ∼ 10^8^ to 10^6^ *ind*.*L*^−1^), slightly less than two orders of magnitude, for the most virulent viruses. Including a partially resistant cell type reduces this effect (decreasing to ~ 10^−2^ *µmolN*.*L*^−1^ *i*.*e*. ∼ 10^7.25^ *ind*.*L*^−1^), and the transitions between the three coexistence regimes (Figure 4p) become apparent (Figure 4b). In the case of the fully resistant type, the *R* type dominated regime is not reached (Figure 7c). The decomposition of the phytoplankton population into susceptible, infected, and resistant types, along with the percentage of resistant cells (relative to *S* + *R*), shows a rapid transition from a susceptible-dominated population to a resistant-dominated population as virulence increases (Figure S16g-p). For the zooplankton, as the virulence of the virus increases, its equilibrium concentration is similarly affected (Figure 7g-i, decreasing from ∼ 10^−1.3^ to ∼ 10^−2.3^ *µmolN*.*L*^−1^ *i*.*e*. ∼ 10^5.5^ to 10^4.5^ *ind*.*L*^−1^) and is excluded for the most virulent virus in the *SIV Z* model (Figure 7g). For the virus, a relatively broad concentration peak is observed for moderately virulent viruses in the *SIV Z* model (Figure 7e), which narrows in the *SIV RZ* model as the resistant type takes over (Figure 7e-f). For the *SIV RZ* model with a partially resistant type a second peak of virus concentration is observed corresponding to the regime of R dominance (Figure 7e). Conversely, in the case of the fully resistant type, the coexistence of the *R* and *S* types for very virulent viruses is enabled (full coexistence regime, Figure 4x). Surprisingly, in this case, the equilibrium viral concentration becomes independent of the latent period and decreases relative to viral concentrations in the *S* and *R* types dominated regimes (Figure 7e, f and the theoretical equilibria, equation 42-43). The quadratic mortality term used for the virus appears to be relatively high (Beckett et al. 2024), so the maximum concentrations reached are around ~ 10^−3.4^ *µmolN*.*L*^−1^ *i*.*e*. ∼ 10^8.5^ *ind*.*L*^−1^. Overall, extending the latent period (*τ* ) shifts the transitions between states towards more virulent viruses, with a modest effect for *τ <* 1 *day* and a more pronounced effect as the latent period is extended. Extending *τ* also narrows the zone of maximum viral concentration. We also estimate the impact of the virus on Net Primary Productivity (NPP, Figure 7j-l) a key metric for ocean ecosystems. NPP is directly related to phytoplankton concentration (equation 37, Figure 7a-c). Consequently, very virulent viruses have a dramatic effect that decreases NPP (Figure 7j). Although the decrease in NPP is partially mitigated by the inclusion of the resistant type, the resistance cost decreases NPP when comparing the *S* type dominant regime to the mixed or *R* type dominant regimes (Figure 7k-l).

**Figure 7.**
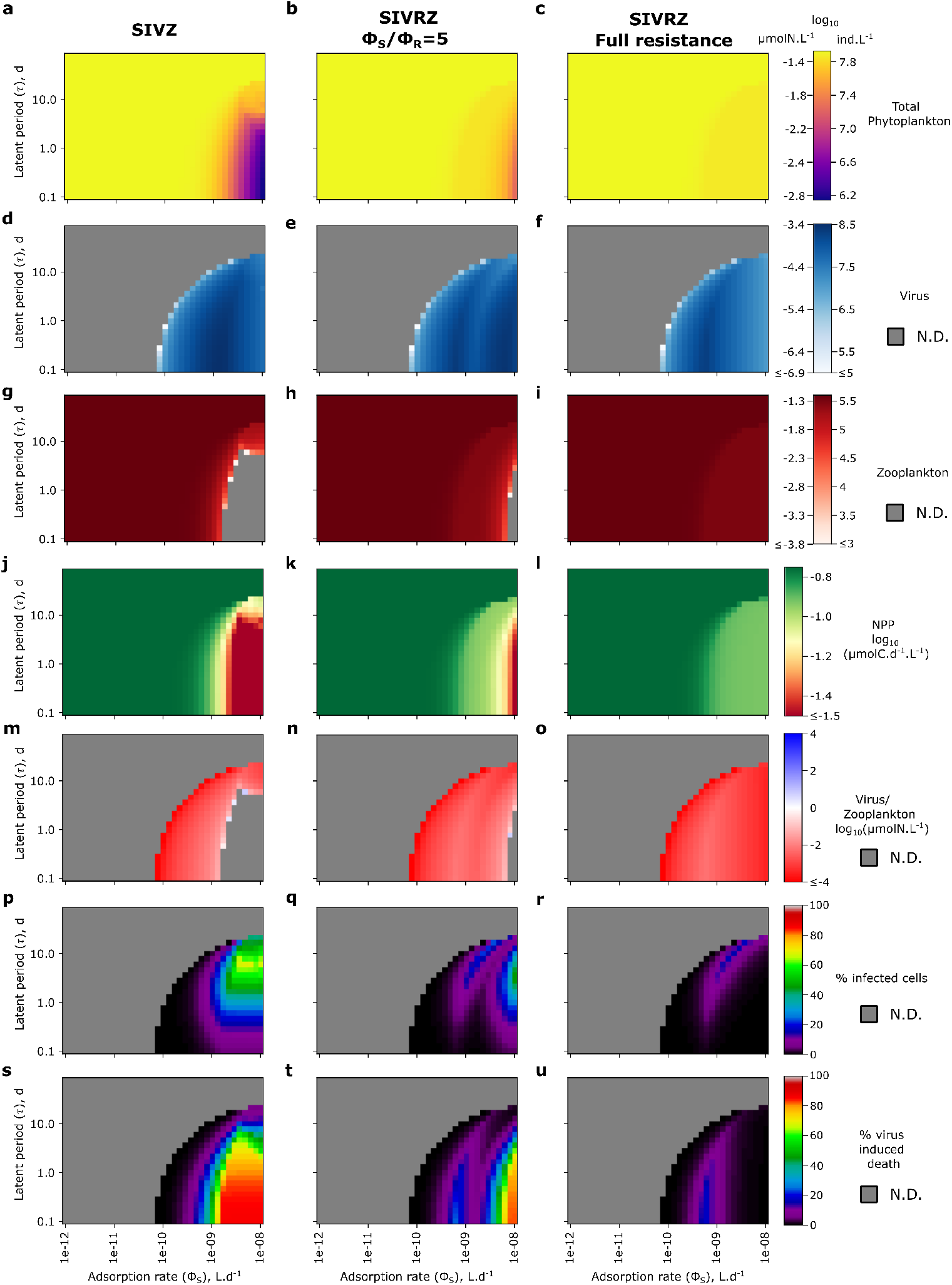
Equilibrium concentration and ecological features of the *SIV Z* and the extracellular resistance *SIV RZ* models for *Prochlorococcus* across the adsorption rate and latent period parameter space. From left to right panels: *SIV Z* model, *SIV RZ* model with a partially resistant type: 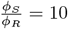, and *SIV RZ* model with a fully resistant type. (**a**-**c**) Total phytoplankton concentration. (**d**-**f** ) Virus concentration. (**g**-**i**) Zooplankton concentration. (**j**-**l**) Ratio of virus to zooplankton concentrations. (**m**-**o**) Percentage of infected cells. (**p**-**r**) Percentage of virus induced death. In panels (**m**–**o**), the grey areas fall outside the coexistence domain, so these metrics are not defined. N.D.: Not Defined: the tracer is excluded or one of the necessary tracers to define the metric is excluded.

We next investigate the relative roles of the two predators in the system, along with the percentage of infections induced by the virus at equilibrium (Figure 7m-u). The zooplankton is generally more abundant in molar concentration than the virus, with a minimal molar ratio occurring at the maximum of free virus concentration (Figure 7m-o). For short latent periods (*τ <* 1 day), the percentage of infected cells is generally low, transitioning from near 0 to 10% in the *SIV Z* model as virulence increases (Figure 7p). For the *SIV RZ* model, the highest infection percentages are found around the virus concentration peaks, although they remain quite low (*<* 10 %) when the latent period is sufficiently short (Figure 7p-r). Extending the latent period generally increases importantly the percentages of infected cells at equilibrium which also increases with the viral adsorption rate although to a lesser extent (Figure 7p-r). The percentage of virus-induced mortality increases drastically with the virulence of the virus. For the *SIV Z* model, it transitions from nearly 0% to ∼ 90% over ∼ 1.3 orders of magnitude in the virulence spectrum (Figure 7s), highlighting the strong sensitivity of the system to the virulence of the virus. This effect is primarily driven by the decrease in abundance of the zooplankton and does not necessarily imply a much higher percentage of infected cells (for short latent periods such as the one of cyanophages). Finally, adding the partially resistant type creates two peaks in the maximal percentage of virus-induced mortality (Figure 7t), while the fully resistant type results in a single peak (Figure 7u). These two peaks align with the two peaks of viral concentration corresponding to the regimes of *S* and *R* dominance respectively.

#### Bridging model and field ecology: models suggest the dominance of viral resistance

We assess the performance of each model against marine field measurements to identify parameter combinations that yield potentially realistic ecological outcomes. We define concentration ranges and targets for total phytoplankton, viruses, and zooplankton, together with expected ranges for the percentage of infected cells and virus-induced mortality in two idealized epipelagic environments (Table S4; Mojica et al. 2016; Carlson et al. 2022; Schartau et al. 2010; Methods; Figure 8a). To illustrate the *in situ* measurements for the oligotrophic environment, we show the distributions from Carlson et al. (2022) in the North Pacific Subtropical Gyre (Figure S17, latitude*<*33°N). The measured concentrations in the literature were reported in *cell*.*L*^−1^ and *virion*.*L*^−1^ so we convert to this unit for this analysis. Using the set of default parameters (Table S2), the *SIV Z* model minimizes the distance to target concentrations in the model mesotrophic environment for moderately virulent viruses near the maximum concentration of free virus (Figure 8b). Decomposing this distance among the different tracers shows that it is minimized around the peak concentration of free virus for all tracers, except for the zooplankton in the *SIV RZ* model with the fully resistant type (Figure S18). For the *SIV RZ* model with a partially resistant type, the minimum distance to the target concentrations is found in the regime dominated by the resistant type (Figure 8c). Conversely, the model with a fully resistant type shows poorer performance (Figure 8d). When the target concentration assumptions are removed, the entire coexistence regime falls within realistic concentration ranges for all models (Figure 8e-g). However, further constraining the output with the percentage of infected cells, virus-induced mortality, and a *V/P* count ratio greater than 1 (Figure S19), a much narrower region of the virulence spectrum and latent period parameter space aligns with observed empirical measurements (Figure 8h-j). These regions rarely encompass the coexistence regime including both the *S* and *R* types in abundance; instead, they mainly occur in regions where only one type is dominant (Figure 4 versus Figure 8h-j).

**Figure 8.**
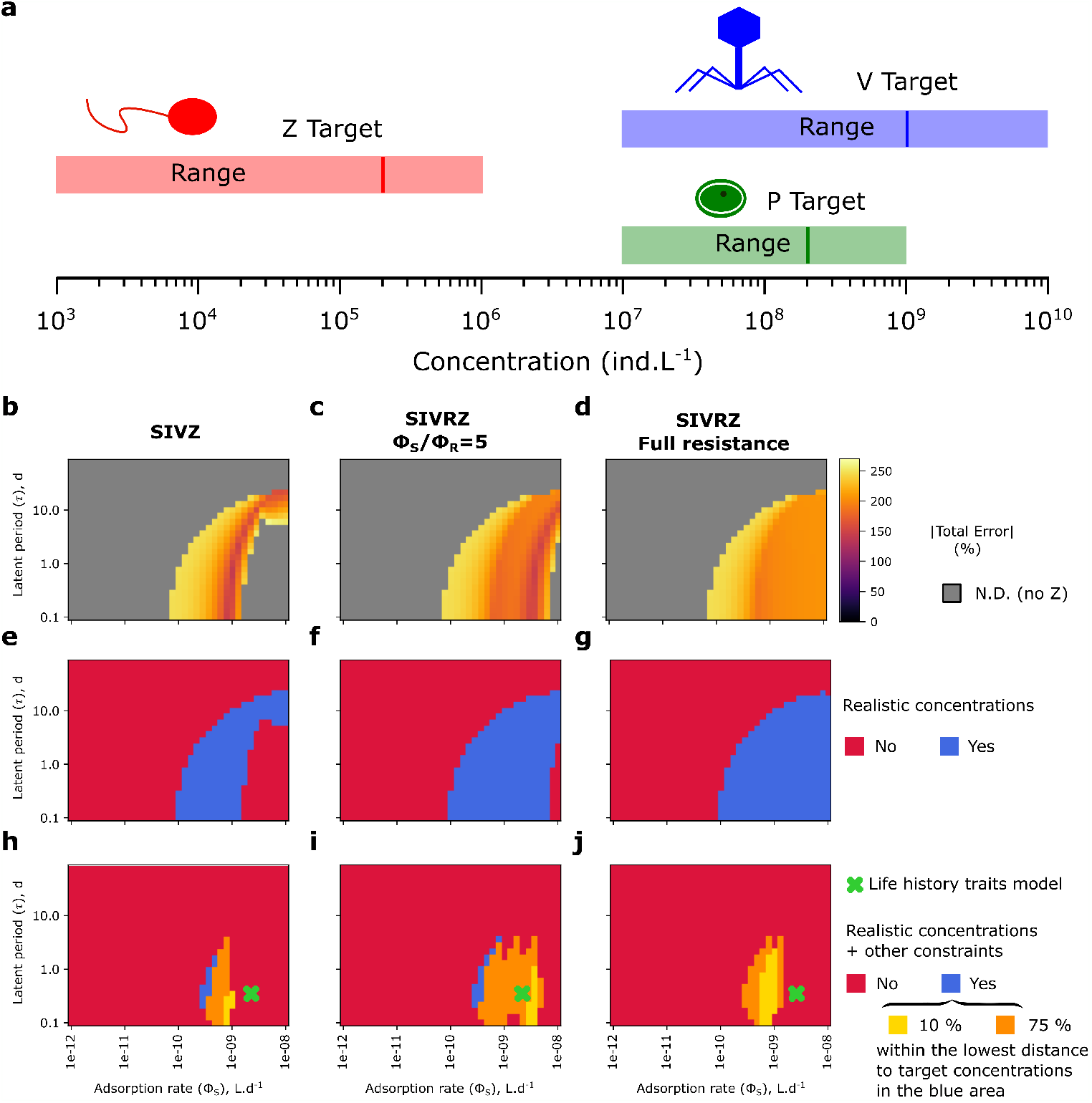
Model output compared to expected concentration and ecological features measured in the field across the adsorption rate and latent period parameter space. (**a**) Target and concentration ranges in a mesotrophic environment for the zooplankton (red), *Prochlorococcus* (green) and the virus (blue). (**b**-**d**) From left to right panels: *SIV Z* model, *SIV RZ* model with a partially resistant type: 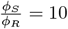, *SIV RZ* model with a fully resistant type: (**b**-**d**) Distance to target concentrations defined in Table S4, (**e**-**g**) Realistic concentrations in the range defined in Table S4 and (**h**-**j**) Realistic concentrations and constraints on other expected ecological features (percentage of infected cells, percentage of virus-induced mortality and V/P ratio, see Table S4). The green cross represents the life history traits model. N.D.: Not Defined: at least one tracer is excluded.

Importantly, the “realistic” ecological parameter space is found within a virulence range with a lower adsorption rate, compared to a theoretical encounter model by Murray and Jackson (1992), where the swimming speed of the phytoplankton (zero in the case of cyanobacteria) is considered to have a multiplicative effect on diffusion (Figure 8h-j, equation 13, Methods). The *SIV RZ* model with partial resistance represents the best match, as it minimizes the error to target concentrations close to the life-history trait model while also satisfying the other ecological constraints that we impose (Figure 8i). In general, these results suggest that including a level of extracellular resistance (compared to the encounter rate model) to the virus leads to a more accurate representation of the equilibrium achieved (Figure 8i).

To further test this hypothesis, we leverage the theoretical equilibria of the *SIV Z* model (Supplementary Information). We allow for resistance and cost on the maximal growth rate, to minimize model error in modeling target concentrations for both a mesotrophic and an oligotrophic environment (avoiding overfitting). The procedure is followed for four considered types of phytoplankton: *Prochlorococcus, Synechococcus*, a picoeukaryote (non-diatom), and a small diatom (Table S4). The theoretical equilibria allow us to exhaustively explore a large ecological parameter space using a grid search approach. We choose to focus on the intracellular and extracellular resistance, the resistance cost, the linear and quadratic mortality of the virus (that can be set to equal zero), and the quadratic mortality of the zooplankton (Table 1). We define the extracellular resistance as the ratio between the theoretical encounter rate and the fitted adsorption rate 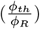 while the intracellular resistance is defined as the inverse of the probability of infection where new virions are produced 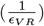. We find that the optimal extracellular resistance is around ∼ 3 − 20 for *Prochlorococcus, Synechococcus* and the picoeukaryote, while it is ∼ 290 for a small diatom. Intracellular resistance factors were found to be in the same order of magnitude as extracellular resistance. Multiplying both types of resistance, we get total resistance factors of 46, 29, 5 and 292 respectively for *Prochlorococcus, Synechococcus*, the picoeukaryote and the small diatom in order to minimize distance to target concentrations. Among the 200 best-fitting models, extracellular and intracellular resistance were significantly anticorrelated (*p <* 0.0001) for *Prochlorococcus, Synechococcus*, and the small diatom, with correlation coefficients of *r* = –0.58, –0.71, and −0.53 respectively. The correlation was weaker for the eukaryote (*r* = −0.22, *p <* 0.01), but the best-fitting models exhibited a narrow range of extracellular resistance values. A resistance cost (1 − *ζ*) of ∼ 0 − 0.5 is found to be a good fit across organisms although other parameters might also contribute to decreasing the effective growth rate in larger ocean models. The linear and quadratic mortality of the virus are also found to be lower compared to the default parameter. The quadratic mortality of the virus appeared relatively poorly constrained across the 200 best-fitting models for the four phytoplankton types. Finally, the best-fitting models suggest a higher quadratic mortality for the zooplankton compared to the default value used although the fitted range of the 200 best-fitting models remained wide and it might depend on the grazing function used.

We typically find low fitting errors in the differences between target and the modeled concentrations of the best fitting models (Table 2). Target virus concentrations are generally achieved for all 4 organisms. Generally, the percentages of infected cells reported by the models are small for cyanobacteria (0.75 to 2.0% in the oligotrophic environment) and 3.1% for the picoeukaryote, which are in good agreement compared to average oligotrophic field measurements for cyanobacteria (Carlson et al. 2022; Beckett et al. 2024 and Figure S17). In the mesotrophic environment, the percentage of infected cells were 1.4 and 4.1% for *Prochlorococcus* and *Synechococcus* respectively which is below peak percentages observed in such environments (Carlson et al. 2022). The observed peaks most likely come from out of equilibrium dynamics or increase in latent period which our model does not allow here. Our fitted models also correspond to a relatively low percentage of virus-induced mortality between 7.4% (*Prochlorococcus*) and 17% (*Synechococcus*) in agreement with oligotrophic field measurements (Beckett et al. 2024). In the mesotrophic environment, these percentages of virus-induced mortality increase to a range of ∼ 12% (*Prochlorococcus*) to ∼ 38% (picoeukaryote). For the small diatom, the percentages of infected cells reaches ∼ 8% (most likely due to the the longer latent period) in the mesotrophic environment but still corresponds to a relatively low percentage of virus-induced mortality (15%).

**Table 2.** Model performances compared to target concentrations for a simplified oligotrophic and mesotrophic epipelagic environment for the four types of phytoplankton. The absolute percentage error (rows Best and Range) represents the total error across target environments cumulating errors to the target concentrations of the phytoplankton, the zooplankton and the virus. For each environment, the “minimum error” corresponds to the error of best model for which the average absolute error across environments is minimized. The “Range” reflects the minimum and maximum values of the first 100 best models. For *T otP, V*, and *Z*, the equilibrium value is indicated in *ind*.*L*^−1^ and the minimum percentage error to the target is in parenthesis. For % Inf, % V Kill and *V/P* the equilibrium value is shown.

| Ocean |  | *Prochlorococcus* | *Synechococcus* | Picoeukaryote | Diatom |
| --- | --- | --- | --- | --- | --- |
| Oligotrophic | Minimum total error (%) | 9 | 24 | 36 | 78 |
|  | Error Range (%) | 5 – 41 | 8 – 64 | 14 – 128 | 12 – 158 |
| | Tot $P$ | $1.4 \cdot 10^8$ (−6.2%) | $2.3 \cdot 10^7$ (+13.6%) | $1.1 \cdot 10^7$ (+9.2%) | $5.5 \cdot 10^5$ (+10.6%) |
| | $V$ | $5.08 \cdot 10^8$ (+1.7%) | $4.9 \cdot 10^8$ (−2.5%) | $9.4 \cdot 10^7$ (−5.7%) | $1.7 \cdot 10^8$ (−14.3%) |
| | $Z$ | $1.01 \cdot 10^5$ (+1.2%) | $1.1 \cdot 10^5$ (+7.5%) | $1.2 \cdot 10^4$ (+21%) | $1.5 \cdot 10^4$ (+53.4%) |
|  | % Inf | 0.75 | 2.05 | 3.1 | 1.7 |
|  | % V kill | 7.4 | 17.4 | 27.6 | 6.5 |
| | $V/P$ | 3.6 | 21.3 | 8.5 | 310 |
| Mesotrophic | Minimum total error (%) | 21 | 29 | 53 | 28 |
|  | Error Range (%) | 4 – 44 | 6 – 64 | 13 – 70 | 16 – 88 |
| | Tot $P$ | $2.1 \cdot 10^8$ (+4.8%) | $3.7 \cdot 10^7$ (−25%) | $2 \cdot 10^7$ (−0.8%) | $1.5 \cdot 10^6$ (−25.0%) |
| | $V$ | $10^9$ (−0.3%) | $1 \cdot 10^9$ (+2.5%) | $2 \cdot 10^8$ (+1.4%) | $1 \cdot 10^9$ (+2.9%) |
| | $Z$ | $1.7 \cdot 10^5$ (−15.7%) | $2 \cdot 10^5$ (−0.4%) | $2.4 \cdot 10^4$ (−51.0%) | $4.8 \cdot 10^4$ (−0.3%) |
|  | % Inf | 1.4 | 4.10 | 6.3 | 7.9 |
|  | % V kill | 12.0 | 25.7 | 38.4 | 15.1 |
| | $V/P$ | 4.8 | 27 | 10 | 690 |
| Weighted Average | Minimum error (%) | 15 | 26 | 47 | 45 |
|  | Error Range (%) | 15 – 24 | 26 – 37 | 47 – 52 | 45 – 64 |

## Discussion

We proposed several models that incorporate viral lysis of phytoplankton designed for integration into large-scale ocean ecosystem models. Our approach transforms the *SIV* count model (Levin et al. 1977; Weitz 2015) into an elemental concentration *SIV* model that incorporates cellular and virion quota (Jover et al. 2014). We explicitly wrote elemental fluxes equations linked to the viral “shunt” (Fuhrman 1999; Wilhelm and Suttle 1999) and the viral “shuttle” (Sullivan et al. 2017; Guidi et al. 2015; Zimmerman et al. 2020). This step is essential for integration into large-scale biogeochemical models, as it accounts for the distinct stoichiometry of viruses and their hosts, particularly their C:N:P ratios (Redfield 1934; Jover et al. 2014) which is expected to have consequences on DOM and POM stoichiometric composition. To validate the feasibility of integrating viral lysis models into large-scale biogeochemical frameworks, we explored the mechanisms that could facilitate the coexistence of both a virus and a zooplankton preying on the same phytoplankton. Inclusion of the infected class (*I*) had a dual effect, either enabling limit cycles or driving the system to extinction through oscillatory dynamics. These oscillations in the infected class may play a crucial role in spatially resolved models by inducing out-of-equilibrium conditions, particularly in transition zones (Carlson et al. 2022). Next, the addition of a resistant type facilitated the coexistence of the two predators through niche partitioning. In particular, for the most virulent viruses, full resistance enabled the coexistence of both susceptible and resistant types, while partial resistance led to the exclusion of the susceptible type. The latter case (partial extracellular resistance) corresponds to the coexistence regime of a less virulent virus and includes an eventual cost to the growth rate of the phytoplankton. The inclusion of this kind of resistant type in biogeochemical models, although it may be computationally costly, could lead to the system alternating between different coexistence regimes, with a potential decrease of primary productivity due to the resistance cost. The resistance cost may depend on the mutation (Bohannan and Lenski 2000; Chao et al. 1977) and follow trade-offs (Våge et al. 2013). In addition, the range of resistance, *i*.*e*. the resistance to multiple viruses, which we do not resolve here, can vary (Avrani and Lindell 2015). Importantly, natural populations might evolve to a reduced resistance range and favor maximal growth rates through compensatory mutations (Avrani and Lindell 2015). Finally, the inclusion of higher-order quadratic mortality terms for the predators expanded the coexistence regimes by stabilizing the system. Quadratic mortality terms also have a controlling role on the concentrations reached by the predators. These terms are not well constrained and here we fitted values that are lower than those previously reported for viruses (Beckett et al. 2024) and higher than those used for zooplankton (Dutkiewicz et al. 2015a). Importantly, in large scale ecosystem models, other mechanisms of coexistence might be at play like advection by currents and the size fractionation of zooplankton grazers (Dutkiewicz et al. 2020).

We evaluated the impact of introducing the virus into the system, focusing on its effects on phytoplankton mortality and abundance, as well as the partitioning between zooplankton and virus-induced mortality. The outcomes were strongly influenced by the virulence of the virus: mild viruses had minimal impact, whereas highly virulent viruses induced substantial changes in the system’s dynamics and equilibrium. As previously reported, a maximal concentration of free viruses was observed for mildly virulent strains, whereas the most virulent viruses were less abundant (Maidanik et al. 2022). This finding is consistent with field observations of cyanophages infecting cyanobacteria populations in the global oceans (Maidanik et al. 2022). In addition, the system exhibited strong sensitivity, characterized by steep transitions in its ecological properties. In particular, we observed rapid shifts in the percentage of infected cells and virus-induced mortality across the virulence spectrum. The percentage of infected cells was predominantly influenced by the latent period, while virus-induced mortality showed a pronounced increase with higher viral adsorption rates. Thus, a high percentage of infected cells does not necessarily correspond to a high percentage of virus-induced mortality, and vice versa. Paradoxically, a highly virulent virus with a short latent period, despite its low abundance, can drive substantial phytoplankton mortality even when the percentage of infected cells at equilibrium is low. In the model, this phenomenon is driven primarily by the partial exclusion of zooplankton, which may, for instance, explain the negative correlations between viral and grazing rates observed in field studies (Biggs et al. 2021). These results further suggest that transient dynamics could readily emerge in spatially explicit models, potentially explaining the variability in ecological properties observed in the field, such as the percentage of infected cells and virus-induced mortality (Mojica et al. 2016; Biggs et al. 2021; Carlson et al. 2022; Beckett et al. 2024).

Finally, we compared the outputs of our models to field measurements of virus, phytoplankton, and zooplankton concentrations. Using the default ecological parameter set, the *SIV Z* and *SIV RZ* models performed similarly in minimizing the distance to target concentrations. However, the *SIV RZ* model aligned more closely with the life history trait model describing the encounter rate between phytoplankton and their viruses. Notably and as expected (Weitz et al. 2015), this result suggests that incorporating inefficiencies and/or resistance into the encounter rate model provides a better fit to empirical measurements. Interestingly, modulating both extracellular resistance, directly affecting the adsorption rate and intracellular resistance, that decreased the probability of infection leading to the production of new virions, were necessary to fit the models to field measurements with the relative, total resistance factor ranging from low to medium to high, for picoeukaryotes, cyanobacteria and diatoms, respectively. Nevertheless, a certain degree of uncertainty remains in this parameterization. Anti-phage intracellular defense mechanisms are well-documented (Labrie et al. 2010; Doron et al. 2018; Rousset et al. 2022), although some remain unidentified in marine cyanobacteria (Zborowsky and Lindell 2019). Notably, studies have shown that cyanophage resistance is predominantly extracellular against specialist phages, which target the most abundant high-light–adapted *Prochlorococcus* strains. In contrast, resistance is primarily intracellular against generalist phages, which infect the less abundant *Synechococcus* and low-light–adapted *Prochlorococcus* ecotypes (Sullivan et al. 2003; Dekel-Bird et al. 2015). These observations support our findings, suggesting that both types of resistance are important in marine phytoplankton. Future models may wish to include the potential for intracellular resistance to halt infections at various stages (Zborowsky and Lindell 2019). In our approach, we assumed that intracellular resistance reduced the probability of transitioning to the infected state that successfully lead to the production of new virions. This could be further explored by modeling a transition rate from the infected class back to the susceptible class or by using more complex models that include intermediate infected states (Hinson et al. 2023; Dominguez-Mirazo et al. 2024).

Importantly, the encounter rate and the adsorption rate can be influenced by several factors not explored in our models. The encounter rate represents the biophysical maximum rate of interaction, while the adsorption rate reflects the actual rate of successful attachment, which is a fraction of the encounter rate. For instance, not all virions attach to their host upon encounter, as receptor heterogeneity on the cell surface (Murray and Jackson 1992; Talmy et al. 2019b) and cell surface properties (Yamada et al. 2023) can limit attachment. Furthermore, turbulent diffusion can either increase or decrease encounter rates depending on the turbulence intensity, with high turbulence potentially reducing encounter rates (Stocker 2012; Pecseli et al. 2014). Additionally, the adsorption rate might be decreased due to nonspecific binding to other particles in the environment, such as cellular debris or other cells (Yamada et al. 2020), as well as through the formation of marine snow via the viral “shuttle” mechanism (Sullivan et al. 2017; Guidi et al. 2015; Zimmerman et al. 2020). This effect could be represented in a model as a Hill function inhibiting viral adsorption to the host (Igler 2022). The magnitude of this effect may vary significantly in large-scale ocean models and in field conditions due to the pronounced heterogeneity of marine environments. These effects may not be as pronounced as the extracellular resistance proposed here, especially in oligotrophic and mesotrophic environments, such as those modeled here, which have low POM concentrations (Tanioka et al. 2022) but high dissolved organic matter (DOM) concentrations (Hansell and Orellana 2021). The extracellular resistance used here can thus be viewed as the cumulative effect of these abiotic mechanisms in addition to biological resistance. Nevertheless, further research is needed to accurately account for and parameterize these effects.

Our community ecology model comes with caveats. First, despite identifying potential flux terms, we did not explicitly account for feedback with the microbial loop. This would require the explicit inclusion of heterotrophic bacteria and inorganic nutrients in the model (Weitz et al. 2015). Heterotrophic bacteria remineralize dissolved organic nitrogen (DON) to dissolved inorganic nitrogen (DIN), thereby increasing nutrient availability to phototrophs, which is expected, in turn, to increase primary productivity (Fuhrman 1999; Wilhelm and Suttle 1999; Weitz et al. 2015). Nonetheless, the system of ordinary differential equations we propose (equations 1a to 1c) and the fluxes to *DON* and *PON* (equation 1d and equation 1e) could be sufficient for integration of viral lysis into large-scale ocean ecosystem models. To fully close the model, one should also redistribute mortality terms to the *DIN, DON* and *PON* compartments. Future work will seek to couple models such as those we develop here into more complex ecosystem models.

Despite its relative simplicity, the *SIV RZ* model, with a dominant *R* type, can effectively recapitulate the observed ecology of viral infections in oligotrophic and mesotrophic environments. By integrating algebraic theoretical results, field measurements, and life history trait models, we are able to accurately reproduce viral and phytoplankton concentrations, percentages of infected cells, and virus-induced mortality in both prokaryotic and eukaryotic phytoplankton. We characterized the effect of adding an infected class, which can have a dual impact on system stability, either stabilizing or destabilizing, depending on the trade-off between the latent period and the adsorption rate of the virus. In addition, the inclusion of the *I* class enables the model to accurately recapitulate the percentages of infected cells observed in the field. The percentage of infected cells in our model does not necessarily reflect the percentage of virus-induced mortality, as it depends on the virulence of the virus and whether it impacts the zooplankton. Finally, our models suggest that incorporating significant extracellular resistance into biophysical encounter rate models may be critical for accurately modeling viral lysis in large biogeochemical models, which is consistent with the measured resistance of phytoplankton strains Here we developed only one host-one virus models and did not account for viral (Gregory et al. 2019) and phytoplankton diversity (Dutkiewicz et al. 2020). The inclusion of multiple host-virus pairs is expected to increase phytoplankton coexistence (Thingstad 2000; Flynn et al. 2022). However, in systems with multiple host-virus pairs it might be challenging to maintain coexistence of all virus-host pairs (*i*.*e*. certain virus might be excluded and not others). We also used a very generic model of lytic viral infection and neglected the potential uptake of virions by grazers (DeLong et al. 2022). Other types of viral infection cycles exist, such as persistent infections (Tuttle and Buchan 2020; Sanchez-Martinez et al. 2025), or chronic infections, for instance in *G. huxleyi*, where virions are continuously produced by exocytosis (Homola et al. 2024).

Integrating viral lysis in ocean ecosystem models may improve estimates of carbon cycling in the oceans and quantify the relative role of viruses and zooplankton in phytoplankton mortality. The inclusion of viruses might also reduce the magnitude of discrepancies between models shown to to be linked to the parameterization of grazing (Rohr et al. 2023). In closing, we anticipate that the inclusion of viral lysis into large ecosystem models will identify knowledge gaps and catalyze experimental and field-based studies to assess how viruses shape community ecology and biogeochemical cycles.

## Code and data availability statement

All code and data necessary to generate all figures and tables are available at https://github.com/PaulFremont3/SIVZ_coexistence/ and are archived on Zenodo at https://doi.org/10.5281/zenodo.15866512 (Frémont 2025).

## Acknowledgments

The authors acknowledge the University of Maryland supercomputing resources (http://hpcc.umd.edu) made available for conducting the research reported in this paper. This work is supported by grants from the Simons Foundation: no. 721231 to JSW, no. 549931 (Simons Collaboration on Computational Biogeochemical Modeling of Marine Ecosystems (CBIOMES)) and SFI-LS-Project-00011157 to SD, SFI-LS-Project-00010539 to DT and no. 721254 and SFI-LS-Project-00011154 to DL. Paul Frémont is supported by a postdoctoral fellowship from the Simons Collaboration on Ocean Processes and Ecology (SCOPE), funded by the Simons Foundation through grants awarded to JSW. Stephen J. Beckett and Joshua S. Weitz are investigators at the University of Maryland-Institute for Health Computing, which is supported by funding from Montgomery County, Maryland and The University of Maryland Strategic Partnership: MPowering the State, a formal collaboration between the University of Maryland, College Park and the University of Maryland, Baltimore. DT acknowledges NSF OCE grants no 2445508 and no 2023680. This is a contribution of the Simons Collaboration on Ocean Processes and Ecology (SCOPE). We thank members of the vDARWIN working group including Barbara Duckworth, Michael J. Follows, Oliver Jahn, and Daniel Muratore for constructive discussions and feedback.

## Author Contributions

Conceptualization: PF, SJB, DD, JSW; methodology: PF; investigation: PF; model development: PF, SJB, DD, EC, CLF, DL, DT, SD, JSW; code review: EC; writing—original draft: PF; writing—review & editing: PF, SJB, DD, CLF, DL, DT, SD, JSW; funding acquisition: JSW, SD, DT, DL.

