## Supplementary Information for "Coexistence of Photosynthetic Marine Microorganisms, Viruses, and Grazers: Toward Integration in Ocean Ecosystem Models"

#### From the $SIV$ count model to the $SIV$ concentration model

##### $SIV$ count model

In the count model, $S$, $I$ and $V$ are, respectively, in units of number of susceptible cells, infected cells and number of free virions per liter. The $SIV$ count model can be written as follow:

$$\frac{dS}{dt} = \overbrace{\mu.S}^{\text{growth}} - \overbrace{d_S.S}^{\text{mortality}} - \overbrace{\epsilon_V.\phi_S.S.V}^{\text{successful infection}} \tag{S1}$$

$$\frac{dI}{dt} = \overbrace{\epsilon_V.\phi_S.S.V}^{\text{successful infection}} - \overbrace{d_S.I}^{\text{mortality}} - \overbrace{\frac{1}{\tau}.I}^{\text{lysis}} \tag{S2}$$

$$\frac{dV}{dt} = \overbrace{\frac{\beta}{\tau}.I}^{\text{viral burst}} - \overbrace{d_V.V}^{\text{viral decay}} - \overbrace{\phi_S.S.V}^{\text{adsorption}}\ . \tag{S3}$$

In the above, $\mu$ is the division rate of susceptible cells ($S$), $d_S$ is the cell mortality rate, $\phi_S$ is the adsorption rate of viruses to susceptible cells, $\epsilon_V$ represents an intracellular resistance to the virus - that is the fraction of adsorption events that lead to a successful infection, and $d_V$ represents the decay rate of infectious virions. Infections have an average duration of $\tau$, denoting the latent period, after which a burst size, $\beta$, of new virions are released into the environment. Note that the dynamics of intracellular virus production are not represented in this framing. With the aim of incorporating this type of model within a larger ocean biogeochemical modeling framework, we can represent the "shunt" and "shuttle" of organic matter released by the lysis of the infected cells. The viral lysate is "shunted" to the dissolved organic matter compartment ($DOM$), which here is $DON$, dissolved organic nitrogen, as we use nitrogen as the currency element for this model, and "shuttled" to the particulate organic nitrogen compartment ($PON$). The flux to $DON$ and $PON$, including only terms related to the viral lysis, in $\mu molN.L^{-1}$ are the following:

$$J_{DON} = \overbrace{\gamma.(\frac{Q_p}{\tau} - \frac{Q_v.\beta}{\tau}).I}^{\text{shunt}}, \tag{S4}$$

$$J_{PON} = \overbrace{(1-\gamma).(\frac{Q_p}{\tau} - \frac{Q_v.\beta}{\tau}).I}^{\text{shuttle}}, \tag{S5}$$

where $Q_p$ and $Q_v$ are the respective phytoplankton cell and viral molar quotas in nitrogen.

##### $SIV$ concentration model

We define the following relationships between the cell/virion molar quota concentrations and counts:

$$\widetilde{S} = Q_p.S, \tag{S6}$$

$$\widetilde{I} = Q_p.I, \tag{S7}$$

$$\widetilde{V} = Q_v.V. \tag{S8}$$

By substituting these definitions (equations S6 to S8) into the system of differential equations for the $SIV$ model (equations S1 to S5), we find the following concentration model (where we have removed the tildes for clarity):

$$\frac{dS}{dt} = \mu.S - d_S.S - \frac{\epsilon_V.\phi_S}{Q_v}.S.V \tag{S9}$$

$$\frac{dI}{dt} = \frac{\epsilon_V.\phi_S}{Q_v}.S.V - d_S.I - \frac{1}{\tau}.I \tag{S10}$$

$$\frac{dV}{dt} = \frac{\beta}{\tau}.\frac{Q_v}{Q_p}I - d_V.V - \frac{\phi_S}{Q_p}.S.V \tag{S11}$$

$$J_{DON} = \gamma.(\frac{1}{\tau} - \frac{\beta}{\tau}.\frac{Q_v}{Q_p}).I \tag{S12}$$

$$J_{PON} = (1-\gamma).(\frac{1}{\tau} - \frac{\beta}{\tau}.\frac{Q_v}{Q_p}).I \tag{S13}$$

### Derivation of model equilibria

To suggest parameterization for biogeochemical models, we derive approximate equilibria of the $SIVZ$ and $SIVRZ$ models in two cases:

- the quadratic mortality of the virus is null: $d_{V2} = 0$
- the quadratic mortality of the virus is positive: $d_{V2} > 0$

In both cases, we consider a positive quadratic mortality for the zooplankton ($d_{Z2} > 0$).

#### $SIVZ$ model: $d_{V2} = 0$

- Coexistence:
  By assuming that the mortality of viruses due to its adsorption by the $S$ class is negligible compared to its linear mortality ($\phi_S S < d_1$), we identify the following equilibrium:

$$S^* = \frac{\frac{1}{\tau} + d_S + \frac{g_Z^2.\epsilon_Z}{d_{Z2}}.(\frac{a}{c} - d_z)}{\frac{\epsilon_V.\phi_S.\beta}{d_V.\tau.Qp} + \frac{g_Z^2.\epsilon_Z}{d_{Z2}}.(\frac{b}{c} - 1)} \tag{S14}$$

$$I^* = \frac{a - b.S^*}{c} \tag{S15}$$

$$V^* = \frac{\beta.Q_v.I^*}{d_V.\tau.Q_p} \tag{S16}$$

$$Z^* = \frac{\epsilon_Z.g_Z.(S^* + I^*) - d_Z}{d_{Z2}} \tag{S17}$$

  With:
  $a = \mu - d_S + g_Z.\frac{d_Z}{d_{Z2}}$; $b = \frac{1}{\tau_K.K} + \frac{g_Z^2.\epsilon_Z}{d_{Z2}}$ and $c = \frac{\epsilon_V.\phi_S.\beta}{d_V.\tau.Q_P} + \frac{g_Z^2.\epsilon_Z}{d_{Z2}}$

  We consider the assumption to be valid for $\frac{\phi_S.K}{Q_p} < 10.d_s$, assuming $K$ as a large upper limit for $S$.

- Alternate equilibria:
  In addition to the equilibrium where the phytoplankton, the virus and the grazer coexist, four other possible equilibria are possible for the system of equations of the $SIVZ$ model:

– $S^*, I^*, V^*, 0$:

$$S^* = \frac{(d_S + \frac{1}{\tau}).d_V.\tau.Q_v}{\phi.\beta} \tag{S18}$$

$$I^* = \frac{(\mu - d_S - \frac{S^*}{\tau_K.K}).d_V.\tau.Q_v}{\phi.\beta} \tag{S19}$$

$V^*$ is unchanged.

– $S^*, 0, 0, Z^*$:

$$S^* = \frac{\mu - d_S + g_Z\frac{d_Z}{d_{Z2}}}{\frac{1}{\tau_K.K} + \frac{\epsilon_Z.g_Z^2}{d_{Z2}}} \tag{S20}$$

$$Z^* = \frac{\epsilon_Z.g_Z.S^* - d_Z}{d_{Z2}} \tag{S21}$$

– $S^*, 0, 0, 0$:

$$S^* = (\mu - d_S).\tau_K.K \tag{S22}$$

– $0, 0, 0, 0$

#### $SIVZ$ model: $d_{V2} > 0$

- Coexistence:
In the case where $d_{V2} > 0$, the virus equilibrium concentration $V^*$ is solution to the following quadratic equation:

$$\frac{\beta}{\tau}.\frac{Q_v}{Q_p}.I^* - (d_V + \frac{\phi_S}{Q_p}.S^*).V^* - d_{V2}.V^{*2} = 0. \tag{S23}$$

Which gives:

$$V^* = \frac{-d_V - \frac{\phi_S}{Q_p}.S^* \pm \sqrt{(d_V + \frac{\phi_S}{Q_p}.S^*)^2 + 4.d_{V2}.\frac{\beta}{\tau}.\frac{Q_v}{Q_p}.I^*}}{2.d_{V2}} \tag{S24}$$

We then define two cases (the first case being more appropriate in terms of assumptions):

– Case 1: $(d_V + \frac{\phi_S}{Q_p}.S^*)^2 < 10.(4.d_{V2}.\frac{\beta}{\tau}.\frac{Q_v}{Q_p}.I^*)$

– Case 2: $(d_V + \frac{\phi_S}{Q_p}.S^*)^2 > 10.(4.d_{V2}.\frac{\beta}{\tau}.\frac{Q_v}{Q_p}.I^*)$

We first need an approximation of $S^*$ and $I^*$ to determine which case we are in depending on the parameters. To do so, we first approximate $S^*$ and $V^*$ using the equilibrium of the $SVZ$ model. Particularly, we find reasonable to estimate $S^*$ using the $SVZ$ model for latent periods inferior to 7 days:

$$S^*_{SVZ} = \frac{\mu - d_S + g.\frac{d_Z}{d_{Z2}} + \frac{\epsilon_V.\phi_S}{Q_p}.\frac{d_V}{d_{V2}}}{\frac{1}{\tau_K.K} + \frac{\epsilon_Z.g^2}{d_{Z2}} + \frac{\beta.\epsilon_V\phi_S^2}{2.d_{V2}.Q_p.Q_v}} \tag{S25}$$

$$V^*_{SVZ} = \frac{\beta.\epsilon_V\frac{\phi_S}{Qp}.S^*_{SVZ} - d_V}{d_{V2}} \tag{S26}$$

Then we find that $I^*$ is solution to a quadratic equation in the $SIVZ$ model:

$$I^* = \frac{-b_1 - \sqrt{b_1^2 - 4.a_1.c_1}}{2.a_1} \tag{S27}$$

With:
$a_1 = -\frac{\epsilon_Z g^2}{d_{Z2}}$; $b_1 = -d_S - \frac{1}{\tau} - \frac{\epsilon_Z g^2}{d_{Z2}}.S^* + \frac{g.d_Z}{d_{Z2}}$ and $c_1 = \frac{\phi_S}{Q_v}.S^*.V^*$

To estimate in which case we are in, for each parameter combination, we follow the procedure of case 2 (see below).

– Case 1:

In the case 1, we can approximate equation S24 to:

$$V^* = -\frac{d_V}{2.d_{V2}} - \frac{\phi_S}{2.d_{V2}.Q_p}.S + \sqrt{\frac{\beta.Q_v}{d_{V2}.\tau.Q_p}}.\sqrt{I} \tag{S28}$$

We note $a = \frac{\beta.Q_v}{d_{V2}.\tau.Q_p}$.
Then we inject equations S26 and S19 (unchanged $Z^*$) in the equation of the S class, giving:

$$S^* = c - b.\sqrt{I} - d.I \tag{S29}$$

With:
$b = \frac{1}{e}.\frac{\phi_S.\sqrt{a}}{Q_v}$; $c = \frac{1}{e}.(\mu - d_S + \frac{g.d_Z}{d_{Z2}})$; $d = \frac{1}{e}.\frac{\epsilon_Z.g^2}{d_{Z2}}$ and $e = \frac{1}{\tau_K.K} + \frac{\epsilon_Z.g_Z^2}{d_{Z2}} - \frac{\phi_S^2}{Q_v.Q_p.2.d_{V2}}$

In the term $e$, the term $f = \frac{\phi_S^2}{Q_v.Q_p.2.d_{V2}}$ acts as a quadratic source term for $S$ which is not the case in the equation. This term results from the sink of virus to the $I$ type. In practice, we find that setting $f = 0$ yields better results. Then we inject equation S29 in the equation of the I class. We find that $\sqrt{I}$ is a solution to the following quartic equation:

$$A.I^2 + B.I.\sqrt{I} + C.I + D.\sqrt{I} + E = 0 \tag{S30}$$

With:
$A = (1-d).\frac{\epsilon_Z.g^2}{d_{Z2}} + d^2.f$; $B = b.\frac{\epsilon_Z.g^2}{d_{Z2}} - d.\frac{\phi_S}{Q_v}.\sqrt{a} - 2.d.b.f$; $C = h - \frac{\phi_S}{Q_v}.\sqrt{a}.b - c.\frac{\epsilon_Z.g^2}{d_{Z2}} + (2.d.c - b^2).f + \frac{d_V}{2.d_{V2}}.d.\frac{\epsilon_V.\phi_S}{Q_v}$; $D = c.\frac{\phi_S}{Q_v}.\sqrt{a} + 2.c.b.f + \frac{d_V}{2.d_{V2}}.b.\frac{\epsilon_V.\phi_S}{Q_v}$ and $E = -c^2.f - \frac{d_V}{2.d_{V2}}.c.\frac{\epsilon_V.\phi_S}{Q_v}$

where $h = -d_S - \frac{1}{\tau} + g.\frac{d_Z}{d_{Z2}}$.

We solve equation S30 using the function *numpy.roots* in Python.

- Case 2: for $\tau < 7\ days$, we first estimate $S^*$, $V^*$ and then $I^*$ using equations S25 and S26. Then to refine the estimate we iterate a thousand times by updating $V^*$ and $I^*$ using equation S24 and S27, which empirically is found to be efficient in determining the feasibility of the coexistence equilibrium of the $SIVZ$ model. Then we calculate whether the current parameters correspond to cases 1 or 2.

- Alternate equilibria:
  - $S^*, I^*, V^*, 0$: same as the case of coexistence with all terms related to grazing by the zooplankton equal to 0.
  - $S^*, 0, 0, Z^*$: equation S20 and S21
  - $S^*, 0, 0, 0$: equation S22
  - $0, 0, 0, 0$

#### $SIVRZ$ model: $d_{V2} = 0$

- Coexistence:
For the $SIVRZ$ model we have the same assumption as for the $SIVZ$ model, and we neglect terms associated with mutations from S type to R type and reversely. First, $Z^*$ is changed to:

$$Z^* = \frac{\epsilon_Z.g.(S^* + I^* + R^*) - d_Z}{d_{Z2}} \tag{S31}$$

Then from equating the equations of $S$ and $R$ to 0 and subtracting one from the other, we find:

$$V^* = \frac{a}{b} \tag{S32}$$

And given the assumption of neglecting viral mortality due to adsorption:

$$I^* = \frac{a}{c} \tag{S33}$$

With:
$a = \mu.(\zeta - 1)$; $b = \frac{1}{Q_v}.(\epsilon_{VR}.\phi_R - \epsilon_V.\phi_S)$ and $c = \frac{\beta.Q_v}{\tau.d_v.Q_p}.b$;

Then from the equation of $R$, we find:

$$S^* + R^* = \frac{d}{e} \tag{S34}$$

With: $d = \mu.\zeta - d_S - (\frac{\epsilon_{VR}.\phi_R.\beta}{\tau.d_V.Q_P} + \frac{g_Z^2.\epsilon_Z}{d_{Z2}}).\frac{a}{b} + \frac{g_Z.d_Z}{d_{Z2}}$; $e = \frac{1}{\tau_K.K} + \frac{g_Z^2.\epsilon_Z}{d_{Z2}}$;

From the equation of $I$:

$$S^* + f.R^* = \frac{h}{i} \tag{S35}$$

With:

$f = \frac{\epsilon_{RV}.\phi_R}{\epsilon_V.\phi_S}$; $h = \frac{I^*}{\tau} + \frac{\epsilon_Z.g^2}{d_{Z2}}.(I^{*2} + \frac{c}{d}.I^*) - g.\frac{d_z}{d_{Z2}}.I^*$ and $i = V^*.\frac{\epsilon_V.\phi_S}{Q_v}$

From equation S34 and S35, we finally have:

$$R^* = \frac{\frac{h}{i} - \frac{d}{e}}{f - 1} \tag{S36}$$

$$S^* = \frac{d}{e} - R^* \tag{S37}$$

For this equilibrium, we empirically find that after first approximating $I^*$ to equation S33, we can update it to:

$$I^* = \frac{j}{k} \tag{S38}$$

With:

$j = d_V.V^* + (\frac{\phi_S}{Q_p}.S^* + \frac{\phi_R}{Q_p}.R^*).V^*$ and $k = \frac{\beta.Q_v}{\tau.Q_p}$

$S^*$ and $R^*$ are then updated again using equations S36 and S37 yielding a close to perfect theoretical equilibrium point and allowing to omit the assumption. However, for alternate equilibria, including the virus, the assumption still holds.

We note that the equation of $V^*$ imposes that $\zeta < 1$ (to satisfy $V_i^* > 0$) as by default we have $\phi_R < \phi_S$. Note also that this system of equations could also be solved differently and has other possible equilibrium points. Using simulations, we find that the equilibrium we wrote is the one that works for the parameter values we use.

- Alternate equilibria:
  In total, ten other equilibria are possible:
  - $S^*, I^*, V^*, \delta_{R^*}, Z^*$: same as $SIVZ$ model and considering $S^* >> R^*$:

$$\delta_{R^*} = -\frac{\mu.r.S^*}{\zeta.\mu - d_S - \frac{S^*}{\tau_K.K} - g_Z.Z^* - \frac{\phi_R.V^*}{Q_v} - \zeta.\mu.r} \tag{S39}$$

  - $\delta_{S^*}, I^*, V^*, R^*, Z^*$: same as previous case with $S^* << R^*$ (switching $S$ and $R$)
  - $S^*, I^*, V^*, R^*, 0$: same as the coexistence case with terms associated to grazing set to 0
  - $S^*, I^*, V^*, \delta_{R^*}, 0$: same as $SIVZ$ model (alternate equilibrium $S^*, I^*, V^*, 0$)
  - $\delta_{S^*}, I^*, V^*, R^*, 0$: same as previous case with $S^* << R^*$
  - $S^*, 0, 0, \delta_{R^*}, Z^*$: same as $SIVZ$ model (alternate equilibrium $S^*, 0, 0, Z^*$)
  - $\delta_{S^*}, 0, 0, R^*, Z^*$: same as previous case with $S^* << R^*$
  - $S^*, 0, 0, \delta_{R^*}, 0$: same as $SIVZ$ model (alternate equilibrium $S^*, 0, 0, 0$)
  - $\delta_{S^*}, 0, 0, R^*, 0$: same as previous case with $S^* << R^*$
  - $0, 0, 0, 0, 0$

#### $SIVRZ$ model: $d_{V2} > 0$

- Coexistence: In the case where $d_{V2} > 0$, the virus equilibrium concentration $V^*$ is solution to the following quadratic equation:

$$\frac{\beta}{\tau}.\frac{Q_v}{Q_p}.I^* - (d_V + \frac{\phi_S}{Q_p}.S^* + \frac{\phi_R}{Q_p}.R^*).V^* - d_{V2}.V^{*2} = 0 \tag{S40}$$

  Similarly to the $SIVZ$ model, we define two cases:
  - Case 1: $(d_V + \frac{\phi_S}{Q_p}.S^* + \frac{\phi_R}{Q_p}.R^*)^2 < 10.(4.d_{V2}.\frac{\beta}{\tau}.\frac{Q_v}{Q_p}.I^*)$
  - Case 2: $(d_V + \frac{\phi_S}{Q_p}.S^* + \frac{\phi_R}{Q_p}.R^*)^2 > 10.(4.d_{V2}.\frac{\beta}{\tau}.\frac{Q_v}{Q_p}.I^*)$

  Like for the $SIVZ$ model, we use approximation from the $SVRZ$ model to determine the case:

$$R^*_{SVRZ} = -\frac{C - C_1.\frac{A}{A_1}}{B - B_1.\frac{A}{A_1}} \tag{S41}$$

$$S^*_{SVRZ} = \frac{-B_1.R^*_{SVRZ} - C_1}{A_1} \tag{S42}$$

  With:

$A = -\frac{\epsilon_V.\phi_S}{Q_v}.a - \frac{1}{\tau_K.K} - \frac{g^2.\epsilon_Z}{d_{Z2}}$; $B = -\frac{\epsilon_V.\phi_S}{Q_v}.b - \frac{1}{\tau_K.K} - \frac{g^2.\epsilon_Z}{d_{Z2}}$ and $C = \mu - d_S - \frac{\epsilon_V.\phi_S}{Qv}.c + \frac{g.d_Z}{d_{Z2}}$

Symmetrically:

$A_1 = -\frac{\epsilon_{VR}.\phi_R}{Q_v}.a - \frac{1}{\tau_K.K} - \frac{g^2.\epsilon_Z}{d_{Z2}}$; $B_1 = -\frac{\epsilon_{VR}.\phi_R}{Q_v}.b - \frac{1}{\tau_K.K} - \frac{g^2.\epsilon_Z}{d_{Z2}}$ and $C_1 = \zeta.\mu - d_S - \frac{\epsilon_{VR}.\phi_R}{Qv}.c + \frac{g.d_Z}{d_{Z2}}$

And with:

$a = \frac{\phi_S}{Q_p.d_{V2}}.(\epsilon_V.\beta - 1)$; $b = \frac{\phi_R}{Q_p.d_{V2}}.(\epsilon_{VR}.\beta - 1)$ and $c = -\frac{d_V}{d_{V2}}$

Like for the $SIVZ$ model, we then estimate $I^*$ by modifying equation S27:

$$I^* = \frac{-b_2 - \sqrt{b_2^2 - 4.a_2.c_2}}{2.a_2} \tag{S43}$$

With:

$a_2 = -\frac{\epsilon_Z g^2}{d_{Z2}}$; $b_2 = -d_S - \frac{1}{\tau} - \frac{\epsilon_Z g^2}{d_{Z2}}.(S^* + R^*) + \frac{g.d_Z}{d_{Z2}}$ and $c_2 = (\frac{\phi_S}{Q_v}.S^* + \frac{\phi_R}{Q_v}.R^*).V^*$

We then follow the procedure described for case 2 (see below) to determine which case we are in.

– Case 1:

Like in the case of the $SIVZ$ model where $d_{V2} > 0$, we find that $V^*$ follows equation S38. Then, given that we are in case 1, we first make the approximation:

$$V^* = \sqrt{\frac{\beta.Q_v}{d_{V2}.\tau.Q_p}}.\sqrt{I} \tag{S44}$$

Yielding:

$$I^* = \left(\frac{a}{b}\right)^2 \tag{S45}$$

With:

$a = \mu.(\zeta - 1)$ and $b = \sqrt{\frac{\beta.Q_v}{\tau.Q_p.d_{V2}}}.(\epsilon_{VR}.\phi_R - \epsilon_V.\phi_S).\frac{1}{Q_v}$

Then using the equation of $R$ and of $I$, $R^*$ and $S^*$ follow the modified equations S36 and S37 by modifying $I^*$ in the parameters using equation S38:

$$R^* = \frac{\frac{h}{i} - \frac{d}{e}}{f - 1} \tag{S46}$$

$$S^* = \frac{d}{e} - R^* \tag{S47}$$

For this equilibrium, we also empirically find that after first approximating $I^*$ to equation S45, we can update it to:

$$I^* = \frac{j}{k} \tag{S48}$$

With:

$j = d_V.V^* + d_{V2}.V^{*2} + (\frac{\phi_S}{Q_p}.S^* + \frac{\phi_R}{Q_p}.R^*).V^*$ and $k = \frac{\beta.Q_v}{\tau.Q_p}$

$S^*$ and $R^*$ are then updated again using equations S46 and S47, generating a theoretical equilibrium close to perfect.

– Case 2: for $\tau < 7\ days$, we first keep the approximations $S^*_{SVRZ} = S^*_{SIVRZ}$ and $R^*_{SVRZ} = R^*_{SIVRZ}$. Then, we first approximate $I^*$ using equation S43 with $V^*$ following equation S40. Then, we iteratively update $I^*$, $R^*$ and $S^*$ a thousand times using equations S42, S47 and S45. Finally, we determine in which case we are in and keep the final approximation if we are in case 2.

- Alternate equilibria: same as above (see case $d_{V2} = 0$) and referring to the $SIVZ$ model with $d_{V2} > 0$ when $V$ is present and $R$ or $S$ excluded.

| Parameter | Unit | ***Prochlorococcus*** | ***Synechococcus*** | **Picoeukaryote** | **Diatom** |
|---|---|---|---|---|---|
| Maximum growth rate | $d^{-1}$ | 0.75 | 0.87 | 1.25 | 2.66 |
| Cell radius | $\mu m$ | 0.3 | 0.55 | 1 | 3 |
| $Q_p$ | $\mu molN.ind^{-1}$ | $6.1.10^{-10}$ | $4.10^{-9}$ | $1.10^{-8}$ | $1.7.10^{-7}$ |
| Virus radius | $nm$ | 35 | 35 | 80 | 20 |
| Burst size | $\#\ of\ infective\ virions$ | 15 | 30 | 180 | 270 |
| Latent period | $d$ | 0.37 | 0.37 | 0.37 | 0.95 |
| $Q_v$ | $\mu molN.ind^{-1}$ | $1.4.10^{-12}$ | $1.4.10^{-12}$ | $1.5.10^{-11}$ | $3.10^{-13}$ |
| $Q_z$ | $\mu molN.ind^{-1}$ | $1.6.10^{-7}$ | $1.6.10^{-7}$ | $10^{-6}$ | $10^{-6}$ |

**Table S1 | Parameters associated with the growth of the four phytoplankton types, their respective viruses and zooplankton.** Maximum growth rates are defined from allometric laws from Dutkiewicz et al. (2020). Cellular quota for the cyanobacteria are derived from Verity et al. (1992) and from Menden-Deuer and Lessard (2000) for the picoeukaryote (non-diatom), the small diatom and the zooplankton. Virion quotas are derived from Jover et al. (2014). Latent period and burst size are based on our life history trait model which takes the host volume, the virus radius, and the virus-host type pair as predictors.

| Parameter | Symbol | Unit | Value or Range | Reference |
|---|---|---|---|---|
| Linear mortality of the phytoplankton | $d_S$ | $d^{-1}$ | 0.1 | Dutkiewicz et al. 2020 |
| Carrying capacity (Mesotrophic) | $K$ | $\mu molN.L^{-1}$ | 0.76 | see Table S3 |
| Relaxation time | $\tau_K$ | $d$ | $\frac{1}{\mu - d}$ | - |
| Adsorption rate | $\phi_S$ | $L.d^{-1}$ | $10^{-12} - 10^{-7}$ | Talmy et al. 2019b |
| Adsorption rate of the resistant type | $\phi_R$ | $L.d^{-1}$ | $0 - 7.7.10^{-8}$ | Talmy et al. 2019b |
| Adsorption efficiency | $\epsilon_V$ | $fraction$ | $0 - 1$ | - |
| Linear mortality of the virus | $d_V$ | $d^{-1}$ | 0.1 | - |
| Quadratic mortality of the virus | $d_{V2}$ | $(\mu molN.L^{-1})^{-1}.d^{-1}$ | 1800 | Beckett et al. 2024 |
| Latent period | $\tau$ | $d$ | $0.1 - 90$ | - |
| Mutation rate | $r$ | $fraction$ | $10^{-6}$ | - |
| Growth penalty of the resistant type | $\zeta$ | $fraction$ | 0.8 | - |
| Grazing rate | $g_Z$ | $(\mu molN.L^{-1})^{-1}.d^{-1}$ | 9.8 | Dutkiewicz et al. 2020 |
| Grazing efficiency | $\epsilon_Z$ | $fraction$ | 0.3 | Straile 1997 |
| Linear mortality of the zooplankton | $d_Z$ | $d^{-1}$ | 0.067 | Dutkiewicz et al. 2020 |
| Quadratic mortality of the zooplankton | $d_{Z2}$ | $(\mu molN.L^{-1})^{-1}.d^{-1}$ | 1.4 | Dutkiewicz et al. 2015b |

**Table S2 | Default model parameters without environmental effects.**

| Parameter | Formula | Unit | Oligotrophic | Mesotrophic |
|---|---|---|---|---|
| Temperature | $T$ | $°C$ | 25 | 20 |
| Temperature limitation | $\tau_T.e^{-A_T.(\frac{1}{T}-\frac{1}{T_N})}$ | $fraction$ | 1 | 0.8 |
| Nutrient concentration | $N$ | $\mu molN.L^{-1}$ | 0.1 | 1 |
| Surface-deep mixing rate | $w$ | $d^{-1}$ | 0.01 | 0.05 |
| Deep inorganic N | $N_{deep}$ | $\mu molN.L^{-1}$ | 10 | 10 |
| Carrying capacity | $K = -w.(N - N_{deep}).\frac{N+N_c}{\mu.N}$ | $\mu molN.L^{-1}$ | 0.15 | 0.76 |
| Nutrient limitation | $\frac{N}{N+Nc}$ | $fraction$ | 0.96 | 0.99 |

**Table S3 | Factors modulating model parameters to represent different ocean environments for *Prochlorococcus*.**

| Phytoplankton | | | Oligotrophic | Mesotrophic | Reference |
|---|---|---|---|---|---|
| Small *Prochlorococcus* | Tot P | Range | $10^7 - 10^9$ | $10^7 - 10^9$ | Mojica et al. 2016<br>Carlson et al. 2022<br>Beckett et al. 2024<br>Schartau et al. 2010 |
| | | Target | $1.5.10^8$ | $2.10^8$ | |
| | V | Range | $10^7 - 10^{10}$ | $10^7 - 10^{10}$ | |
| | | Target | $5.10^8$ | $10^9$ | |
| | Z | Range | $10^3 - 10^6$ | $10^3 - 10^6$ | |
| | | Target | $10^5$ | $2.10^5$ | |
| | % Inf | Range | 0 – 5 | 0.5 – 10 | |
| | % V kill | Range | 0 – 50 | 0 – 50 | |
| Small *Synechococcus* | Tot P | Range | $10^7 - 10^9$ | $10^7 - 10^9$ | Mojica et al. 2016<br>Carlson et al. 2022<br>Beckett et al. 2024<br>Schartau et al. 2010 |
| | | Target | $2.10^7$ | $5.10^7$ | |
| | V | Range | $10^7 - 10^{10}$ | $10^7 - 10^{10}$ | |
| | | Target | $5.10^8$ | $10^9$ | |
| | Z | Range | $10^3 - 10^6$ | $10^3 - 10^6$ | |
| | | Target | $10^5$ | $2.10^5$ | |
| | % Inf | Range | 0 – 5 | 0.5 – 10 | |
| | % V kill | Range | 0 – 50 | 0 – 50 | |
| Other eukaryote | Tot P | Range | $10^6 - 10^8$ | $10^6 - 10^8$ | Mojica et al. 2016<br>Schartau et al. 2010 |
| | | Target | $10^7$ | $2.10^7$ | |
| | V | Range | $10^7 - 10^9$ | $10^7 - 10^9$ | |
| | | Target | $10^8$ | $2.10^8$ | |
| | Z | Range | $10^3 - 10^6$ | $10^3 - 10^6$ | |
| | | Target | $10^4$ | $5.10^4$ | |
| | % Inf | Range | 0 – 5 | 0.5 – 10 | |
| | % V kill | Range | 0 – 50 | 0 – 50 | |
| Small Diatom | Tot P | Range | $10^5 - 5.10^7$ | $10^5 - 5.10^7$ | Mojica et al. 2016<br>Leblanc et al. 2018<br>Tomaru et al. 2021<br>Arsenieff et al. 2019<br>Schartau et al. 2010 |
| | | Target | $10^6$ | $2.10^6$ | |
| | V | Range | $10^8 - 10^{10}$ | $10^8 - 10^{10}$ | |
| | | Target | $6.10^8$ | $10^9$ | |
| | Z | Range | $10^3 - 10^6$ | $10^3 - 10^6$ | |
| | | Target | $10^4$ | $5.10^4$ | |
| | % Inf | Range | 0.5 – 50 | 0.5 – 50 | |
| | % V kill | Range | 0 – 50 | 0 – 50 | |

**Table S4 | Ranges and target concentrations for the four phytoplankton types for the two epipelagic ocean types.** Concentrations are in $ind.L^{-1}$. Tot P: total phytoplankton (Susceptible + Infected + Resistant). % Inf: percentage of infected cells. % V kill: percentage of virus induced mortality.

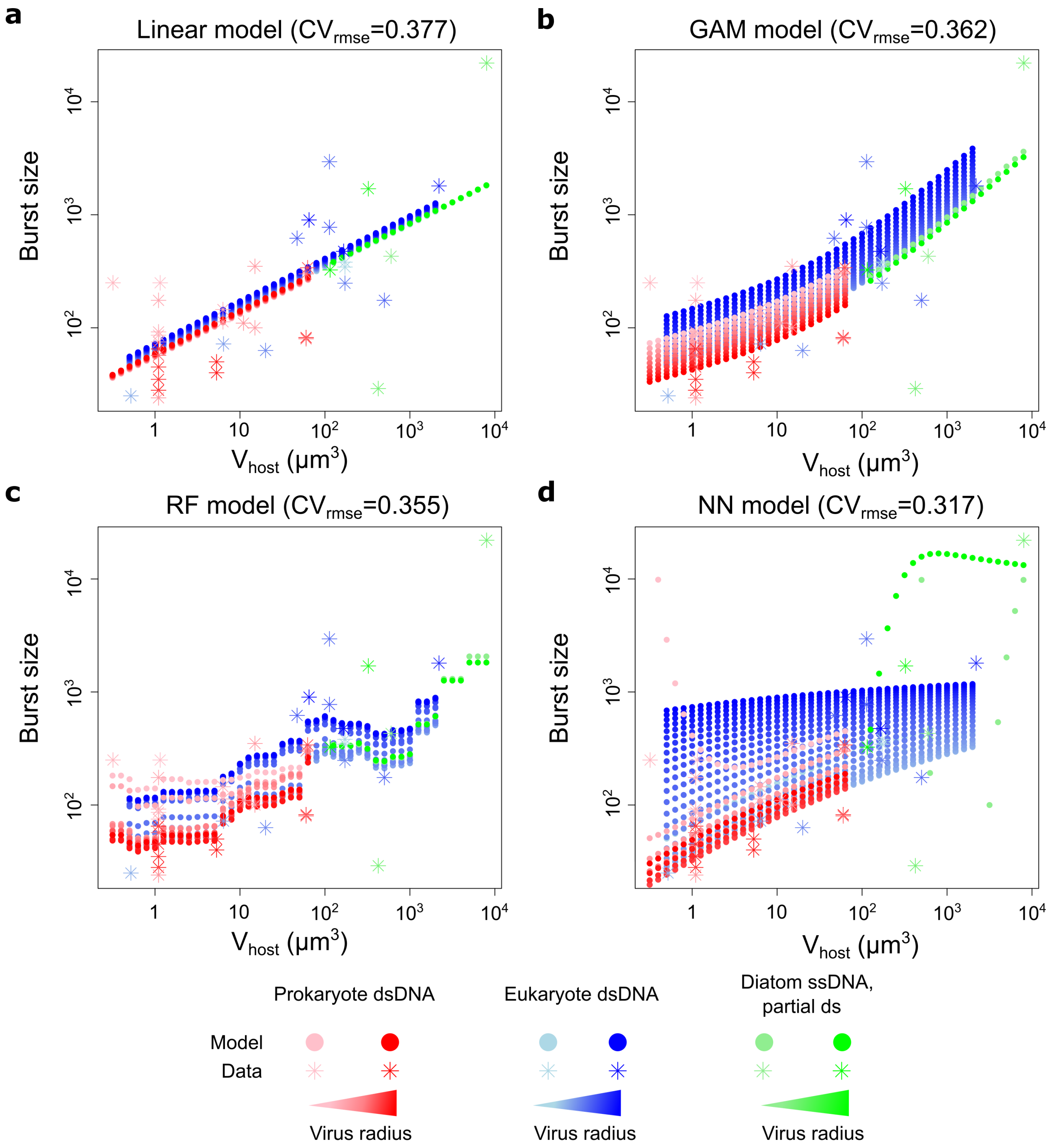


**Figure S1 | Model of burst size as a function of host volume and virion radius for 4 types of models, optimized using leave one out cross validation.** (**a**) Linear model, (**b**) Generalized additive model, (**c**) Random forest model, (**d**) Single layer neural network model. The cross validation root mean square error (rmse) evaluates the performance of the model. We fitted $log_{10}$ values so that the model error (rmse) is in $log_{10}$ scale. Data from Edwards and Steward (2018).

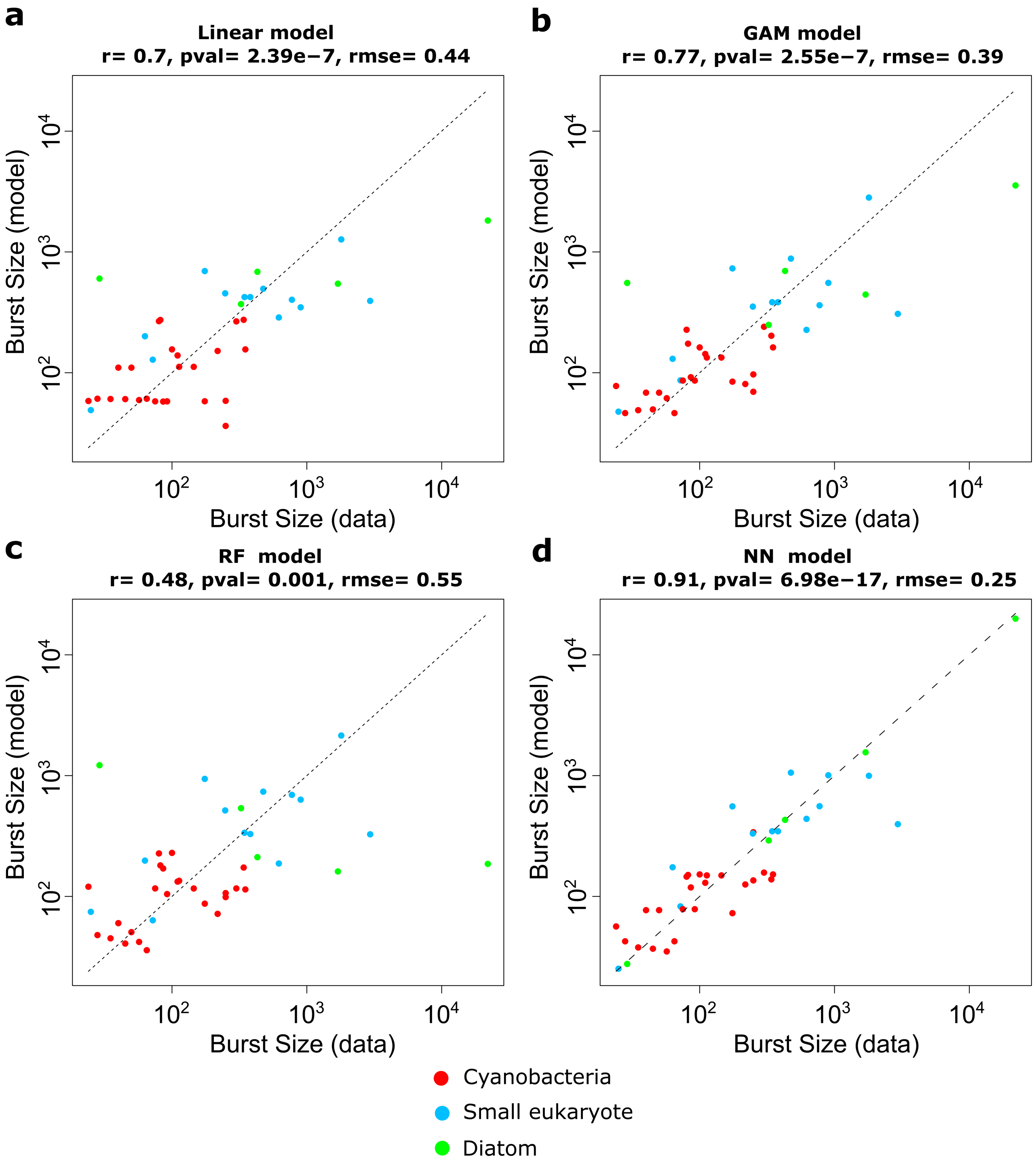


**Figure S2 | Model versus data of burst size for 4 types of models, optimized using leave one out cross validation.** (**a**) Linear model, (**b**) Generalized additive model, (**c**) Random forest model, (**d**) Single layer neural network model. The root mean square error (rmse) is in $log_{10}$ scale. Data from Edwards and Steward (2018).

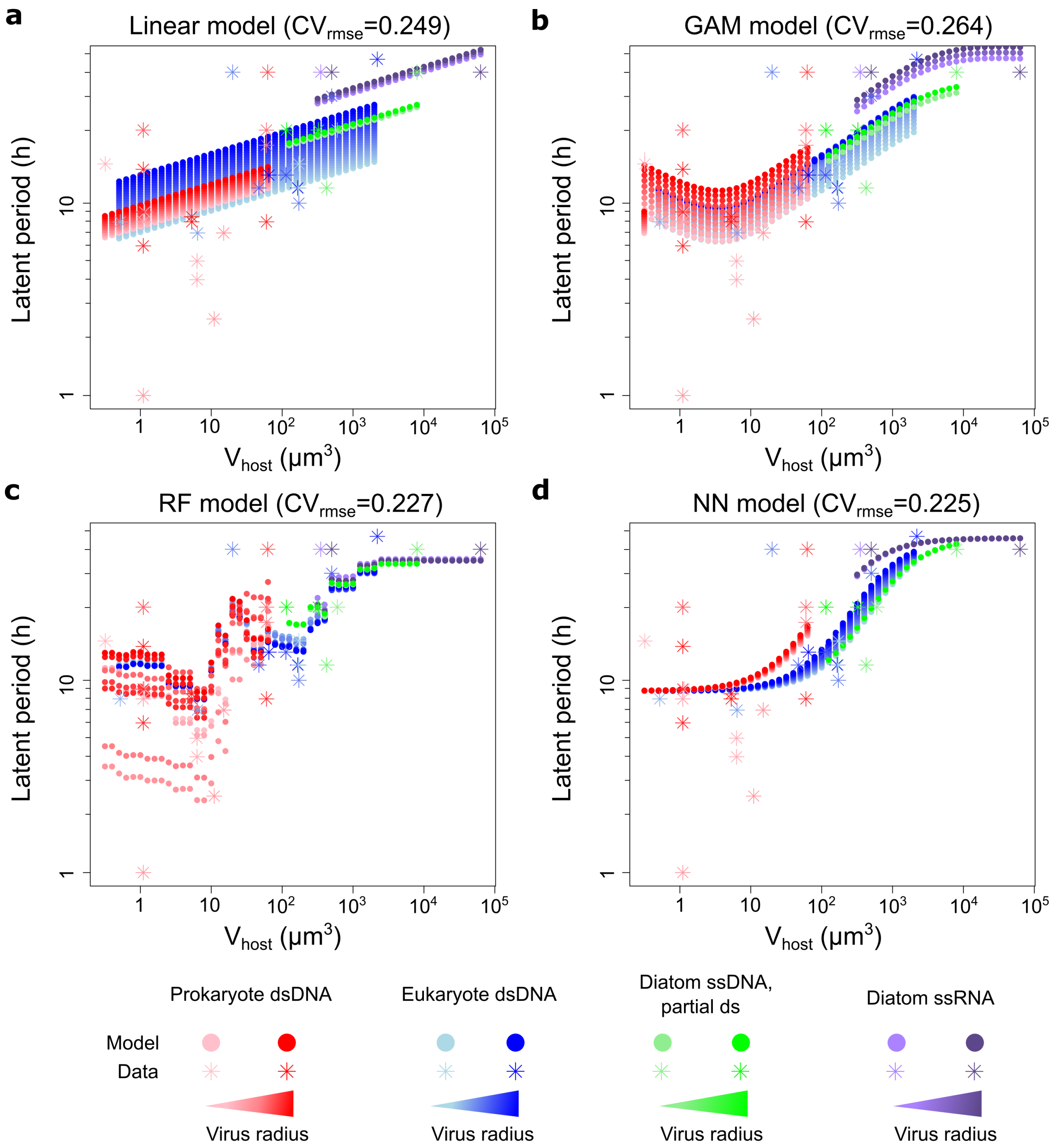


**Figure S3 | Model of latent period as a function of host volume and virion radius for 4 types of models, optimized using leave one out cross validation.** (**a**) Linear model, (**b**) Generalized additive model, (**c**) Random forest model, (**d**) Single layer neural network model. The cross validation root mean square error (rmse) evaluates the performance of the model. We fitted $log_{10}$ values so that the model error (rmse) is in $log_{10}$ scale. Data from Edwards and Steward (2018).

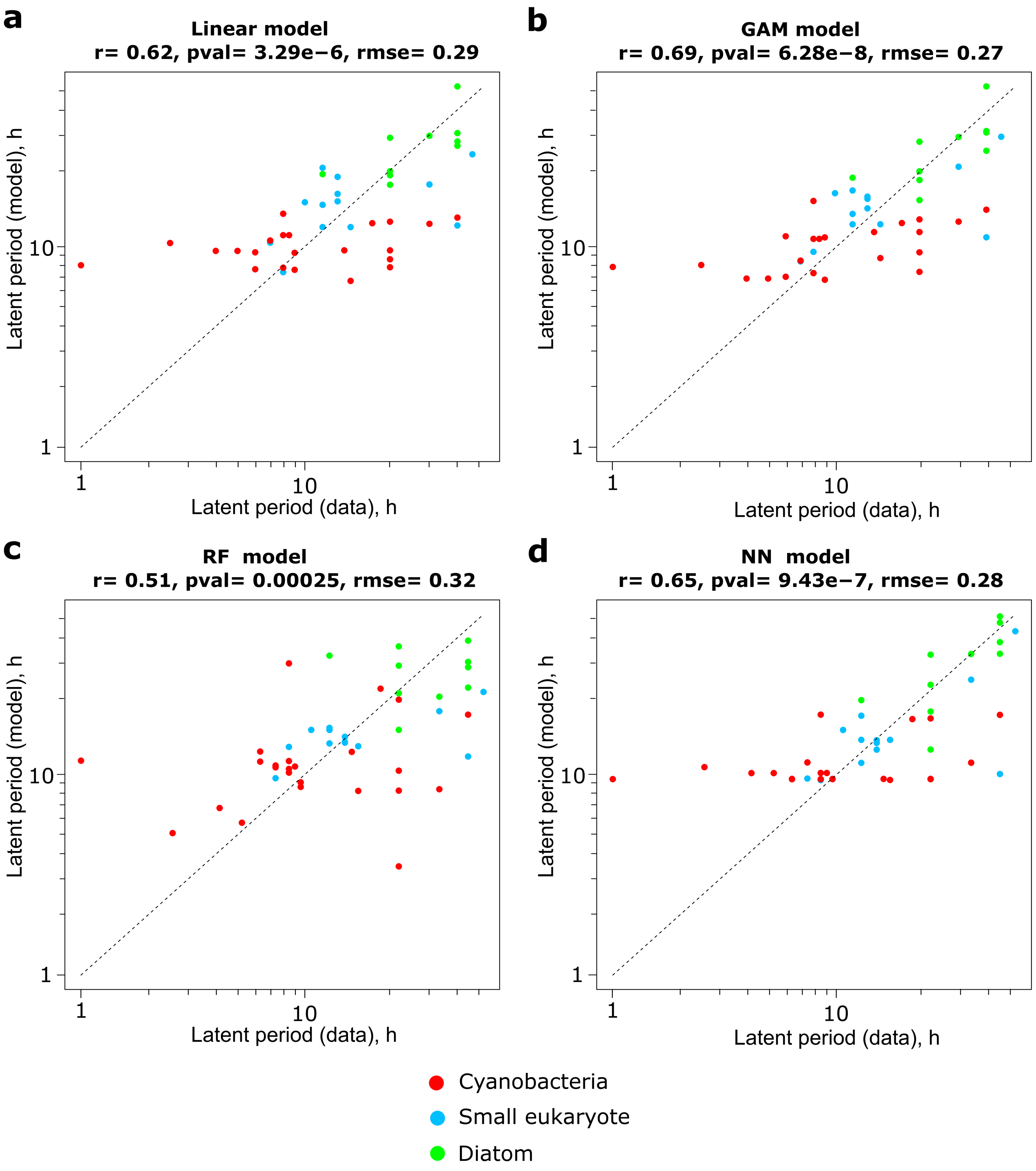


**Figure S4 | Model versus data of latent period for 4 types of models, optimized using leave one out cross validation.** (**a**) Linear model, (**b**) Generalized additive model, (**c**) Random forest model, (**d**) Single layer neural network model. The root mean square error (rmse) is in $log_{10}$ scale. Data from Edwards and Steward (2018).

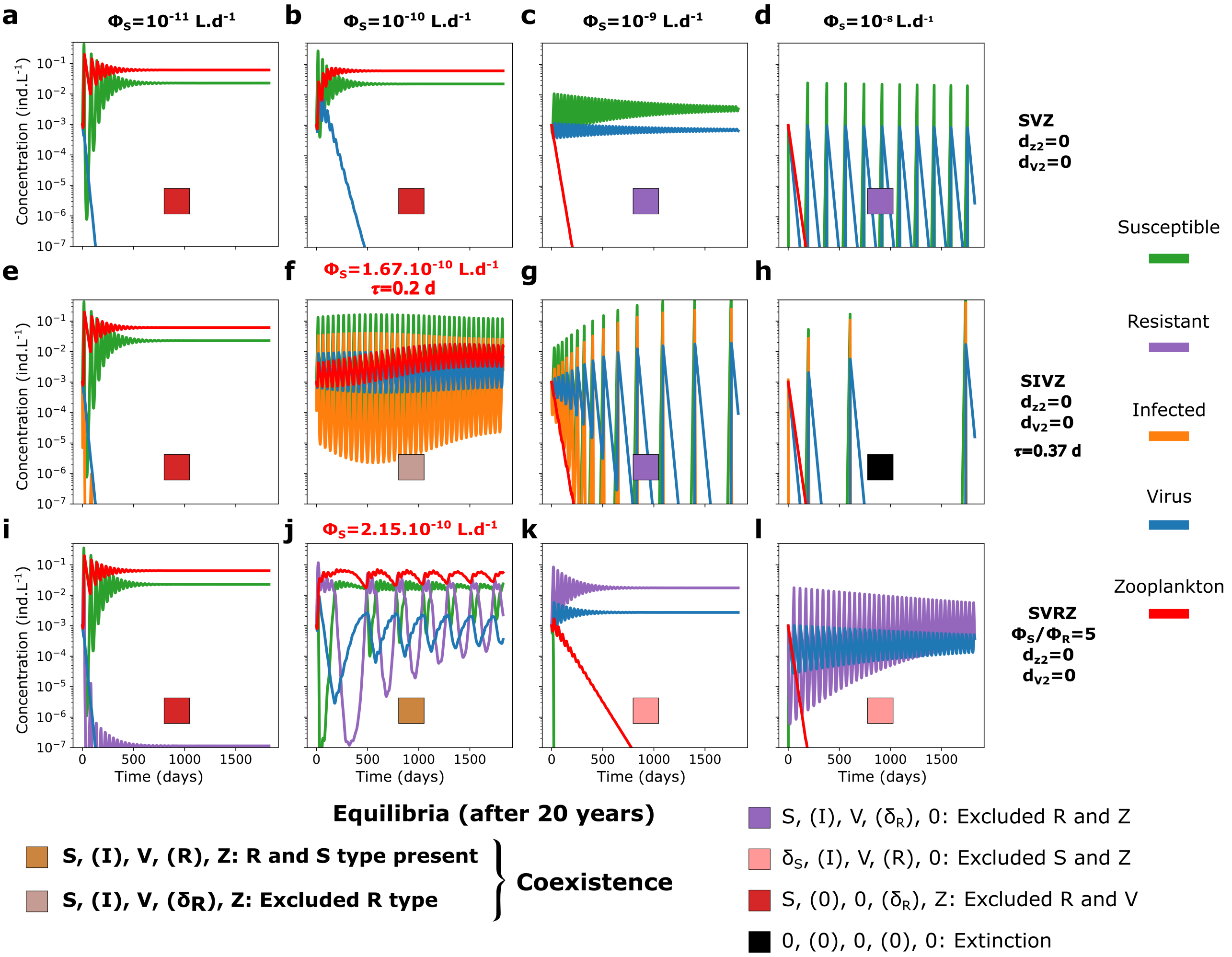


**Figure S5 | Model output for five year time series in nitrogen molar concentrations of the $SVZ$, $SIVZ$ and the $SVRZ$ models for *Prochlorococcus* for four different adsorption rates of the virus without quadratic mortality terms.** Adsorption rate from $\phi_S = 10^{-11}\ L.d^{-1}$ to $\phi_S = 10^{-8}\ L.d^{-1}$ ($\epsilon_V = 1$) for $SVZ$ (**a-d**) the $SVZ$ model, (**e-h**) the $SIVZ$ model and (**i-l**) the $SVRZ$ model. A latent period of 0.37 *days* and a burst size of 15 were used as parameterized by the life history trait model (see Methods). To showscase the coexistence regime facilitated by the $I$ class (Infected) and $R$ (Resistant), we changed the parameters slightly (**g**) $\tau = 2.15d$ and (**j**) $\phi_S = 1.67.10^{-10}$ (highlighted in red).

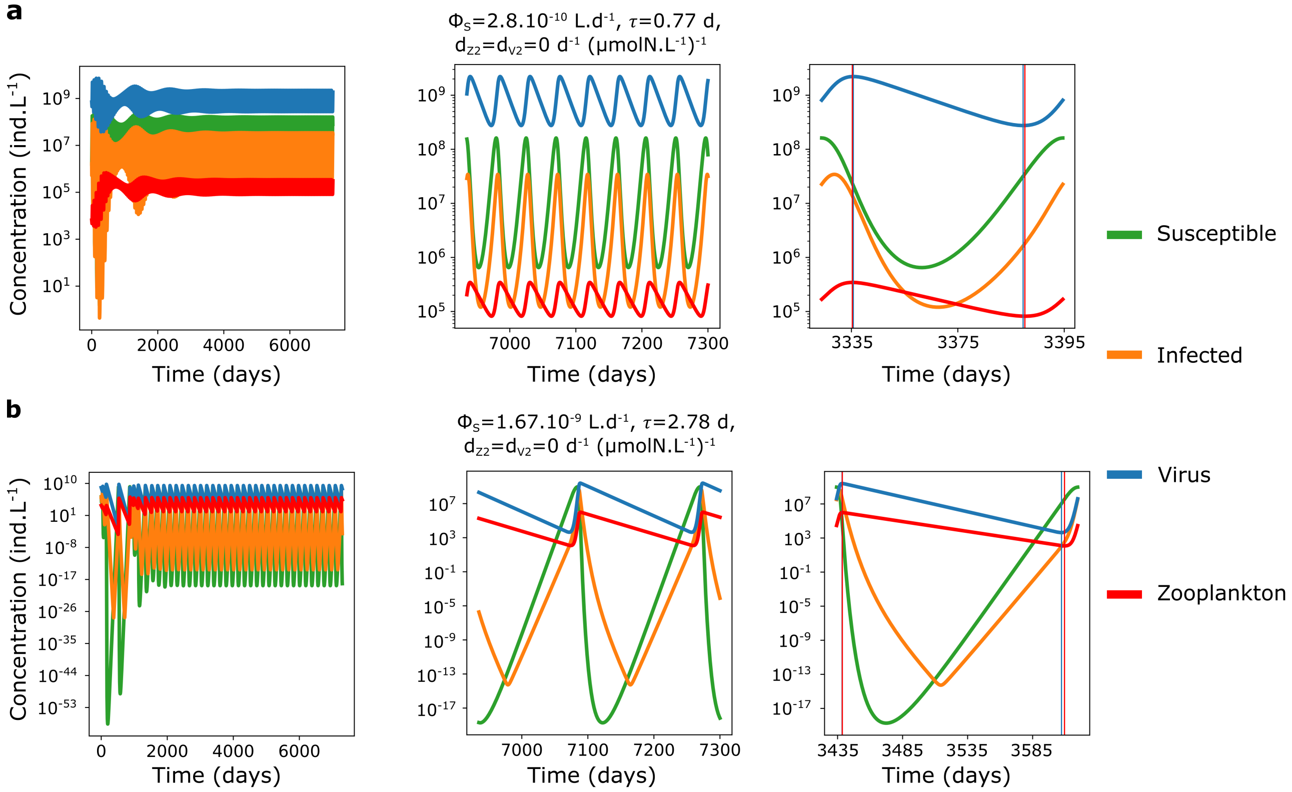


**Figure S6 | Model output of time series of two example coexistence regime between the virus and the zooplankton in the $SIVZ$ model without quadratic mortality terms of the predators.** (**a**) $\phi_S = 1.7.10^{-10}\ L.d^{-1}$ and $\tau = 0.28\ d$, (**b**) $\phi_S = 1.3.10^{-9}\ L.d^{-1}$ and $\tau = 7.7\ d$. Left panels: 20 years of simulations. Middle panels: Last year of simulation. Right panels: one period. In the right panels, vertical lines indicate the peaks (highs and lows) of the virus (blue) and the zooplankton (red).

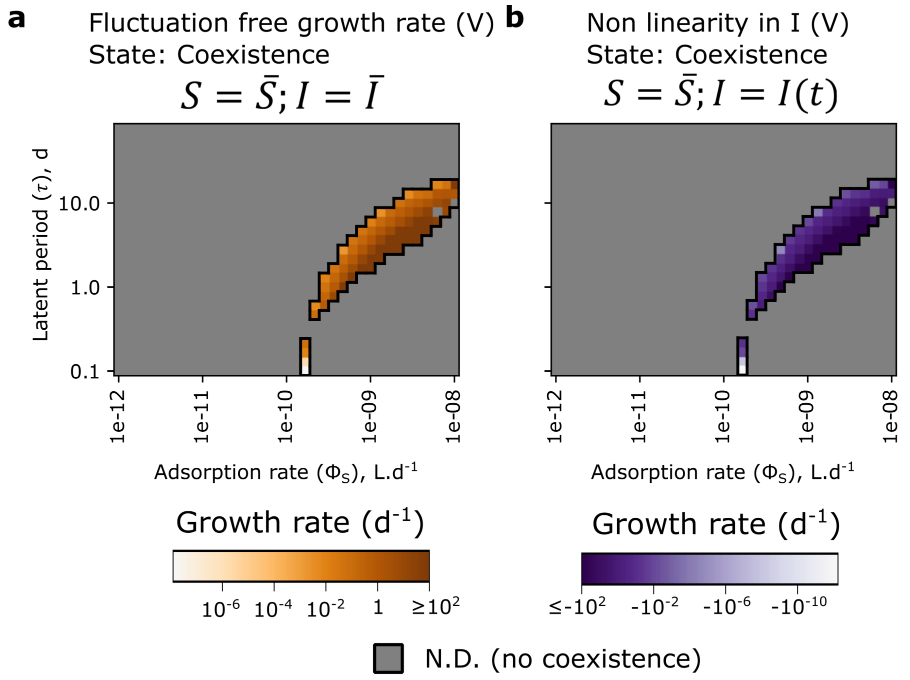


**Figure S7** | **Growth rate analysis of the free virus in the coexistence regime** (**a**) Fluctuation free growth rate of the free virus. (**b**) Non linearity in $I$ component of the growth rate of the free virus: the growth rate of the free virus is decreased due to the nonlinear oscillations of the $I$ class. For each panel, the coexistence regime reached by the $SIVZ$ model after 20 years of simulations, without quadratic mortality terms for the predators, is represented as the black contour. N.D.: Not defined.

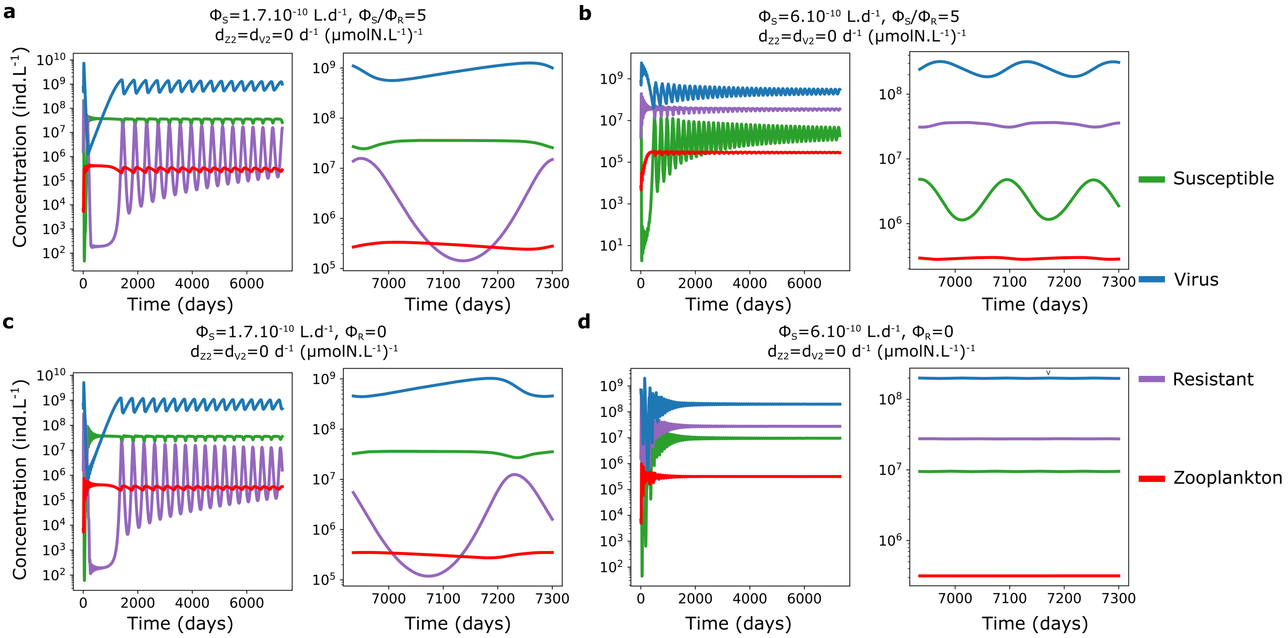


**Figure S8 | Model output of time series of four example coexistence regime between the virus and the zooplankton in the *SVRZ* model without quadratic mortality terms of the predators.** Time series for a partially resistant type, $\frac{\epsilon_V}{\epsilon_{VR}} = 10$, for (**a**) $\phi_S = 10^{-9}\ L.d^{-1}$ and (**b**) $\phi_S = 1.67.10^{-9}\ L.d^{-1}$ and for a fully resistant type, $\epsilon_{VR} = 0$, for (**c**) $\phi_S = 10^{-9}\ L.d^{-1}$ and (**d**) $\phi_S = 1.67.10^{-9}\ L.d^{-1}$. Left panels: 20 years of simulations. Middle panels: Last year of simulation.

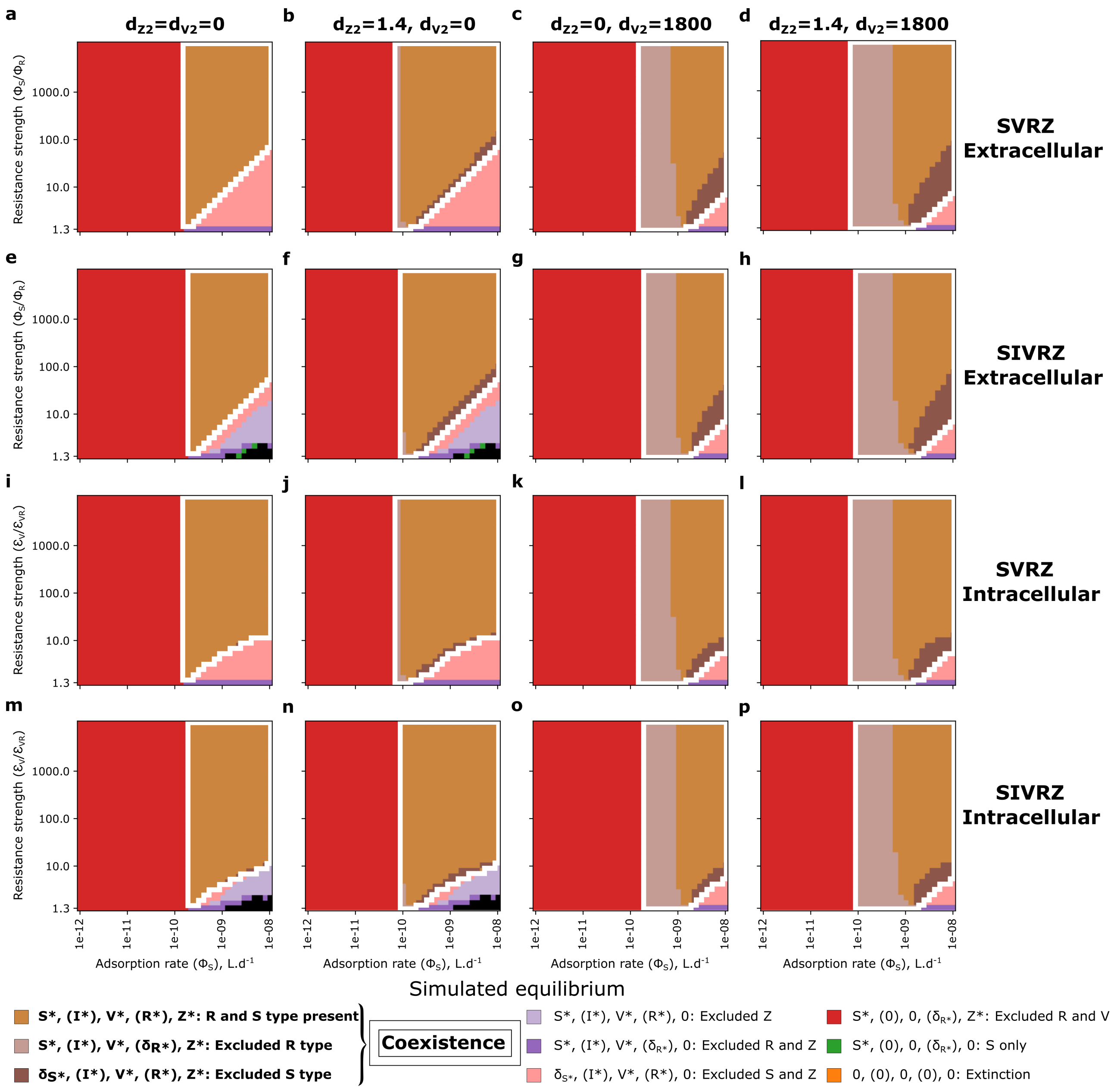


**Figure S9 | Equilibrium regimes of a virus and a zooplankton modeled as predators of a *Prochlorococcus* for different types of predation models across the adsorption rate and resistant strength parameter space.** (**a-d**) $SVRZ$ model with an extracellular resistant cell. (**e-h**) $SIVRZ$ model with an extracellular resistant cell. (**i-l**) $SVRZ$ model with an intracellular resistant cell. (**m-p**) $SIVRZ$ model with an intracellular resistant cell. From left to right panels: quadratic mortality terms respectively absent, present for the zooplankton only, present for the virus only, and present for both the zooplankton and the virus. Each model was run for 20 years. For each panel, the white contour denotes the coexistence regime. The quadratic mortality terms are in $(\mu mol N.L^{-1})^{-1}.d^{-1}$.

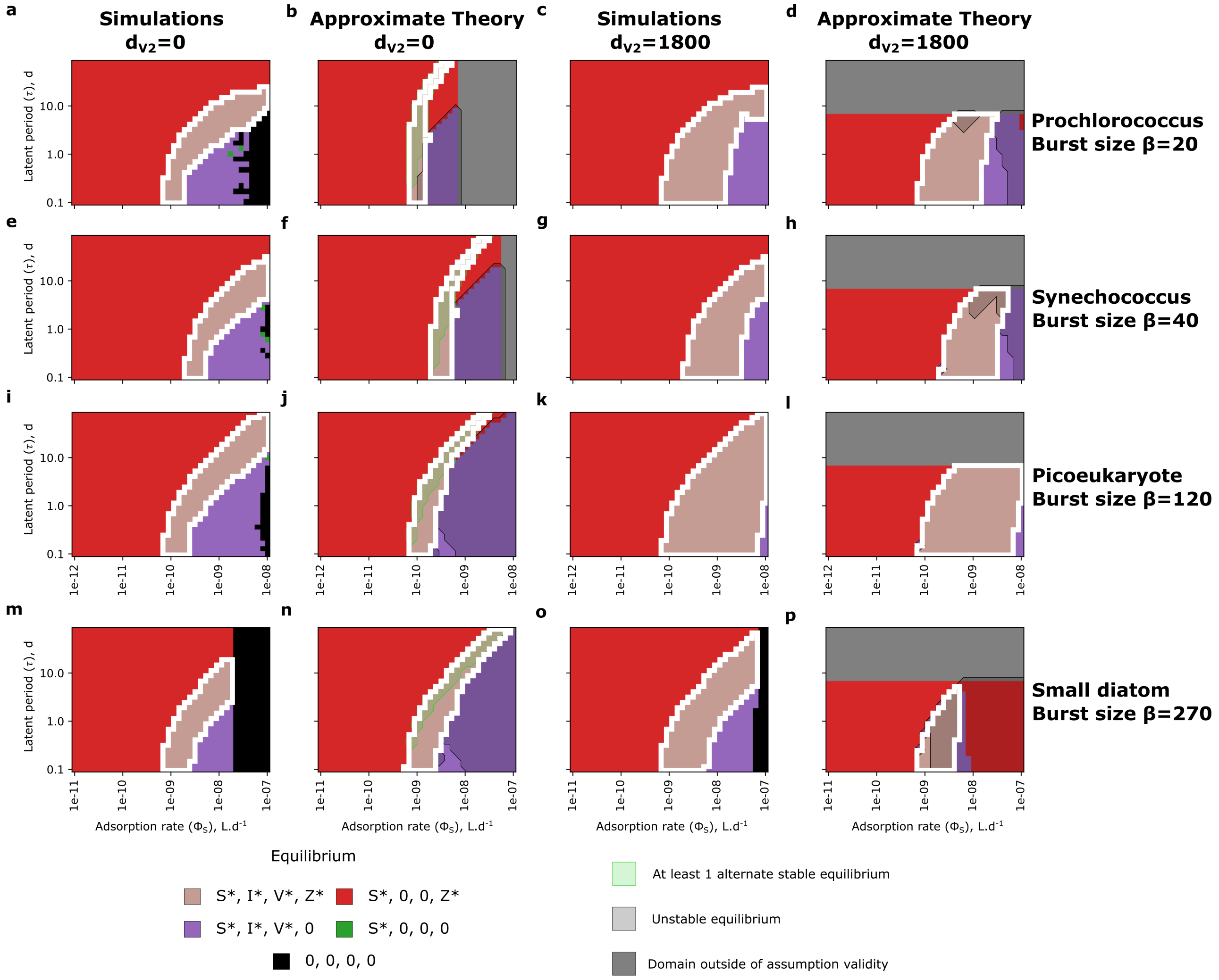


**Figure S10 | Simulated and approximate theoretical equilibrium regimes of the $SIVZ$ model for four different phytoplankton with different molar quotas and burst sizes across the spectrum of latent period and adsorption rate of the virus.** (**a-d**) *Prochlorococcus.* (**e-h**) *Synechococcus.* (**i-l**) Picoeukaryote (non-diatom). (**m-p**) Small diatom. When multiple stable equilibria exist (shaded green area), the most complex, *i.e.* the one with the most non-null tracers, is displayed. Note the different range of adsorption rate of the virus for the small diatom. Two first left panels: quadratic mortality term present for the zooplankton only. Right panels: quadratic mortality terms present for both the zooplankton and the virus. Note that panel (**a**) and (**c**) are also shown in Figure 2. Each model was run for 20 years. For each panel, the white contour denotes the coexistence regime. For the approximate theory, when at least 1 alternate stable equilibria exist, we represent the equilibrium with the largest number of non-null tracers and highlight the zone in transparent green. Unstable equilibria areas are colored in transparent gray. The quadratic mortality terms are in $(\mu mol N.L^{-1})^{-1}.d^{-1}$.

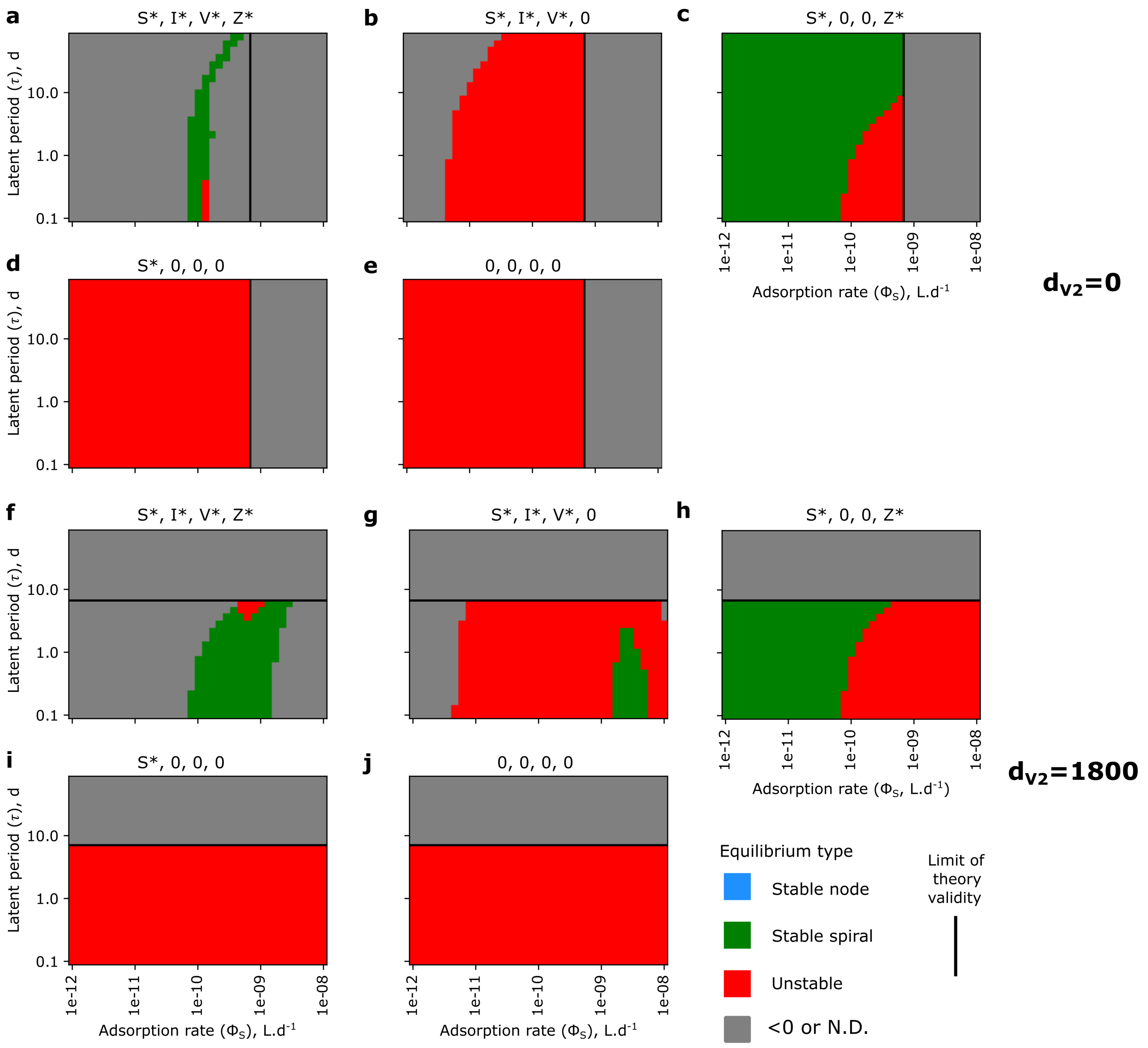


**Figure S11 | Equilibrium types for the five different possible equilibria of the $SIVZ$ model in the parameter space for *Prochlorococcus*.** (**a**, **f**) $S^*, I^*, V^*, Z^*$; (**b**, **g**) $S^*, I^*, V^*, 0$; (**c**, **h**) $S^*, 0, 0, Z^*$; (**d**, **i**) $S^*, 0, 0, 0$ and (**e**, **j**) $0, 0, 0, 0$. Top panels: quadratic mortality term present for the zooplankton only. Bottom panels: quadratic mortality terms present for both the zooplankton and the virus. Regions in gray in the limit of theory validity (left and below of the black bar for top panels and bottom panels respectively) are unfeasible equilibrium: at least one of the $X^*$ is negative. N.D.: Not Defined: outside of limit of theory validity.

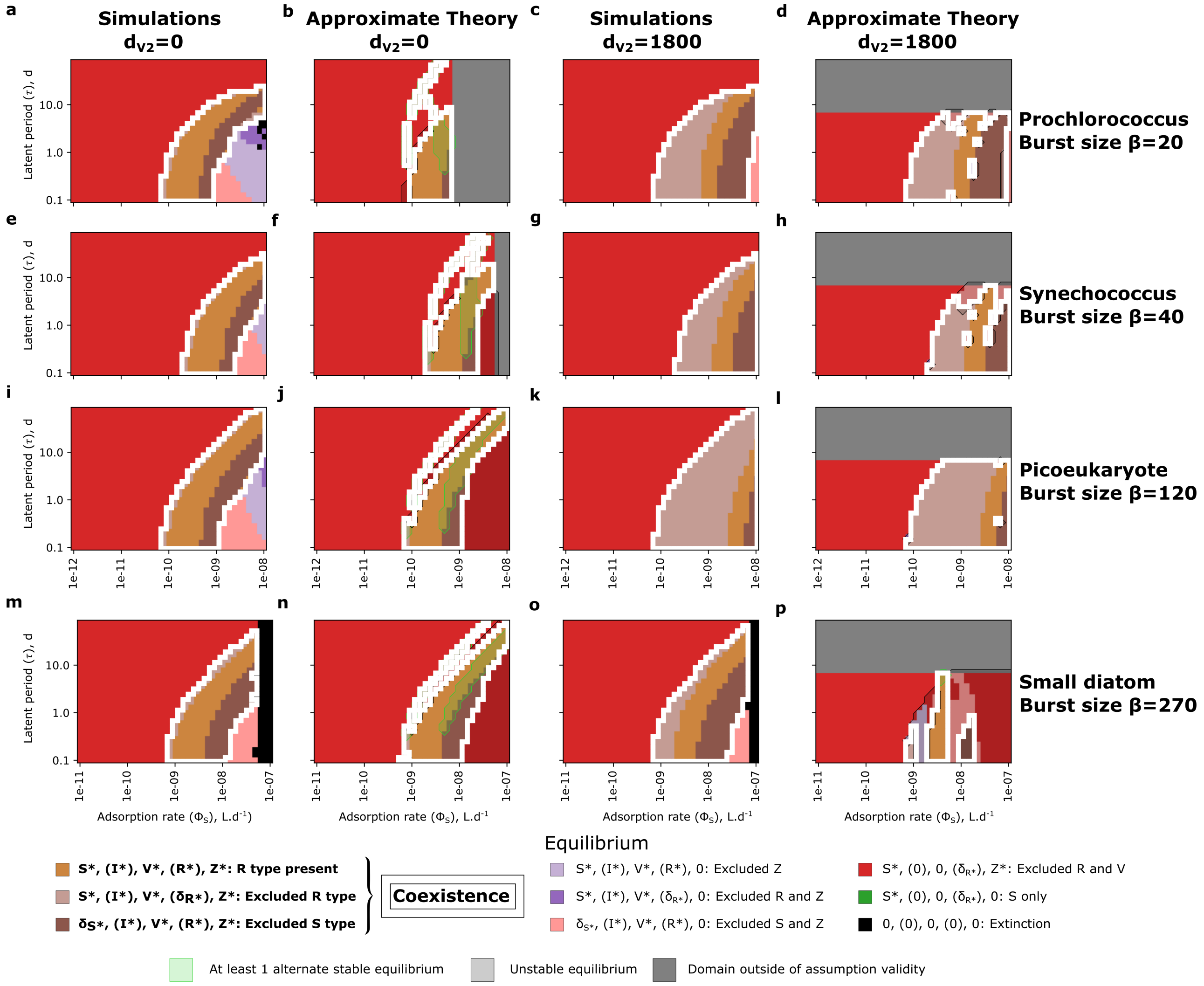


**Figure S12 | Simulated and approximate theoretical equilibrium regimes of the extracellular $SIVRZ$ model for four different phytoplankton with different molar quotas and burst sizes across the spectrum of latent period and adsorption rate of the virus.** (**a-d**) *Prochlorococcus.* (**e-h**) *Synechococcus.* (**i-l**) Picoeukaryote (non-diatom). (**m-p**) Small diatom. When multiple stable equilibria exist (shaded green area), the most complex, *i.e.* the one with the most non-null tracers, is displayed. Note the different range of adsorption rate of the virus for the small diatom. Two first left panels: quadratic mortality term present for the zooplankton only. Right panels: quadratic mortality terms present for both the zooplankton and the virus. Note that panel (**a**) and (**c**) are also shown in Figure 2. Each model was run for 20 years. For each panel, the white contour denotes the coexistence regime. For the approximate theory, when at least 1 alternate stable equilibria exist, we represent the equilibrium with the largest number of non-null tracers and highlight the zone in transparent green. Unstable equilibria areas are colored in transparent gray. The quadratic mortality terms are in $(\mu molN.L^{-1})^{-1}.d^{-1}$.

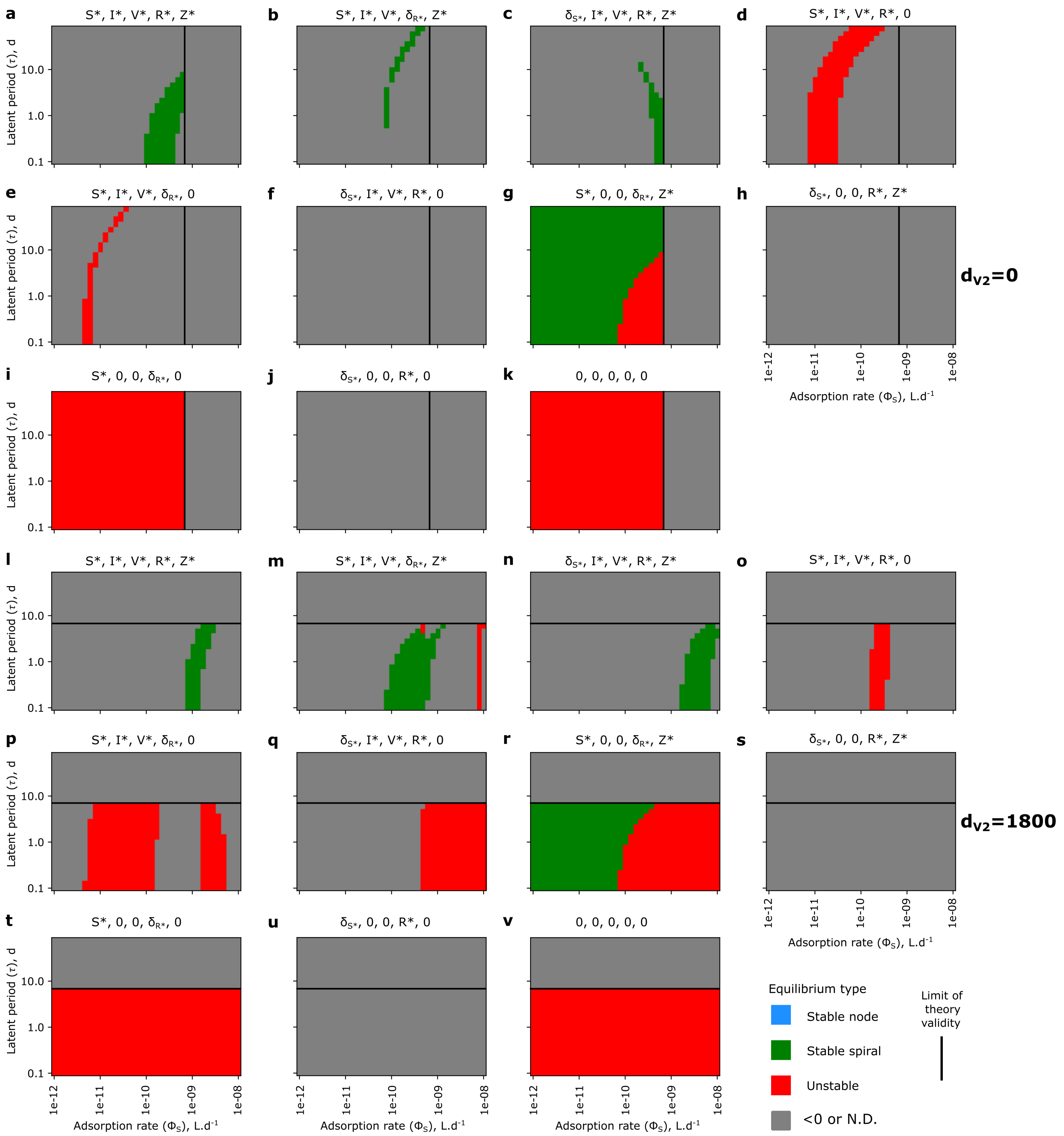


**Figure S13 | Equilibrium types for the eleven different possible equilibria of the $SIVRZ$ model in the parameter space for *Prochlorococcus*.** (**a**, **l**) $S^*, I^*, V^*, R^*, Z^*$; (**b**, **m**) $S^*, I^*, V^*, \delta_{R^*}, Z^*$; (**c**, **n**) $\delta_{S^*}, I^*, V^*, R^*, Z^*$; (**d**, **o**) $S^*, I^*, V^*, R^*, 0$; (**e**, **p**) $S^*, I^*, V^*, \delta_{R^*}, 0$; (**f**, **q**) $\delta S^*, I^*, V^*, R^*, 0$; (**g**, **r**) $S^*, 0, 0, \delta_{R^*}, Z^*$; (**h**, **s**) $\delta S^*, 0, 0, R^*, Z^*$; (**i**, **t**) $S^*, 0, 0, \delta_{R^*}, 0$; (**j**, **u**) $\delta S^*, 0, 0, R^*, 0$ and (**k**, **v**) $0, 0, 0, 0, 0$. Top panels: quadratic mortality term present for the zooplankton only. Bottom panels: quadratic mortality terms present for both the zooplankton and the virus. Regions in gray in the limit of theory validity (left and below of the black bar for top panels and bottom panels respectively) are unfeasible equilibrium: at least one of the $X^*$ is negative.

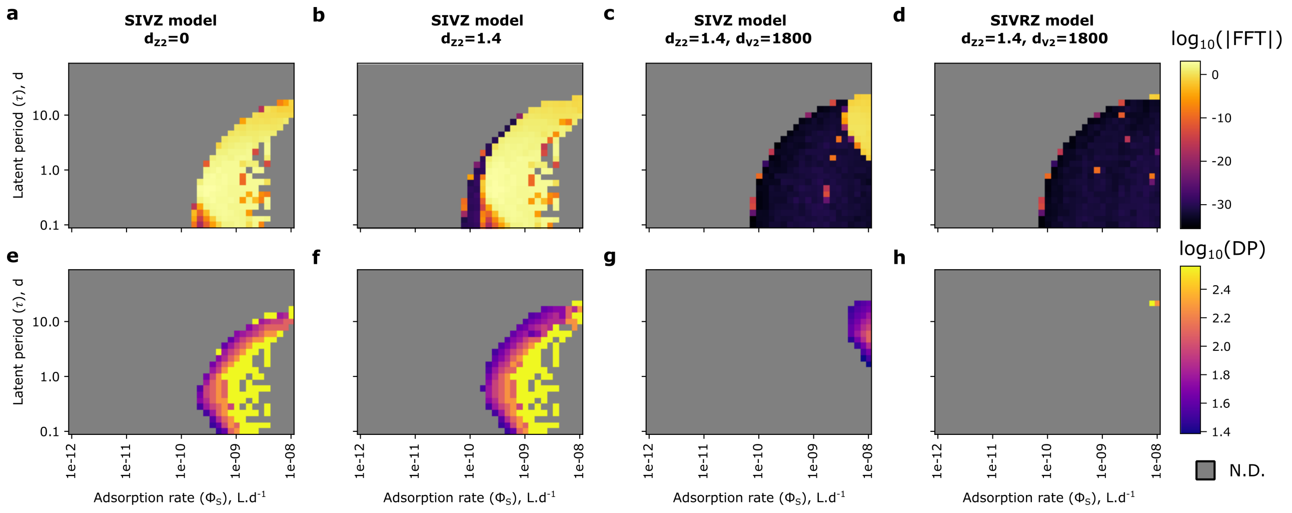


**Figure S14 | Fourier analysis of the virus time series in the last year of simulation for different parameterization of the $SIVZ$ model of *Prochlorococcus*.** (**a**, **e**) $SIVZ$ model without quadratic mortality. (**b**, **f**) $SIVZ$ model with zooplankton quadratic mortality. (**c**, **g**) $SIVZ$ model with zooplankton and virus quadratic mortality. (**d**, **h**) $SIVRZ$ model with zooplankton and virus quadratic mortality. Top panels show the modulus ($log_{10}$) of the Fast Fourier Transform (FFT). The yellow areas in the left panels show the area where the equilibrium points are unstable. Bottom panels show the Dominant Period (DP) of oscillations in $log_{10}(days)$ when the modulus of the Fourier transform is superior to 1 (arbitrary qualitative criterion). The quadratic mortality terms are in $(\mu molN.L^{-1})^{-1}.d^{-1}$. N.D.: Not Defined.

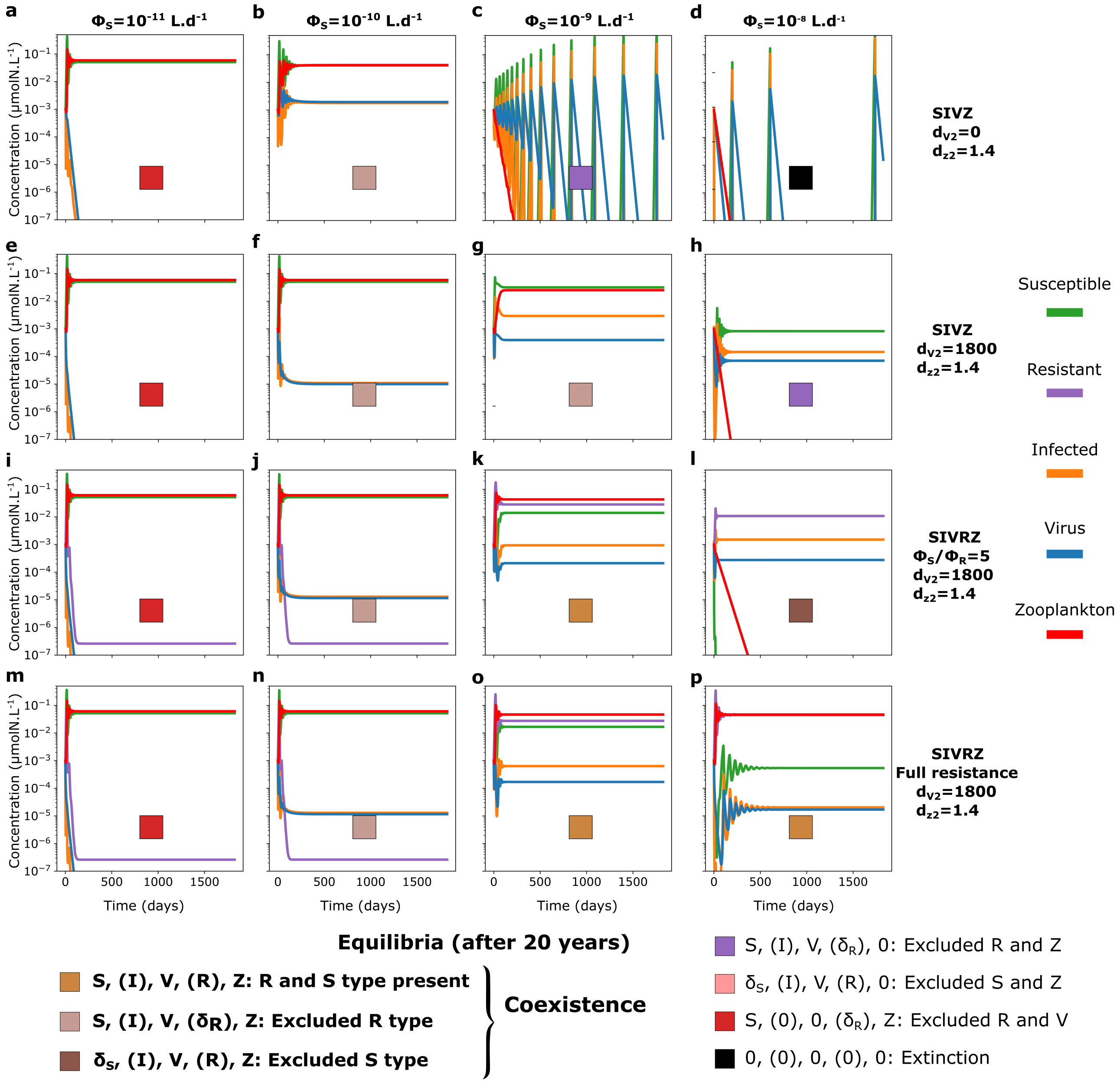


**Figure S15 | Model output of five year time series in nitrogen molar concentrations of the $SIVZ$ and the extracellular $SIVRZ$ models for *Prochlorococcus* for four different adsorption rates of the virus and different resistant type strength.** Adsorption rate from $\phi_S = 10^{-11}\ L.d^{-1}$ to $\phi_S = 10^{-8}\ L.d^{-1}$ for the $SIVZ$ model (**a-d**) without and (**e-h**) with the quadratic mortality term of the virus, for the $SIVRZ$ model (**i-l**) with a partially resistant type and (**m-p**) a fully resistant type, both with the quadratic mortality term of the virus. A latent period of 0.37 *days* and a burst size of 15 were used.

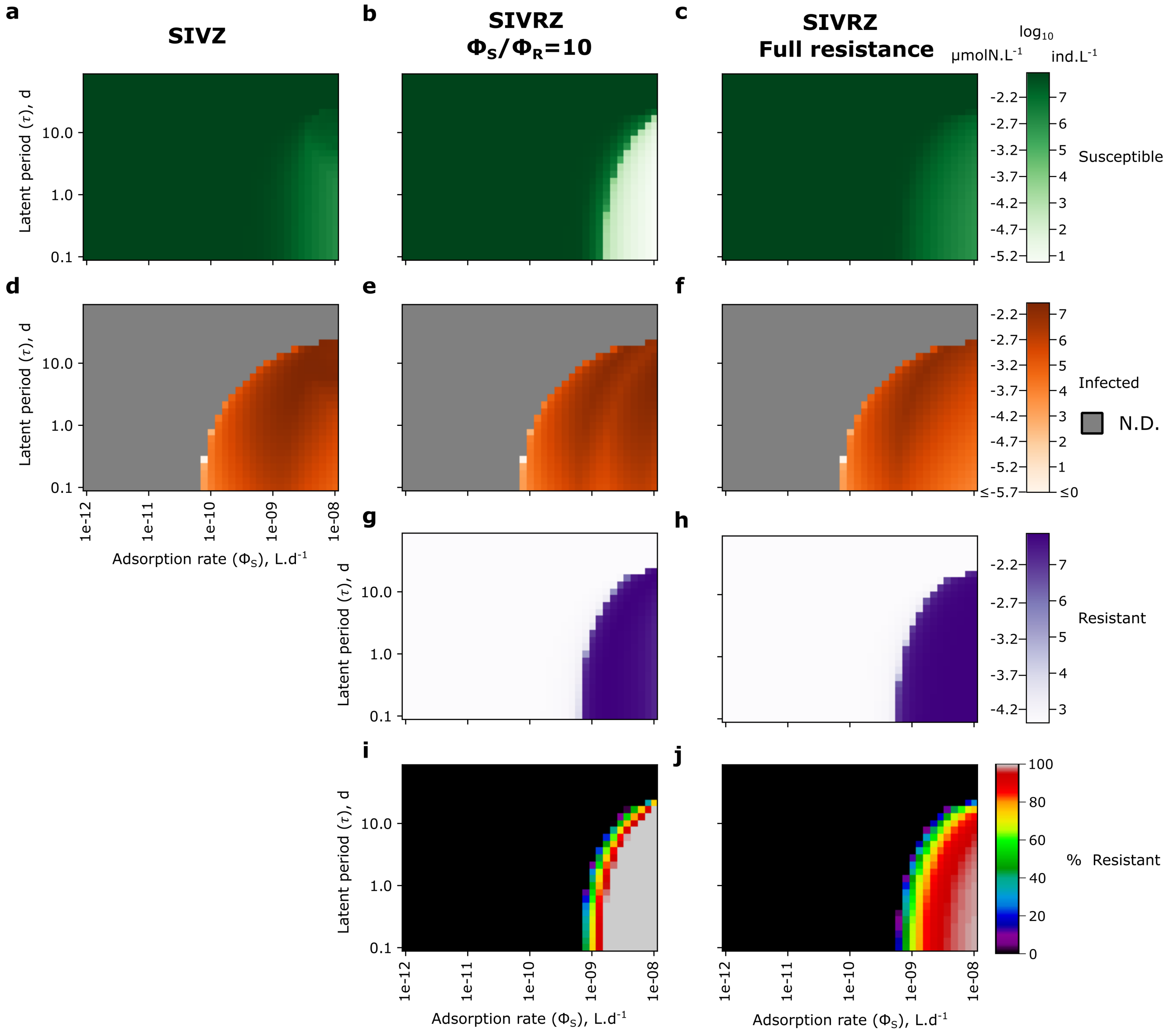


**Figure S16 | Decomposition of the concentrations of the phytoplankton in the $S$, $I$ and $R$ cell types in $SIV(R)Z$ models for Prochlorococcus at equilibrium.** Susceptible cells concentrations at equilibrium for (**a**) $SIVZ$ model, (**b**) $SIVRZ$ model with $\frac{\phi_S}{\phi_R} = 10$ and (**c**) fulll resistance of the $R$ type. (**d-f**) Infected cells concentrations. (**g**,**h**) Resistant cells concentrations. (**i**,**j**) Percentage of resistant cells (relative to $S+R$). N.D.: not defined: the virus is excluded.

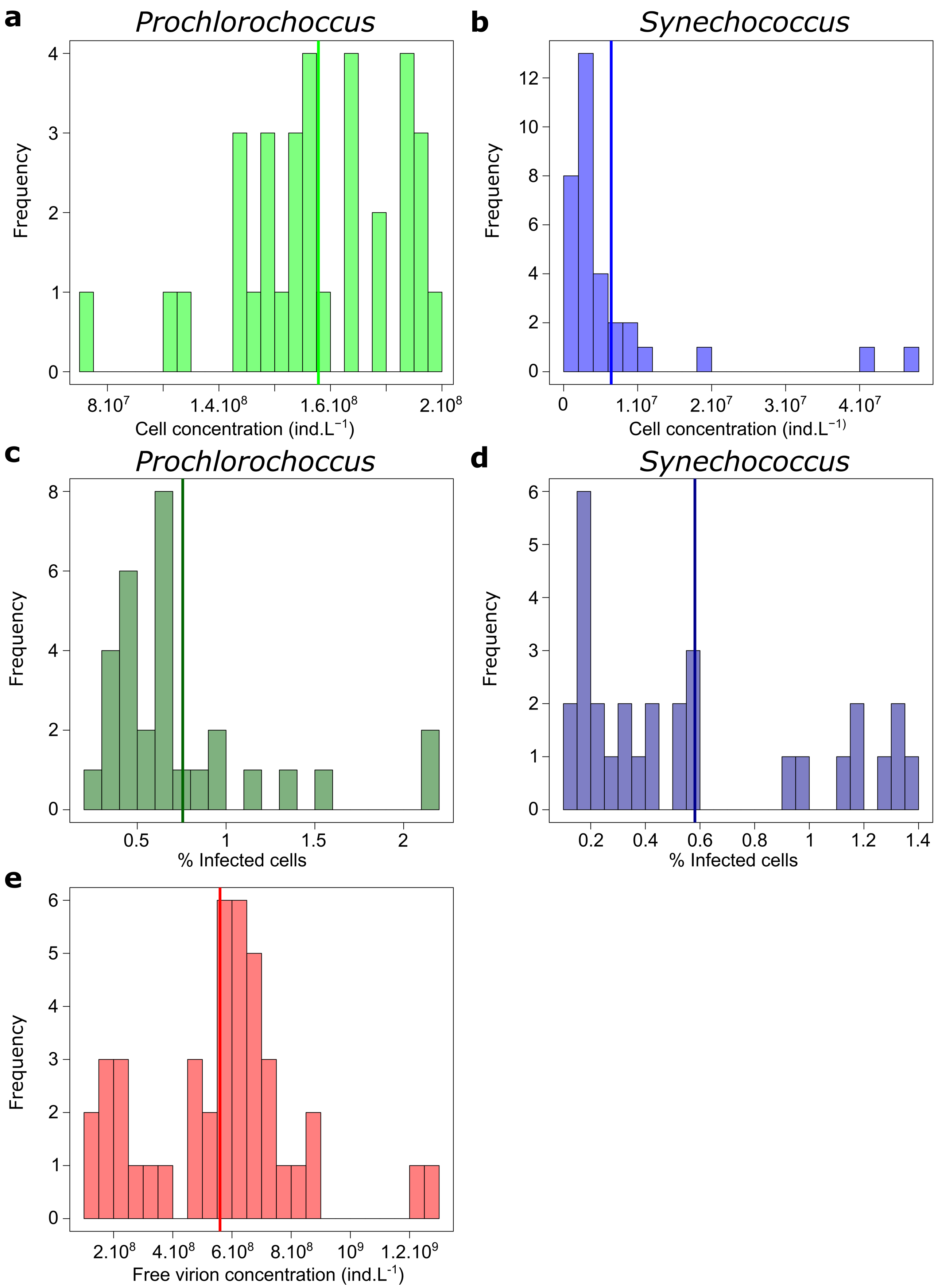


**Figure S17 | Distribution of concentration measurements of *Prochlorococcus*, *Synechococcus*, their viruses and percentages of infected cells in the North Pacific Subtropical Gyre.** Cell concentrations of (**a**) *Prochlorococcus* and (**b**) *Synechococcus*. Percentage of infected (**c**) *Prochlorococcus* and (**d**) *Synechococcus* cells. Concentrations of total free virions concentrations (T7-like plus T4-like cyanophages). All data are from Carlson et al. (2022). Mean of each distribution is represented by a vertical bar. Data is from Carlson et al. (2022) and show measurements only for latitude<33°N which is a rough delimitation of the NPSG.

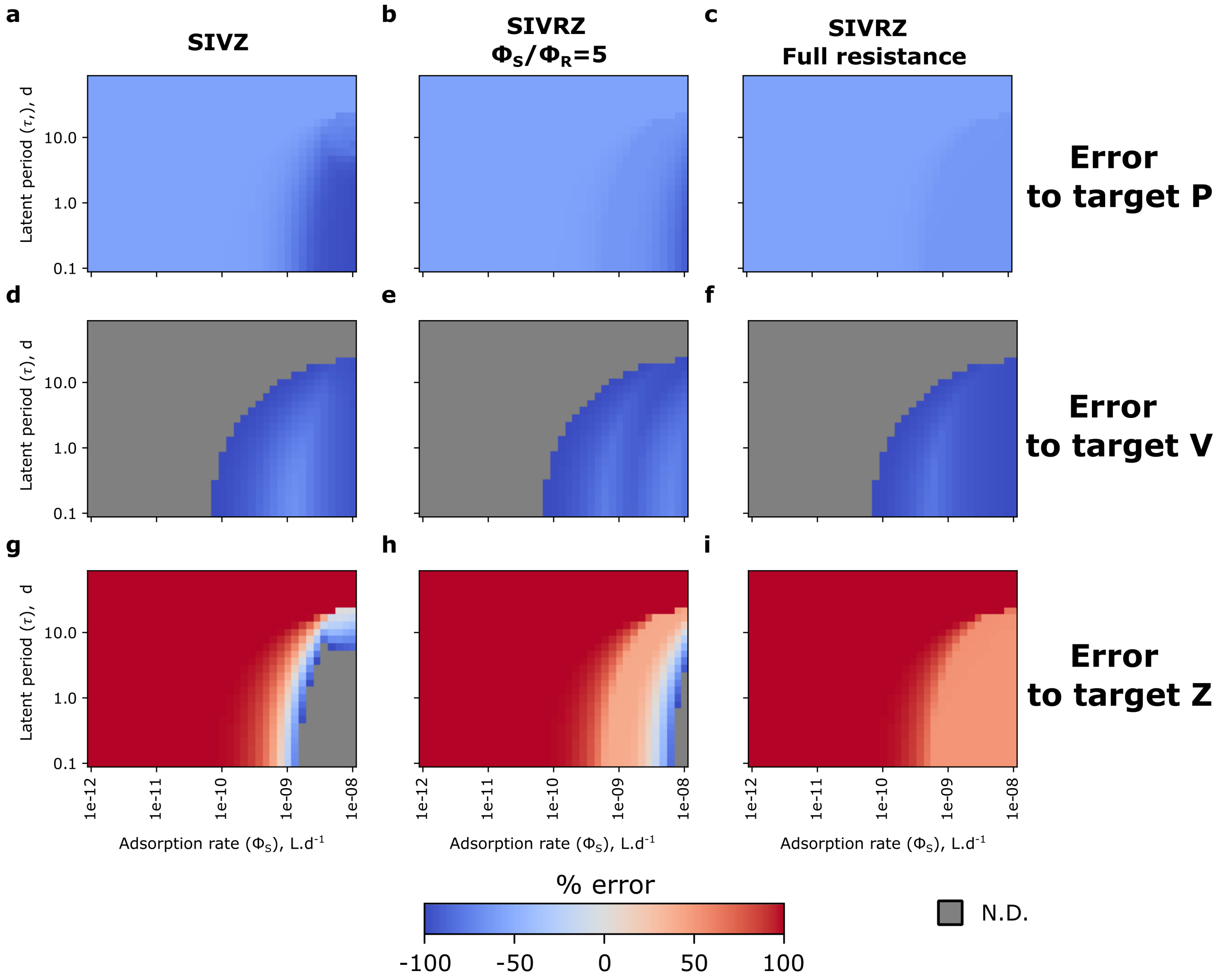


**Figure S18 | Percentage error compared to the target concentration of the different tracers for different models.** Error to target (**a-c**) total phytoplankton concentration, (**d-f**) virus concentration and (**g-i**) zooplankton concentration. N.D.: Not Defined: no coexistence.

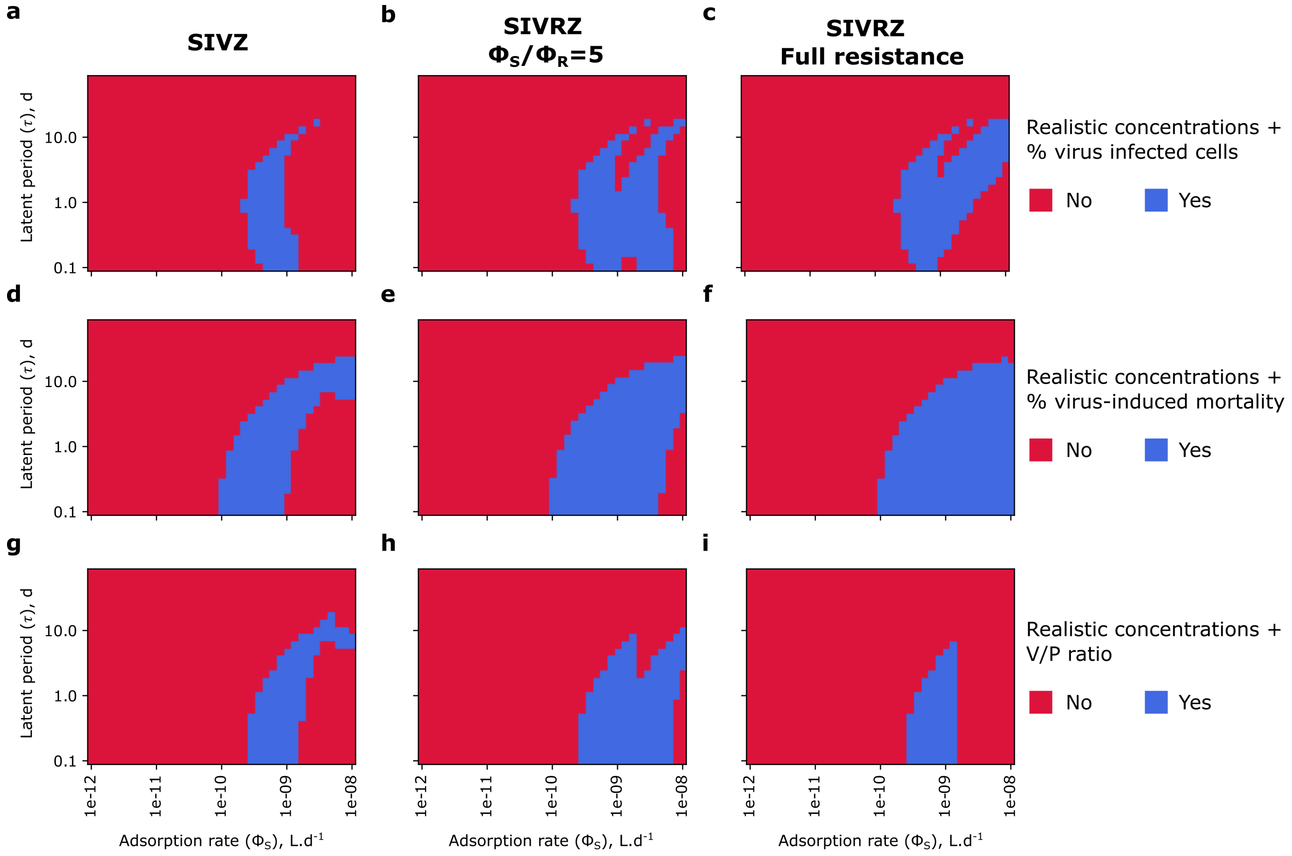


**Figure S19 | Decomposition of the conditions imposed on the models' ecology for three different models in the mesotrophic environment.** Realistic concentrations and (**a-c**) percentage of infected cells, (**d-f**) percentage of virus-induced mortality and (**g-i**) virus to phytoplankton ratio.
